# Vacuolar H^+^-ATPase subunit a was identified as the target protein of the oomycete inhibitor fluopicolide

**DOI:** 10.1101/2023.04.22.537907

**Authors:** Tan Dai, Jikun Yang, Chuang Zhao, Can Zhang, Zhiwen Wang, Qin Peng, Pengfei Liu, Jianqiang Miao, Xili Liu

## Abstract

Approximately 240 fungicides are currently in use. However, only a few direct targets have been identified, which limits the development of fungicides and rapid resistance monitoring. Fluopicolide, which is an excellent oomycete inhibitor, is classified as delocalization of spectrin-like proteins inhibitors by FRAC. In the current study, a *Pcα-actinin* knockout had no effect on the sensitivity of *Phytophthora capsici* to fluopicolide. The vacuolar H^+^-ATPase subunit a (PcVHA-a) was identified using a BSA-seq and DARTS assay. Four kinds of point mutations in PcVHA-a that cause fluopicolide resistance in *P*. *capsici* were confirmed using site-directed mutagenesis. The results of MST, molecular docking, and a DARTS assay indicated that PcVHA-a could bind fluopicolide. Sequence analysis and a molecular docking assay proved the specificity of fluopicolide to oomycetes or fish. Our results suggest that PcVHA-a is the target of fluopicolide, and H^+^-ATPase could be used as a novel target for the development of new fungicides.

## INTRODUCTION

Many oomycetes cause severe plant diseases and pose a threat to agricultural production (*1, 2*). Chemical control has the advantages of a quick effect, convenient use, freedom from regional and seasonal restrictions, and suitability for the control of a large area. Thus, chemical control plays a major role in the management of outbreaks and epidemic diseases and is the most effective method to control plant diseases at present. According to the mode of action, fungicides are divided into 11 categories, with a total of 240 compounds. Among them, the target protein of less than 150 fungicides are clear (FRAC, www.frac.info). However, the identification of target protein of the fungicide is critical for optimization and development of new fungicides and establishment of rapid resistance-monitoring method.

Many methods for studying the mode of action of a active compound include microscopy and cellular imaging (*3, 4*), affinity chromatography (*5*), isotope labeling (*6, 7*), transcriptomics (*8, 9*), and metabolomics (*10, 11*). Microscope cannot be used to clearly identify molecular targets. Isotope labeling requires tracers, but there are few stable isotopes tracers and their prices are expensive. The affinity chromatography requires the addition of chemical linker arm to the active compound, which may affect the activity of the active compound. Transcriptomics and metabolomics require the mining of high throughput data and are based on certain physiological and biochemical indicators. These target discovery methods all have certain limitations. With the improvement of target discovery methods, some novel methods such as bulked segregant analysis sequencing (BSA-seq) and drug affinity responsive target stability (DARTS), can be used to analyze the mode of action of active compound.

BSA-seq has been widely used to locate the genes related to phenotypic traits like the growth, disease resistance, and stress resistance of crops. BSA-seq was used to elucidate how PAT1 negatively regulates shoot branching in leafy *Brassica juncea*. Using BSA-seq, researchers successfully mapped the genes related to stripe rust resistance and waterlogging tolerance in wheat (*12, 13*) as well as the resistance genes against the bean mosaic virus and peanut leaf spot disease (*14, 15*). BSA-seq is an effective method to locate and highlight phenotype-related genes. General target protein point mutations can induce plant pathogens to develop resistance to fungicides; thus, BSA-seq can be used to localize resistance genes and explore target proteins in reverse.

DARTS, as a novel method, is applied to study binds of drugs and proteins to identify the target protein of drugs (*16*). DARTS plays an important role in the discovery of target proteins for small-molecule active compounds. DARTS revealed that the target protein of betulinic acid, which has anticancer activity, is the glucose regulatory protein (GRP78) (*17*), and that the target protein of alpha-ketoglutaric acid, which can prolong the life of *Caenorhabditis elegans*, is the ATP synthase β-subunit (*18*). DARTS combined with protein homology modeling revealed that the target protein of bithionol is the NAD-dependent dehydrogenase (*19*). Based on its simple principle and ease of operation, DARTS is an effective method for studying the mode of action of fungicides.

Fluopicolide (Fig. S1A) and fluopimomide (Fig. S1B) are highly effective oomycete inhibitors belonging to the benzamide group of fungicides. Fluopicolide was first registered in the UK and China in 2005. There are few reports on field resistance to this fungicide. Benzamide fungicides have been classified as delocalizing inhibitors of spectrin-like proteins (FRAC, www.frac.info). However, evidence for this classification is insufficient. Immunofluorescence tests of spectrin-like proteins showed that fluopicolide changed the location of the spectrin-like protein of *Phytophthora infestans* (*20*), but the immunofluorescence specificity and the type of spectrin-like protein in *P. infestans* are unknown. Elucidating the mode of action of fluopicolide is crucial for the targeted optimization development of this kind of fungicide and the establishment of a rapid molecular detection method of fungicide resistance.

In this study, we used *Phytophthora capsici* as the research target. We found that Pcα-actinin was not a target protein of fluopicolide through a genetic transformation assay and mutant sequencing validation. We found eight candidate proteins related to fluopicolide resistance by BSA-seq and an optimized DARTS method. Sequencing of resistant isolates and a site-mutation verification assay revealed that the key amino acid sites of fluopicolide resistance were on the vacuolar H^+^-ATPase subunit a (PcVHA-a), and were identified as G767E, N771Y, N846S, and K847R. Through molecular docking and an MST assay, we found that the target protein of fluopicolide was PcVHA-a. The above results indicate that the target protein of fluopicolide is PcVHA-a. We expect that this work will contribute to the targeting optimization development of benzamide fungicide and provide methodological support for the exploration of target proteins of drugs.

## RESULTS

### The sensitivity of Pcα-actinin knockout or overexpression to fluopicolide

There are few studies on the mode of action of fluopicolide. The scarce literature only mentions that its target protein may be associated with the skeleton protein. Taqin et al. used immunofluorescence technology to prove that fluopicolide can affect the localization of the spectrin-like protein on the plasma membrane of *P. infestans* (*20*). The spectrin-like protein superfamily is generally divided into five categories: α-actinin, α-spectrin, β-spectrin, utrophin, and dystrophin (*21*). Protein sequence blast analysis revealed that only one spectrin-like protein (506502; JGI protein ID, https://genome.jgi.doe.gov/Phyca11/Phyca11.home.html) is present in *P. capsici*. Phylogenetic analysis showed that Pc506502, α-actinin, and β-spectrin are relatively close (Fig. 1A). The sequence accession numbers are shown in Table S1. Domain analysis of Pc506502 showed that this protein belongs to the actin-binding domain and exhibits spectrin repeats and an EF-hand motif, which is typical of α-actinin (Fig. 1B). If Pcα-actinin were the target protein of fluopicolide, point mutations might be detected in resistant isolates, or the fungicide sensitivity of the gene overexpression and knockout transformants would change. The *Pcα-actinin* gene was amplified and sequenced in three pairs of sensitive and high-resistance isolates obtained in our previous study, and no point mutation in *Pcα-actinin* was detected. We obtained more than 10 independent *Pcα-actinin* knockout transformants in *P*. *capsici*, using CRISPR/Cas9 (Fig. S2). The fluopicolide sensitivity of the *Pcα-actinin* knockout transformants and GFP-fused Pcα-actinin overexpression transformants was no different from that of the wild-type isolate (Fig. 1, C and D). These results showed that Pcα-actinin is not the target protein of fluopicolide.

**Fig. 1.**
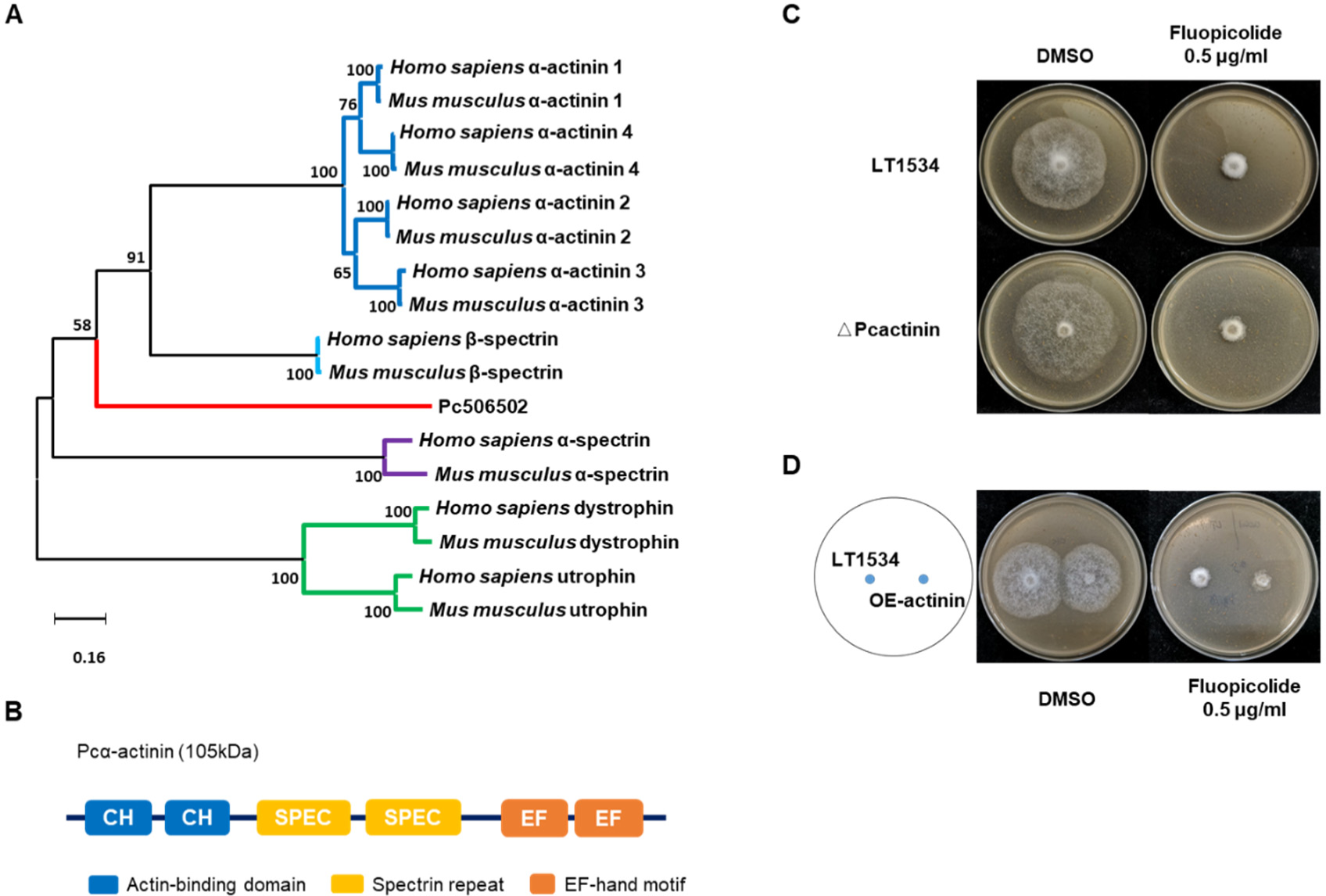
The *Pcα-actinin* knockout has no effect on fluopicolide sensitivity in *Phytophthora capsici*. (A) Phylogenetic relationships of spectrin-like proteins from representative species. The sequence accession numbers are shown in Table S1. Tree branches are colored according to the species category. The scale bar is in the unit of the number of substitutions per site. (B) Domain analysis of Pcα-actinin. (C) Fluopicolide sensitivity of the *Pcα-actinin* knockout transformant. LT1534 is the wild-type strain and △PcActinin is the knockout transformant. (D) Fluopicolide sensitivity of the *Pcα-actinin* overexpression transformant.

### Discovery of vacuolar H^+^-ATPase subunit a by the combination of BSA-seq and DARTS

BSA-seq and DARTS were used to explore the target protein of fluopicolide. For BSA-seq, the first step was to obtain the offspring of character separation. In this study, the sensitive wild-type strain BYA5 and fluopicolide-resistant strain LT-10 (resistance factor > 1000) were hybridized to obtain 71 F1 generations. Two F1-generation resistant isolates were selected for sister crossing to obtain the F2 generations. Among them, the F1 generations were all resistant, whereas the F2 generations contained both sensitive and resistant isolates. Fifty-one resistant isolates (resistance factor > 1000) and 32 sensitive isolates were selected from the F2 generations to construct a resistant mixed DNA pool and a sensitive mixed DNA pool, respectively (Fig. 2A). Reliable single nucleotide polymorphism (SNP) data were obtained through sample genome sequencing and SNP filtering. The SNP-index and ED algorithm were used to perform association analysis on the sequenced clean data and obtain the associated scaffolds (Tables S2 and S3) and SNPs. After functional annotation of the associated SNPs, a total of 284 non-synonymous mutations were obtained (Table S4). The BSA-seq candidate genes were verified by performing sequence alignment analysis of both sensitive and resistant isolates, and no resistance genes were found. Nonetheless, we found that resistance-related genes could be located on scaffold 12 and scaffold 14 (Figs. 2B and S3).

**Fig. 2.**
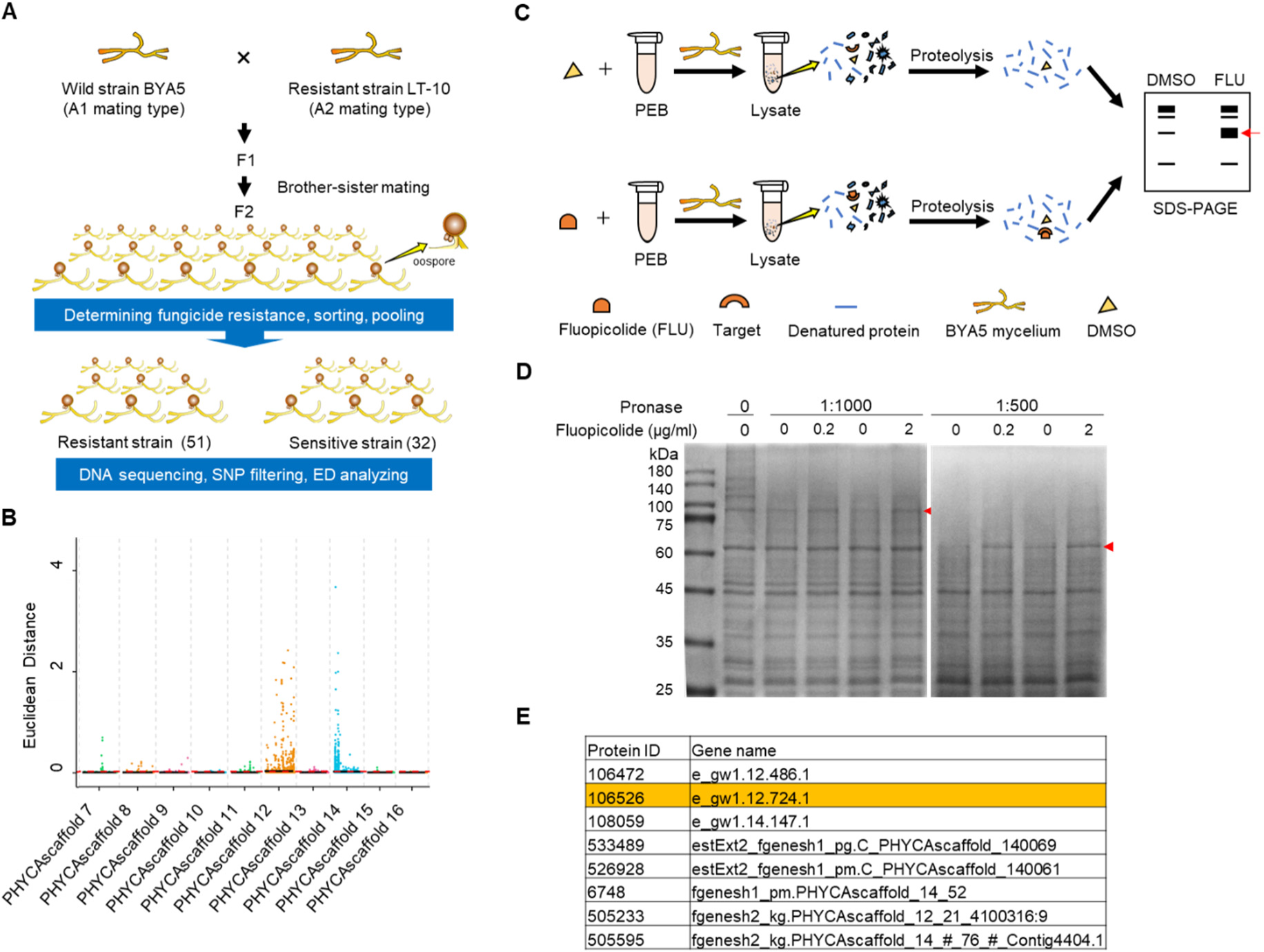
Combination of BSA-seq and optimized DARTS to explore target proteins of fluopicolide. (A) Schematic diagram of the BSA-seq assay. F2 generations were produced by the brother-sister mating of the F1 generation, which was obtained by crossing two parent strains with distinct backgrounds and which were vastly different in fungicide sensitivity. Fifty-one resistant and 32 sensitive F2 generation isolates were used to obtain DNA mixed pooled samples for sequencing and data analysis. (B) Distribution of ED-based linkage values on scaffolds. A larger ED value indicates a stronger linkage of the SNP site to the resistant trait. (C) Scheme for the methodology of optimized DARTS. Fluopicolide or DMSO was added to the protein extract to obtain the BYA5 protein, followed by proteolysis and SDS-PAGE detection. The most notable protein bands were excised for mass spectrometry detection. (D) Target identification for potential fluopicolide-interacting proteins by DARTS. The BYA5 membrane protein lysates were incubated with or without fluopicolide. After protein electrophoresis and Coomassie bright blue staining, two specific protein bands (indicated by arrows) were identified. (E) Candidate proteins were obtained. Only eight candidate proteins were found in the intersection between the proteins obtained by mass spectrometry and the candidate regions of BSA-seq.

Subsequently, we optimized the existing DARTS method (Fig. 2C) to confirm the fluopicolide resistance proteins. In this experiment, the concentration of pronase was set at 1:500 and 1:1000 (m/m). At the concentration of 1:1000, the abundance of the protein bands at 100 kDa in the treatment group was greater than that in the control group. At the concentration of 1:500, there was also a difference in the abundance at 60 kDa (Fig. 2D). The target protein bands were detected by mass spectrometry to obtain a series of candidate proteins (Supplementary Materials 1 and 2), and only less ten andidate proteins were located on scaffold 12 and scaffold 14 (Fig. 2E).

Seventeen resistant mutants and five sensitive isolates were used to sequence the encoding genes of the eight candidate proteins. Protein Pc106526 had point mutation sites in all resistant isolates and was annotated as vacuolar H^+^-ATPase subunit a (VHA-a). The G767E mutation site was found in strains J-7, J-10, J-12, 11-1, 11-2, 11-3, 11-5, 11-6, and 11-7. The N771Y mutation site was found in strains L-2 and L-3. The N846S mutation site was found in strains B-1, B-12, and J-8. In addition, the K847R mutation site was found in strains LT-7, LT-8, and LT-10 (Fig. 3A). Among them, the N846S mutation site was only found in low-resistance isolates, whereas the others were found in high-resistance mutants, and they were all heterozygous except for B-1 and B-12 (Fig. S4). The results showed that the point mutation of PcVHA-a might induce the resistance of *P. capsici* to fluopicolide.

**Fig. 3.**
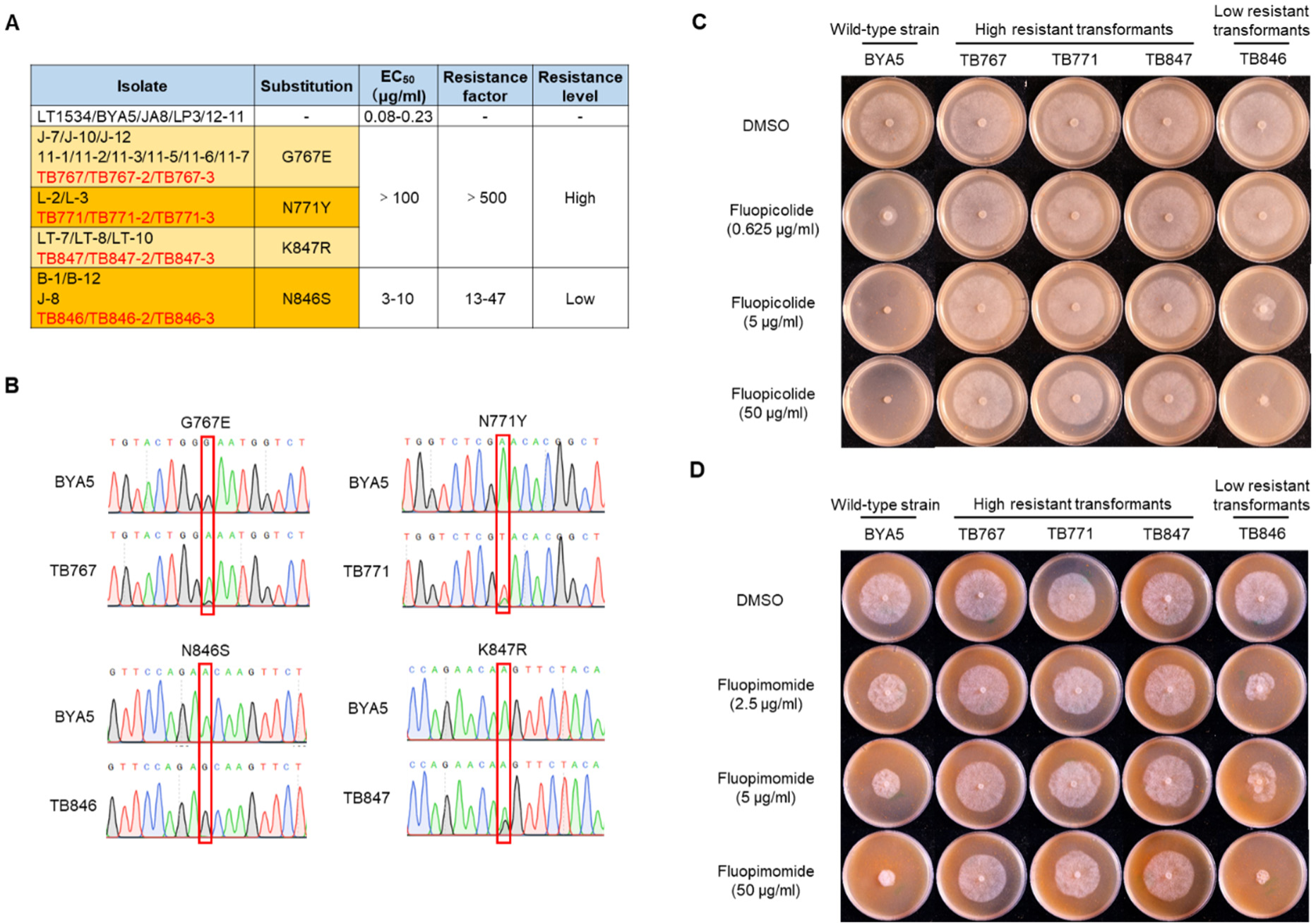
Four kinds of point mutations in PcVHA-a can cause fluopicolide resistance in *Phytophthora capsici*. (A) Fluopicolide sensitivity of mutants obtained by fungicide adaption and site-directed edited transformants. EC_50_, concentration showing 50% inhibition of mycelial growth; resistance factor, the ratio of EC_50_ of fluopicolide for mutants or transformants relative to the that of the parental isolate. (B) Sequence alignment of the VHA-a from transformants and parental isolates. (C) Mycelial growth of *P*. *capsici* site-directed transformants on the fluopicolide-amended V8 medium. (D) Mycelial growth of *P*. *capsici* site-directed and edited transformants on the fluopimomide-amended V8 medium.

### Confirmation of the relationship between the point mutation in PcVHA-a and fluopicolide resistance

Site-directed mutations in the *PcVHA-a* gene from BYA5 were conducted using CRISPR/Cas9. Finally, 12 positive transformants containing G767E, N771Y, N846S, or K847R in PcVHA-a were successfully obtained (Fig. 3A). Among them, TB767, TB767-2, TB767-3, TB771, TB771-2, TB771-3, TB847, TB847-2, and TB847-3 were heterozygous transformants, and TB846, TB846-2, and TB846-3 were homozygous transformants (Fig. 3B). The EC_50_ of the point mutation transformants containing the G767E, N771Y, and K847R sites were all greater than 100 and the resistance factors were greater than 500, and were the same as the sensitivity of the corresponding resistant mutants obtained by fungicide adaption (Fig. 3, A and C). The EC_50_ values of the N846S transformants TB846, TB846-2, and TB846-3 were 4.34, 9.57, and 2.99 μg/ml, respectively, which was the same level of resistance as that of the low-resistance mutants B-1, B-12, and J-8 (EC_50_ of 3.90, 8.31, and 5.44 μg/mL, respectively) (Fig. 3, A and C). The EC_50_ values of the mutant isolates (*22*) and transformants are shown in Table S5.

Fluopicolide and fluopimomide are benzamide fungicides and exhibit positive cross-resistance (*23*). Hence, they may have the same mode of action. Therefore, the fluopimomide sensitivity of the transformants produced by site-directed mutations was also determined. The results showed that the point mutations of G767E, N771Y, and K847R could cause high resistance of *P. capsici* to fluopimomide, whereas the fluopimomide sensitivity of the N846S transformants was not significantly different from that of wild-type isolates (Fig. 3D). The results showed that the resistance of fluopimomide was also caused by the point mutation of PcVHA-a, and the same resistance molecular mechanism applies to both fluopicolide and fluopimomide. Nevertheless, the sites related to fluopicolide and fluopimomide resistance are not exactly the same.

### Identification of the binding ability of fluopicolide and PcVHA-a

Three methods (DARTS + Western blot, MST, and molecular docking) were used to verify the binding ability of fluopicolide and PcVHA-a. First, PcVHA-a (labeled with 3× FLAG) overexpression transformants were obtained using PEG/CaCl_2_-mediated protoplast transformation. The optimized DARTS method and Western blot were used to detect the binding of PcVHA-a and fluopicolide, in which the mass ratio of total protein of the PcVHA-a overexpression transformants to pronase was 1:500 (m/m) and the concentration of fluopicolide was 20 and 200 μg/ml. The band abundance of PcVHA-a was substantially higher than that of the treatment without fluopicolide. Under the same conditions, the band abundance of endogenous β-tubulin did not substantially change compared with the treatment without fluopicolide (Fig. 4A). The results showed that fluopicolide could specifically bind to PcVHA-a.

**Fig. 4.**
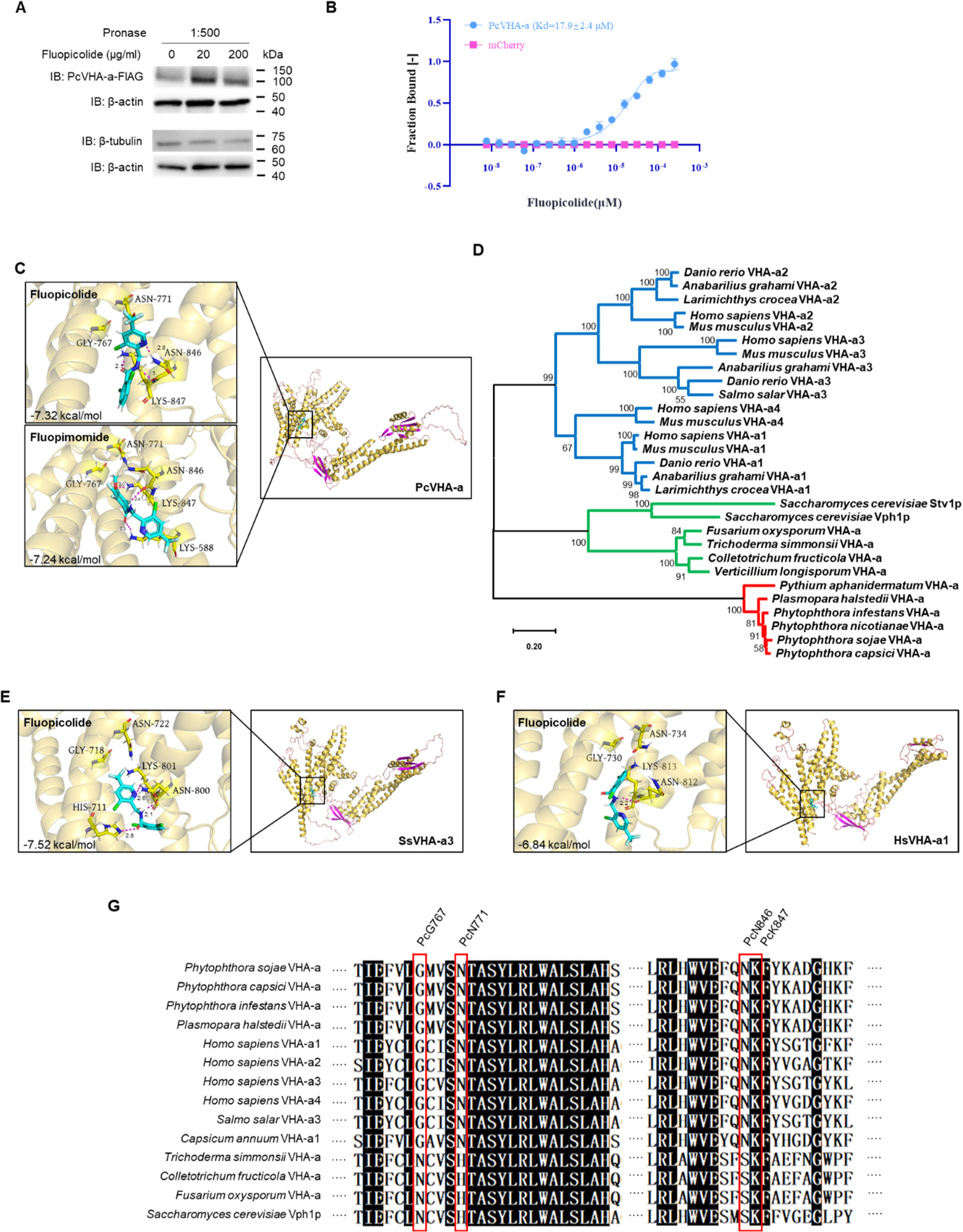
Verification of binding of fluopicolide and PcVHA-a. (A) DARTS confirmed that fluopicolide binds specifically to VHA-a. VHA-a and α-actinin were fused with 3× FLAG, and β-tubulin was the endogenous protein in *P. capsici*. The three proteins were treated with pronase (1:500, m/m) for 15 min. β-actin as an internal reference marker was not treated with pronase. (B) MST assays revealed the binding ability of fluopicolide and PcVHA-a; FNorm means normalized fluorescence. (C) Docking model for fluopicolide or fluopimomide and PcVHA-a; the enlarged view of the VHA-a binding site shows the interactions of the H-bonds (distances are indicated in magenta). (D) Phylogenetic relationships of VHA-a from different representative species; the sequence accession numbers are shown in Table S6. Tree branches are colored according to the species category. The scale bar is in the unit of the number of substitutions per site. (E) Docking model of fluopicolide and SsPcVHA-a3. (F) Docking model of fluopicolide and HsVHA-a1. The fungicides were docked into the potential binding pocket containing 767, 771, 846, and 847 amino acid residues of VHA-a. (G) Sequence alignment of amino acid residues of oomycetes, plants fungus, humans, and fish (767, 771, 846, and 847, respectively); the four key amino acid residues of humans, fish, and oomycetes were the same, while the corresponding site of PcN846 in fungi was the natural resistance site for serine acid.

To quantitatively characterize the affinity of the intermolecular interactions between PcVHA-a and fluopicolide, the binding dissociation constant (Kd) was measured by MST with the protein lysates of PcVHA-a-mCherry overexpression *P. capsici* transformants. As expected, PcVHA-a-mCherry was readily bound by fluopicolide, whereas the mCherry-only control showed no detectable interaction with fluopicolide (Fig. 4B). Moreover, the Kd value of fluopicolide/PcVHA-a-mCherry was calculated as 17.9 μM by MST analysis, indicating a strong interaction between fluopicolide and PcVHA-a.

To further explore the key amino acid sites for the interactions between fluopicolide and PcVHA-a, molecular-docking studies were carried out. A Ramachandran plot of the yielded 3D model showed that 96.8%, 97.6%, and 97.5% of residues were in favored regions (98%), and 99.2%, 99.4%, and 99.9% of residues were in allowed regions (>99.8%), suggesting the stereochemical rationality of the 3D models of PcVHA-a (Fig. S5), SsVHA-a1 (Fig. S6), and HsVHA-a3 (Fig. S7). N846 and K847 were predicted to promote the binding of fluopicolide-PcVHA-a, in which fluopicolide formed three hydrogen bonds (H-bond) with N846 and K847 with a binding force score of −7.32 kcal/mol (Fig. 4C, fluopicolide). The binding modes of fluopicolide and fluopimomide to PcVHA-a were similar. The nitrogen atoms (N) in their acylamino groups both formed an H-bond with the oxygen atom (O) of the N846 backbone. Moreover, in fluopimomide docking (binding force score = −7.24 kcal/mol (Fig. 4C, fluopimomide)), an H-bond was detected between the carbonyl O in the acylamino groups of the two small molecules and the N atom of the K588 sidechain. The binding force score suggested the higher binding affinity of fluopicolide to PcVHA-a. To determine the effect of the four amino acids on the binding force of PcVHA-a and fluopicolide, amino acid substitutions were introduced into each residue: G767E, N771Y, N846S, and K847R. The H-bonds of these key residues disappeared, and their binding force scores decreased to 5.45, 5.53, 5.93, and 6.39 kcal/mol (Fig. S8), respectively. Fluopicolide inhibited the mycelial growth of BYA5 and OE-VHA-a, but not that of OE-VHA-a^G767A^ (Fig. S9). These results indicate that fluopicolide and fluopimomide bind with PcVHA-a to inhibit the function of V-ATPase, and the substitutions of key amino acid residues reduce the binding force of the benzamide fungicides and PcVHA-a.

Fluopicolide is a fungicide with low toxicity to mammals, plants, and fungi, but high toxicity to fish (*24*). According to the protein sequence blast in the NCBI database (https://www.ncbi.nlm.nih.gov/), there are generally four types of VHA-a in mammals: a1, a2, a3, and a4. Among them, a1 has multiple redundant proteins in mammals, but few redundant proteins in fish and plants (*Salmo salar* has only VHA-a3). Oomycetes and most fungi have one VHA-a, (Table S6). Bioinformatics analysis of the VHA-a of different species revealed that fungi, oomycetes, and mammals could be clustered into separate groups, and oomycetes are more distantly related to mammals (Fig. 4D). The sequence accession numbers are shown in Table S7. So, why is fluopicolide low-toxic to humans and fungi but highly toxic to fish and oomycetes? To answer this question, molecular-docking analyses were performed using *S. salar* VHA-a3 (SsVHA-a3) and *Homo sapiens* VHA-a1 (HsVHA-a1). The binding modes of SsVHA-a3 and PcVHA-a to fluopicolide were similar. The nitrogen atoms (N) in the acylamino and pyridine groups formed H-bonds with the oxygen atom (O) and nitrogen atoms (N) of the N846 backbone (Fig. 4E). Compared with HsVHA-a1 docking, the key H-bonds disappeared and the binding force score decreased (Fig. 4F), which was consistent with the low toxicity of fluopicolide observed in humans and plants and the high toxicity observed in fish. Analysis of the key amino acid sites of fluopicolide showed that the fungal amino acids corresponding to the G767, N771, and N846 of *P. capsici* were asparagine, histidine, and serine, respectively, resulting in natural resistance (Fig. 4G).

### The effect of fluopicolide on the localization of Pcα-actinin and PcVHA-a in *Phytophthora capsici*

Fluopicolide changed the location of Pcα-actinin in the plasma membrane of *P. capsici*. However, Pcα-actinin was not the target protein of fluopicolide. So, how did its location change? mCherry and GFP were used to label PcVHA-a and Pcα-actinin to show the location of these two proteins (Fig. 5A). In the Pcα-actinin-mCherry overexpression transformants, Pcα-actinin was mainly located on the cell membrane. After treatment with fluopicolide, the localization of Pcα-actinin changed to a point distribution in the cytoplasm. In the PcVHA-a-GFP overexpression transformants, PcVHA-a was located on the large and small vesicular-like organelles, and no substantial change in the location was observed after treatment with fluopicolide. We also obtained the positive co-expression transformants of Pcα-actinin and PcVHA-a (Fig. 5B) and the two proteins were correctly expressed in *P. capsici* (Fig. 5C). Pcα-actinin and PcVHA-a were co-localized, and this co-localization did not change regardless of fluopicolide treatment. Pcα-actinin exhibited a small, vesicular-like localization under fluopicolide treatment, which was close to the characteristic location of PcVHA-a. These two proteins exhibited substantial co-localization when co-expressed, indicating that the change of location of Pcα-actinin may be affected by PcVHA-a.

**Fig. 5.**
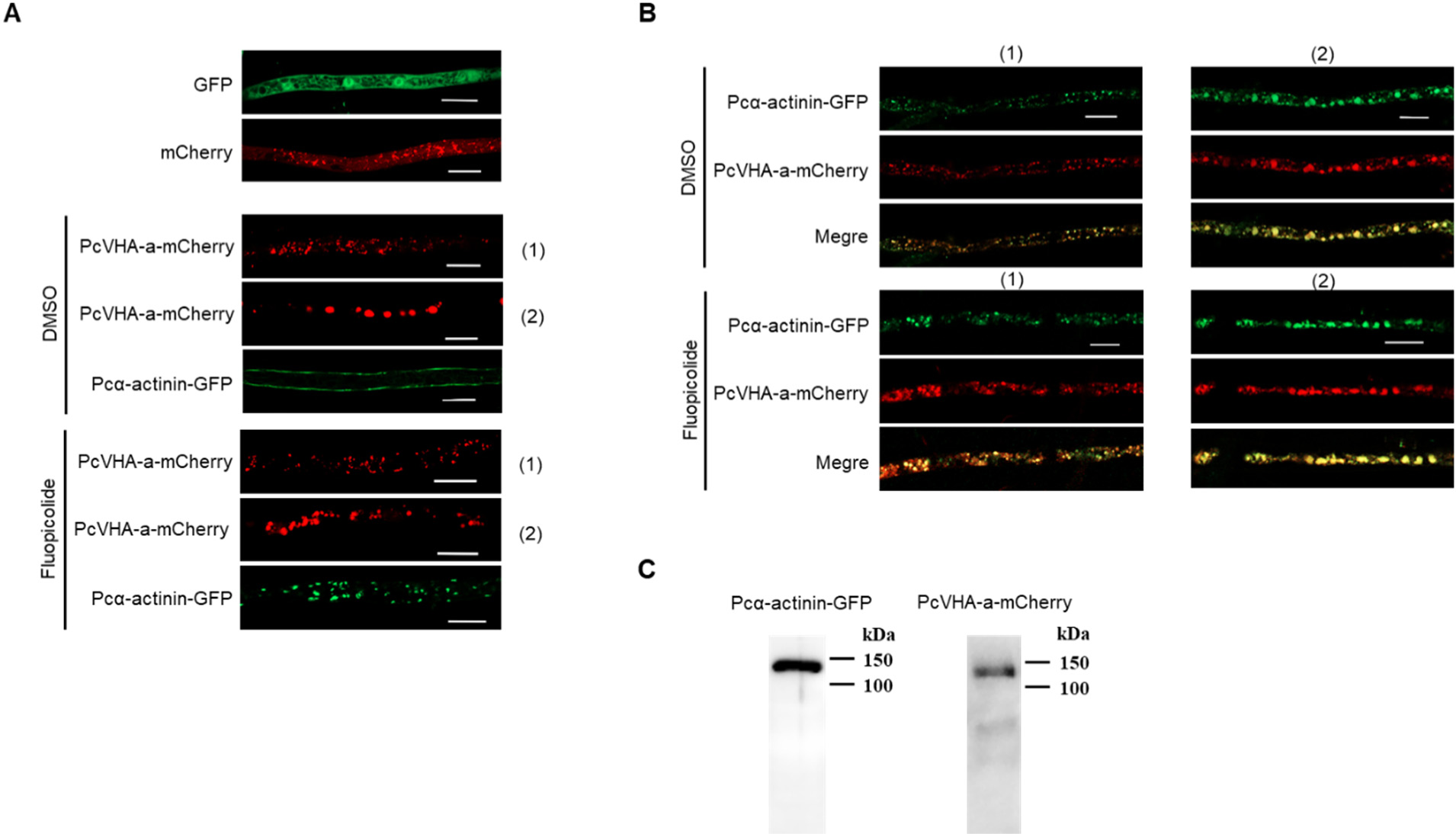
The effect of fluopicolide on the localization of PcVHA-a and Pcα-actinin. (A) The hyphae of transformants expressing only GFP and mCherry. (B) The hyphae of transformants expressing Pcα-actinin-GFP and PcVHA-a-mCherry, with or without 10 μg/mL fluopicolide treatment for 30 min. PcVHA-a-mCherry was located on the small (1) and large (2) vesicular-like organelles. (C) The hyphae of transformants co-expressed Pcα-actinin-GFP and PcVHA-a-mCherry, with or without 10 μg/mL fluopicolide treatment for 30 min. PcVHA-a-mCherry and Pcα-actinin-GFP were located on the small (1) and large (2) vesicular organelles. (D) Western blot analysis of Pcα-actinin-GFP and PcVHA-a-mCherry in the co-expression transformants; the bars represent 10 μm.

## DISCUSSION

Only a few studies on the mode of action of fluopicolide are available. Based on an immunofluorescence assay of spectrin-like proteins in *P. infestans* (*20*), FRAC (fungicide resistance action committee) classified fluopicolide as a delocalization of spectrin-like proteins inhibitor. However, the specificity of the antibodies (S1390) used in this assay was controversial. Marta et al. identified anti αβ-spectrin immunoreacting peptides as the elongation factor 2 and Hsp70 from *Neurospora crassa* and *P. infestans* (*25*). The high-level resistance of phytopathogens to fungicides is mainly caused by point mutation of the target protein (*26-30*). We conducted phylogenetic analysis on spectrin-like proteins and identified the spectrin-like protein Pc506502 in *P. capsici*. We knocked out and over-expressed Pcα-actinin to obtain the corresponding transformants. The sensitivity of the transformants to fluopicolide did not change, and no point mutation of the protein was detected in the resistant strain, which revealed that Pcα-actinin was not the target protein of fluopicolide.

BSA-seq is an efficient method to rapidly and efficiently locate genes of extreme phenotypes. BSA-seq can directly locate the target protein or locate the target protein in a smaller candidate region by combining it with other methods. The target protein candidates can thus be greatly reduced and the target protein can be identified. Nonetheless, BSA-seq exhibits some shortcomings. For instance, locating the target protein causes resistance based on the point mutation, which is required to be able to construct the resistant and sensitive pools for sequencing analysis. In addition, BSA-seq requires a high level of genome assembly. In this study, no resistance genes were detected in the 284 candidates obtained by BSA-seq. This might be explained by the genome of *P. capsici* not having been assembled to the chromosome level because BSA-seq can only locate the target gene in a large candidate region.

Furthermore, we used optimized DARTS to increase the chance of identifying the target protein. The main application of DARTS optimization is the incubation of drugs and proteins. At present, there are two main incubation methods. One is the direct incubation of cell lysates and drugs (*19, 31*), where the target protein may be partially degraded during protein extraction. The other is the incubation of drugs and cells *in vivo* (*17*), which has the risk of producing non-specific false-positive proteins because of the effect of drugs on the cell proteome. Therefore, we added fluopicolide to the protein extract to increase the stability of the protein during both extraction and protease treatment. Furthermore, we divided the proteins into total and membrane proteins for extraction, which further amplified the differences of the final SDS-PAGE protein banding (Fig. 2, C and D). Only eight candidate proteins obtained by DARTS were localized in scaffolds 12 and 14 obtained by BSA-seq, of which protein 106526 exhibited a point-mutation site in the resistant strain. Hence, BSA-seq and DARTS methods complement each other and can accelerate the exploration of target proteins.

Three methods were used to verify the target protein of fluopicolide, and the results revealed that fluopicolide could specifically bind PcVHA-a. The DARTS method not only provided candidate target proteins but also verified the specificity of the binding of a single candidate protein to the small molecules. MST has been widely used to determine the binding affinities of small molecules and fluorescence-labeled target proteins (*32-34*). We obtained the binding dissociation constant of fluopicolide and PcVHA-a by the MST assay and determined the binding force between the target protein and the drug.

AlphaFold2 (AF2), developed by DeepMind, can regularly predict protein structures with atomic accuracy (*35*), making it the most widely used algorithm today. We used AF2 to predict the structure of PcVHA-a and found that G767, N771, N846, and K847 had close spatial locations, which were potential fungicide-binding sites. The molecular docking and fluopicolide sensitivity of OE-VHA-a^G767A^-overexpressing transformants revealed that fluopicolide and fluopimomide bind to PcVHA-a and inhibit the V-ATPase function. Differences in the binding modes of PcVHA-a of the two fungicides were observed. N846 and K847 formed key hydrogen bonds with the fungicides, but G767 and N771 did not. G767 and N771 might have affected the binding of the fungicides with PcVHA-a in the spatial structure. Point mutations in these four residues are required for *P. capsici* to develop resistance.

A molecular-docking assay and bioinformatics analysis showed that differences in the 3D structures of VHA-a in fish and humans as well as the existence of natural resistance sites in fungi resulted in the high toxicity of fluopicolide to fish and oomycetes and its low toxicity to fungi and humans. This limits the use of benzamide fungicides in aquatic crops and for the treatment of diseases caused by pathogenic fungi. Nevertheless, based on the structure of the VHA-a of different species, we could design agents with low toxicity to humans and fish and high toxicity to fungi and oomycetes in the future.

We found that the target protein of fluopicolide was PcVHA-a. So, why would it affect the localization of Pcα-actinin? We found that fluopicolide treatment could indeed cause the membrane localization of Pcα-actinin to almost disappear, but when co-expressing Pcα-actinin and PcVHA-a, both exhibited obvious small vesicular-like co-localization with the same characteristics as the localization of Pcα-actinin alone after fluopicolide treatment. We speculated that the change in location of Pcα-actinin caused by fluopicolide treatment was mediated by the involvement of PcVHA-a, but the specific mechanism needs further study.

PcVHA-a is one of the subunits of vacuolar H^+^-ATPase (V-ATPase), which is an ATP-dependent proton pump that can pump cytoplasmic H^+^ into the intracellular compartments (*36*). V-ATPase is widely distributed in the cell membrane system of eukaryotic cells, including the endosome, lysosomes, Golgi vesicles, secretory vesicles, and cell membranes (*37, 38*). V-ATPase plays a crucial role in the invasion and migration of tumor cells and contributes to the drug resistance of tumor cells (*39, 40*). Thus, VHA-a is the key target protein of new strategies for cancer treatment (*41-43*). In recent years, research on new drugs produced with V-ATPase as the target has attracted increased attention (*44-46*). V-ATPase contains at least 13 subunits among different species, and it is uncertain which subunit is more likely to be the target. Our research indicates that PcVHA-a is a target protein of fluopicolide. However, the mechanism underlying how fluopicolide affects the biological function of V-ATPase by binding to PcVHA-a needs further study.

In conclusion, we used three binding ability verification tools to validate the fluopicolide target proteins identified by DARTS and BSA-seq and found that four-point mutations on PcVHA-a could lead to the resistance of *P. capsici* to fluopicolide. The high toxicity to the target organism and low toxicity to the non-target organisms suggest that VHA-a will become an important drug target in other fields. In addition, a unique combination of BSA-seq and DARTS was used in this study to quickly determine the target protein of the fungicides, providing a crucial means for the exploration of the target proteins of drugs.

## MATERIALS AND METHODS

### Sensitivity of *Phytophthora capsici* to fluopicolide *in vitro*

Five sensitive *P. capsici* isolates (LT1534, BYA5, JA8, LP3, and 12-11) were used for fluopicolide adaption. The LT-7, LT-8, LT-10, B-1, B-12, J-7, J-8, J-10, J-12, L-2, L-3, 11-1, 11-2, 11-3, 11-5, 11-6, and 11-7 resistant mutants were obtained from our previous research (*23*). The sensitivity of all isolates to fluopicolide (99% a.i.) was determined *in vitro*, using the mycelium growth rate method on V8 media (100 mL of V8 juice, 1.4 g of CaCO_3_, and 15 g of agar, adjusted with distilled water to 1 L) in the dark at 25℃, adjusted to final concentrations of 0, 0.125, 0.25, 0.5, 0.625, 1.25, 5, 10, 25, and 50 μg/mL of fluopicolide. Plates containing only 0.1% (v/v) dimethyl sulfoxide (DMSO) in the medium were used as the controls. Each treatment contained three replicate plates. The EC_50_ value of each isolate was obtained according to the method described by Lu et al. (*47*).

### Construction of DNA pools

The mating types were determined using the mating-type standard isolates PCAS1 (A1) and PCAS2 (A2), which were gifted by Professor Michael Coffey of the University of California, Riverside, USA (*48*). LT-10 (A2) and BYA5 (A1) were cultured on V8 medium at 25 ℃ for 45 days to produce oospores, using a confrontation culture method. Then, conventional methods were used to extract oospores and induce germination of the F1 generation (*49, 50*). Isolates 381 (A1) and 414 (A2) were selected from the F1 generation for sister crossing to obtain the F2 generation. The DNA of 51 resistant isolates and 32 sensitive isolates from the F2 generation were extracted by the CTAB method (*51*). The same amount of DNA was mixed to construct a sensitive mixing pool and a resistant mixing pool (Tables S8 and S9).

### Generation and processing of whole-genome sequencing data

The genome database used for sequencing analysis is *P. capsici* LT1534 v11.0 (https://mycocosm.jgi.doe.gov/Phyca11/Phyca11.home.html), which is assembled to the scaffold level. cDNA libraries were prepared using the Illumina TruSeq RNA sample preparation kit v.2 (Illumina, CA, USA). Sequencing was performed on an Illumina HiSeq XTen/NovaSeq/BGI platform by a commercial service (Biomarker Technologies, Beijing, China). High-throughput sequencing with the Illumina HiSeq 2500 system generated approximately 6.8 and 55.5 million clean reads for the parent strain and two pools (sensitive pool and resistant pool), respectively. Finally, high-quality sequence data were obtained for association analysis.

### Association analysis

We used Euclidean distance (ED) and SNP-index algorithms to perform association analysis. The ED algorithm is a method for searching markers with significant differences between the pools, according to the sequencing data, and evaluating the associated regions between markers and traits. The ED is calculated using the following equations (*52*):

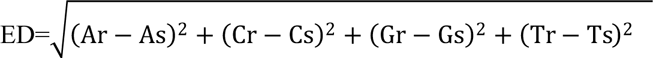

In which Ar, Cr, Gr, and Tr represent the frequency of A, C, G, and T in the resistant pool. As, Cs, Gs, and Ts represent the frequency of A, C, G, and T in the sensitive pool. The sequencing depth of differential SNPs in each mixed pool was counted to calculate the ED on each site. To eliminate the background noise, the ED value to the 5th power was used as the associated value.

The SNP index was calculated to find the significant differences in the genotype frequency between the pools, indicated by Δ(SNP index). A stronger SNP-trait linkage leads to a Δ(SNP index) value closer to 1. The SNP-index method is based on gene linkages and requires the position information of candidate SNPs on chromosomes to be fitted and analyzed. However, the genome of *P. capsici* was only assembled to the scaffold level. Thus, we calculated the threshold through a permutation test to screen out candidate association sites that exceeded the threshold. The probability of associated sites on the scaffold was calculated using a hypergeometric distribution. The Benjamini-Hochberg procedure was performed for multiple test corrections and to calculate the FDR. Finally, the key sites with FDR < 0.01 and a corresponding scaffold were screened out (*53-55*).

### Localization of Pcα-actinin and PcVHA-a

The *GFP* gene was fused to the C-terminus of Pcα-actinin by ligation to *Age*I/*Nhe*I-digested pTOR vector, and the mCherry gene was fused to the C-terminus of PcVHA-a by ligation to *Sac*II/*Spe*I-digested pYF3. The primers used for sequence amplification were Pcα-actinin-GFP-F (5’→3’): GCTTTAATTAAATGGCCGGATACAACGAGGA; Pcα-actinin-GFP-R (5’→3’): TCATGCTAGCGACGCTGAAGATAAAGTCCG; PcVHA-a-mCherry-F (5’→3’): TATCGATAGGCCTCCGCGGATGAAGTGGCTCCGCTCGG; PcVHA-a-mCherry-R (5’→3’): TGCTCACCATACTAGTCTAGGGCAGCTGCGAGTCC. The two PCR fragments were infused into vectors, using the In-Fusion® HD Cloning Kit (Takara Biomedical Technology Co., Ltd, Beijing, China). The pTOR-actinin-GFP and pYF3-PcVHA-a - mCherry vectors were introduced into *P. capcisi* BYA5 by polyethylene glycol (PEG)-mediated protoplast transformation (*56*). The empty vectors pYF3-GFP and pTOR-mCherry were transformed into BYA5 as a negative control. The living hyphae of the transformants were picked from V8 broth containing 50 μg/mL of G418 after 2 days of growth. The hyphae were observed using an FV-3000 confocal microscope (Olympus Life Science).

### CRISPR/Cas9-mediated site mutation and gene knock out

*PcVHA-a* containing four-point mutations was detected in the PcVHA-a of the fluopicolide-resistant mutants (G767E, N771Y, N846S, and K847R), which were transformed into the sensitive wild-type isolate BYA5 to validate their association with fluopicolide resistance. According to the previous CRISPR/Cas9 system protocol (*56*), donor vectors were constructed by cloning the homologous 1000 bp 5’ and 3’ flanking arms outside the target sites, and the two PCR fragments were infused into the pBluescript II KS+ vector. One sgRNA (PcVHA-a-sgRNA-ACAGGTTGTTCCAGCCCTTG) was designed using online sgRNA Designer (http://www.broadinstitute.org/rnai/public/analysis-tools/sgrnadesign). The sgRNA fragment was cloned into the pYF515-Ribo-sgRNA-Cas9 vector by T4 DNA ligase (New England BioLabs, Beijing, China). The Pcα-actinin knockout vectors were constructed as the point-mutations donor vectors, but the *Pcα-actinin* gene was replaced with the NPTII sgRNA sequence △Pcactinin-sgRNA-GATGTCTGACGAGCCGATGT. The PcVHA-a^G767A^ overexpression vectors were constructed using a homologous recombination method. The primers used in this study are listed in Table S10. The vectors were introduced into *P. capsici* BYA5 through a PEG-mediated protoplast transformation.

### Drug affinity responsive target stability (DARTS)

The protein target of fluopicolide was identified by an optimized DARTS strategy according to current methods (*16-18*, *31*, *57*, *58*). The reagents for membrane protein extraction from BYA5 included ultrapure water, Tris-HCl (pH = 7.5), Tris-HCl-glycerin (pH = 7.5, 10% glycerin), and PBS buffer (0.33 M sucrose, 0.1 mM EDTA, and 0.5 mM DTT). The reagents were treated with three concentrations of fluopicolide (0, 0.2, and 2 μg/mL) dissolved in DMSO.

The pretreatment steps of DARTS were as follows: BYA5 was cultured in a light incubator at 25 ℃ for 5–6 days to produce sporangia. The mixture of hyphae and sporangia was frozen using liquid nitrogen immediately after collection and was then ground into powder. The powdered sample was washed with ultrapure water, centrifuged at 10000 ×g for 2 min, and the supernatant was discarded. The sediment was suspended in 10 mM Tris-HCl and centrifuged at 5000 ×g for 10 min to obtain the supernatant. The supernatant was diluted with PBS buffer and centrifuged at 100000 ×g for 1 h. The supernatant was discarded and the sediment was dissolved with Tris-HCl-glycerin. Finally, a BCA protein assay kit (TIANGEN, Beijing, China) was used to measure the membrane protein concentration, and the concentration was adjusted to 1 mg/mL. The above operations were carried out at a low temperature, and protease inhibitors and phosphatase inhibitors were added to the reagents.

The membrane protein containing final concentrations of 0, 0.2, and 2 μg/mL of fluopicolide was obtained by the above operations. The following DARTS steps were applied: The membrane protein was incubated at 4℃ for 1 h and room temperature for 15 min; the samples were shaken 3–5 times during the incubation stage. Pronase (Roche Diagnostics, Indianapolis, USA) was added at concentrations of 1:500 and 1:1000 (m/m), and the same amount of ultrapure water was added to the controls. The samples were incubated at room temperature for 15 min. Digestion was stopped by adding 5× loading dye at 70°C for 10 min. The samples were detected by gel electrophoresis and stained with Coomassie blue dye. Specific gel bands that were identified in the fluopicolide treatment samples, compared to the DMSO controls, were cut out and prepared for mass spectrometry analysis.

### Western blot analysis

The 3× FLAG was fused to the N-terminus of PcVHA-a by ligation to the *Nhe*I*/Sac*II-digested pTOR vector. The primers used for sequence amplification were: 3× FLAG-PcVHA-a-F (5’→3’): CCTTGAGGTTGCTAGCATGAAGTGGCTCCGCTCGG; 3× FLAG-PcVHA-a-R (5’→3’): AGAAGTAGGCACCCCGCGGCTAGGGCAGCTGCGAGTCC. The construction and transformation of vectors were accomplished using the protocol described above. The total proteins of PcVHA-a overexpression transformants were extracted using a Thick-wall Microbial Protein Extraction Kit (Bestbio, Nanjing, China). The protein samples were resolved by 10% SDS-PAGE and transferred to PVDF membranes (Millipore, Billerica, MA) through standard electroblotting procedures. The primary antibodies anti-FLAG (Abways, AB0008), anti-β-tubulin (Abways, AB0039), and anti-β-actin (Abways, AB0035), and the secondary antibody HRP goat anti-mouse immunoglobulin G (IgG) (H+L) (Abways, B0102) were used. The protein was incubated with primary and secondary antibodies according to the manufacturer’s protocols. The PVDF membrane samples were detected with an enhanced chemiluminescence (ECL) kit (DiNing, Beijing, China), using a ChemiScope 6100 (CLiNX, Shanghai, China).

### Microscale thermophoresis (MST) assay

The mycelium samples of the mCherrry and PcVHA-a-mCherry transformants were cultured in liquid V8 at 25 ℃ for 2 days. Then, the total protein was extracted with the Thick-wall Microbial Protein Extraction Kit (Bestbio), incubated with different concentrations of fluopicolide, and loaded into silica capillaries (Polymicro Technologies). MST experiments were performed on a NanoTemper Monolith NT.115 (NanoTemper Technologies, Germany) with red filters at 25 °C and 20% MST power.

### Molecular docking

AlphaFold2 (AF2), developed by Google DeepMind, was used to predict the 3D structures of VHA-a from the amino acid sequences of *P. capsici* and *Salmo salar* (*35, 59*). The 3D model of HsVHA-a (PDB DOI: 10.2210/pdb6wm2/pdb) was downloaded from the PDB database. The quality of the yielded 3D model was assessed using MolProbity (*60, 61*). Molecular-docking simulations were performed using the AutoDock4.2.6 software embedded in YASARA. Fluopicolide and fluopimomide were used as ligands for docking. Fluopicolide and fluopimomide were docked into PcVHA-a. Fluopicolide was docked into the mutants of PcVHA-a (PcVHA-aG767E, PcVHA-aN771Y, PcVHA-aN846S, and PcVHA-aK847R), *S. salar* VHA-a3 (SsVHA-a3), and *Homo sapiens* VHA-a1 (HsVHA-a1). For the molecular-docking calculations, water molecules were deleted, a default pH of 7.4 was chosen, and the energy of the protein models was minimized using the molecular operating environment.

### Phylogenetic analysis

We used the VHA-a and α-actinin of different species for phylogenetic analysis. The protein sequences were obtained from the National Center for Biotechnology Information and were aligned by BLASTP. All steps of the phylogenetic analysis were completed on MEGA 11, and the following parameters were set: All the amino acid sequences were aligned by ClustalW, the statistical method was the maximum likelihood, the phylogeny test comprised 1000 bootstrap replications, and the substitution mode was the Poisson model.

### Statistical analysis

The Kd values of the MST assay are presented as means ± standard deviations (SD). The EC_50_ calculations of fungicide treatments for each isolate were performed according to the method of Lu et al. (*47*).

## ACKNOWLEDGMENTS

This work was funded by the China National Science Foundation (NO. 32001942) and the National Key R&D Program of China (grant no. 2022YFD1400900), and the Innovation Capability Support Plan of Shaanxi Province (2020TD-035).

## SUPPLEMENTARY MATERIALS

**Fig. S1.**
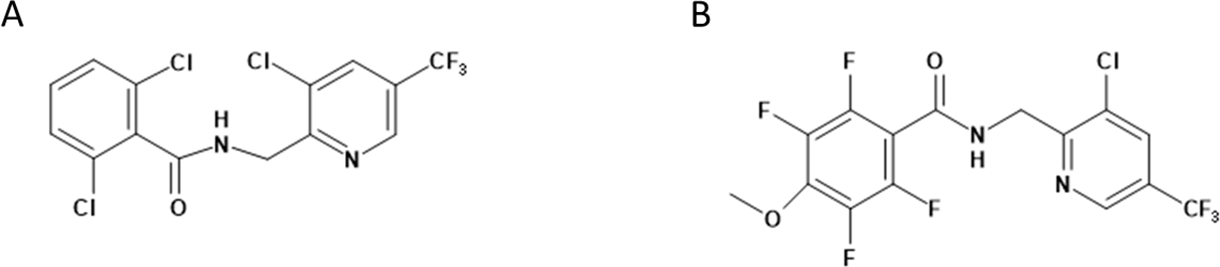
The structural formula of fluopicolide (A) and fluopimomide(B)

**Fig. S2.**
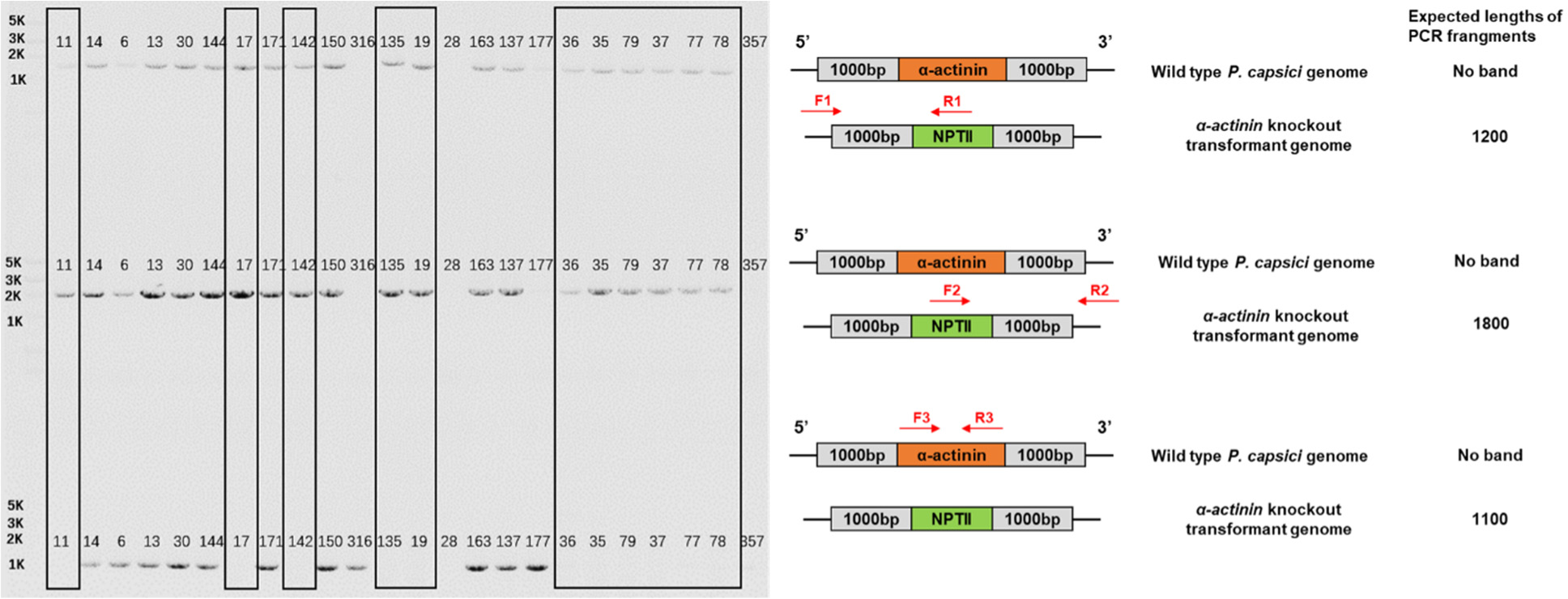
PCR verification of α-actinin knockout transformants in *Phytophthora capsici*. The homologous arm verification primers F1/R1 and F2/R2 can amplify the bands in the transformants 11, 17, 142, 135, 19, 36, 35, 79, 37, 77, 78, but not the α-actinin gene bands.

**Fig. S3.**
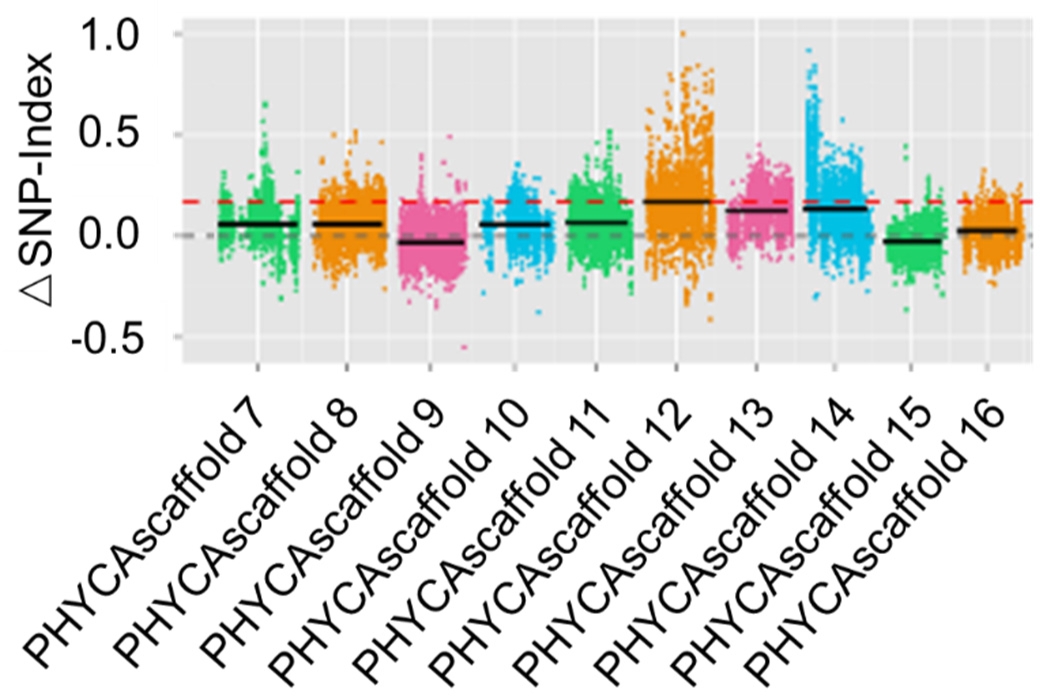
Distribution of SNP-index on scaffold. Coloured dots represents SNP-index(or ΔSNP-index) values of each SNP site. Black line indicates fitted SNP-index (or ΔSNP-index) value. The red line indicates linkage threshold at 99% of confidence interval.

**Fig. S4.**
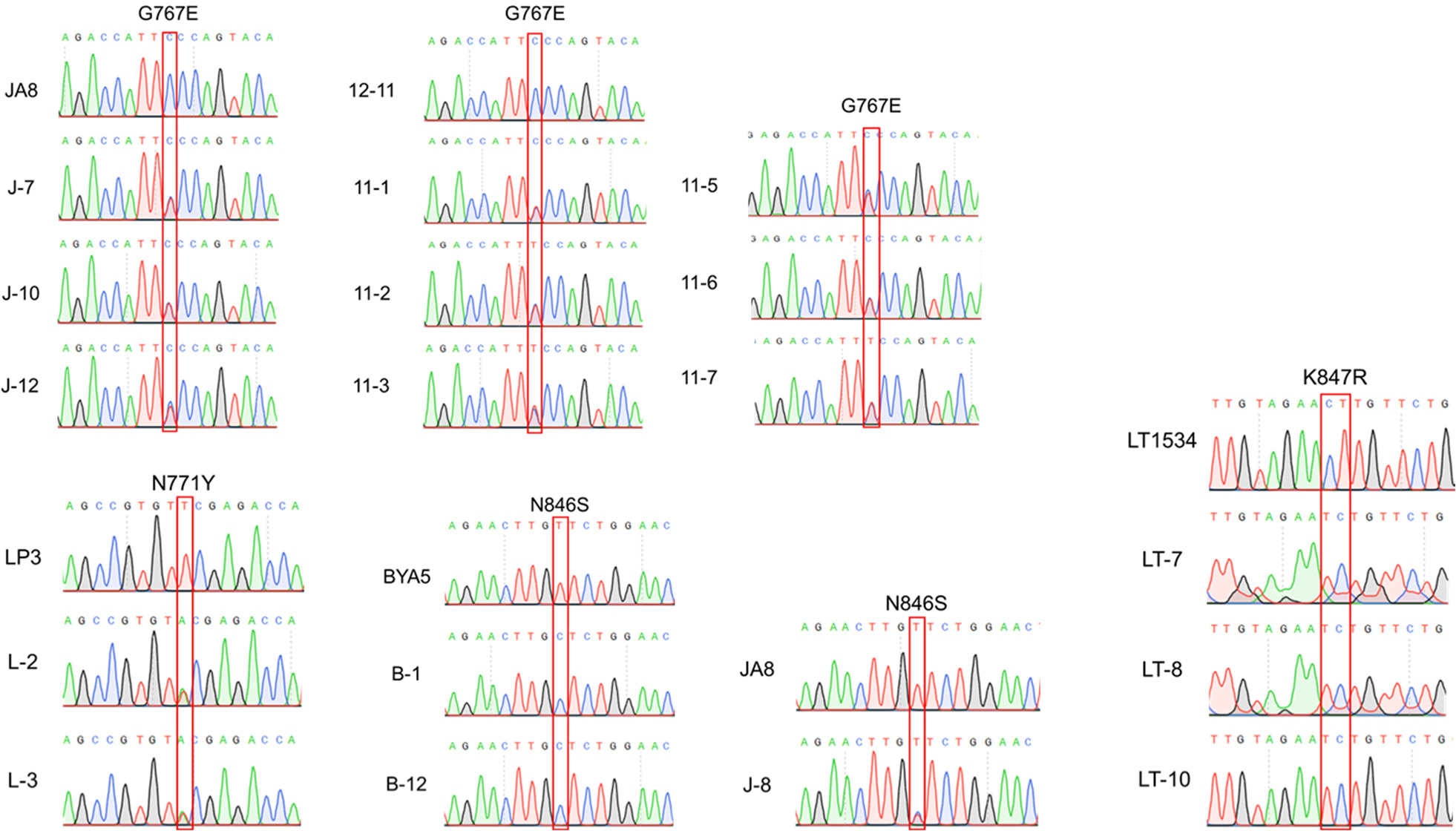
Multiple sequence alignment and sequencing chromatograms of the V-ATPase subunit a gene from fluopicolide resistant P. capsici mutants and the corresponding sensitive parental isolates.

**Fig. S5.**
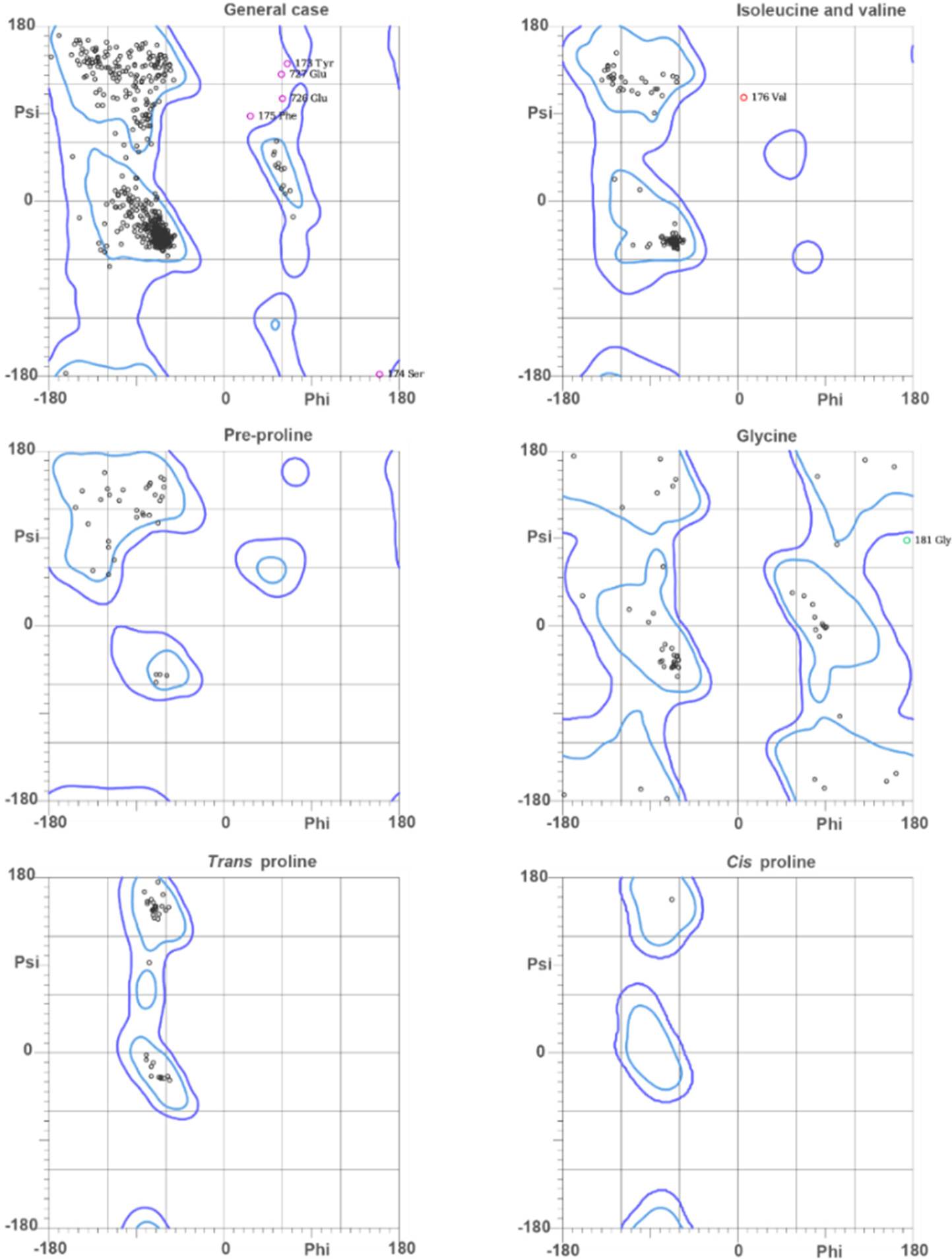
The MolProbity Ramachandran analysis of PcVHA-a. 96.8% (728/747) of all residues were in favored (98%) regions. 99.2% (746/747) of all residues were in allowed (>99.8%) regions.

**Fig. S6.**
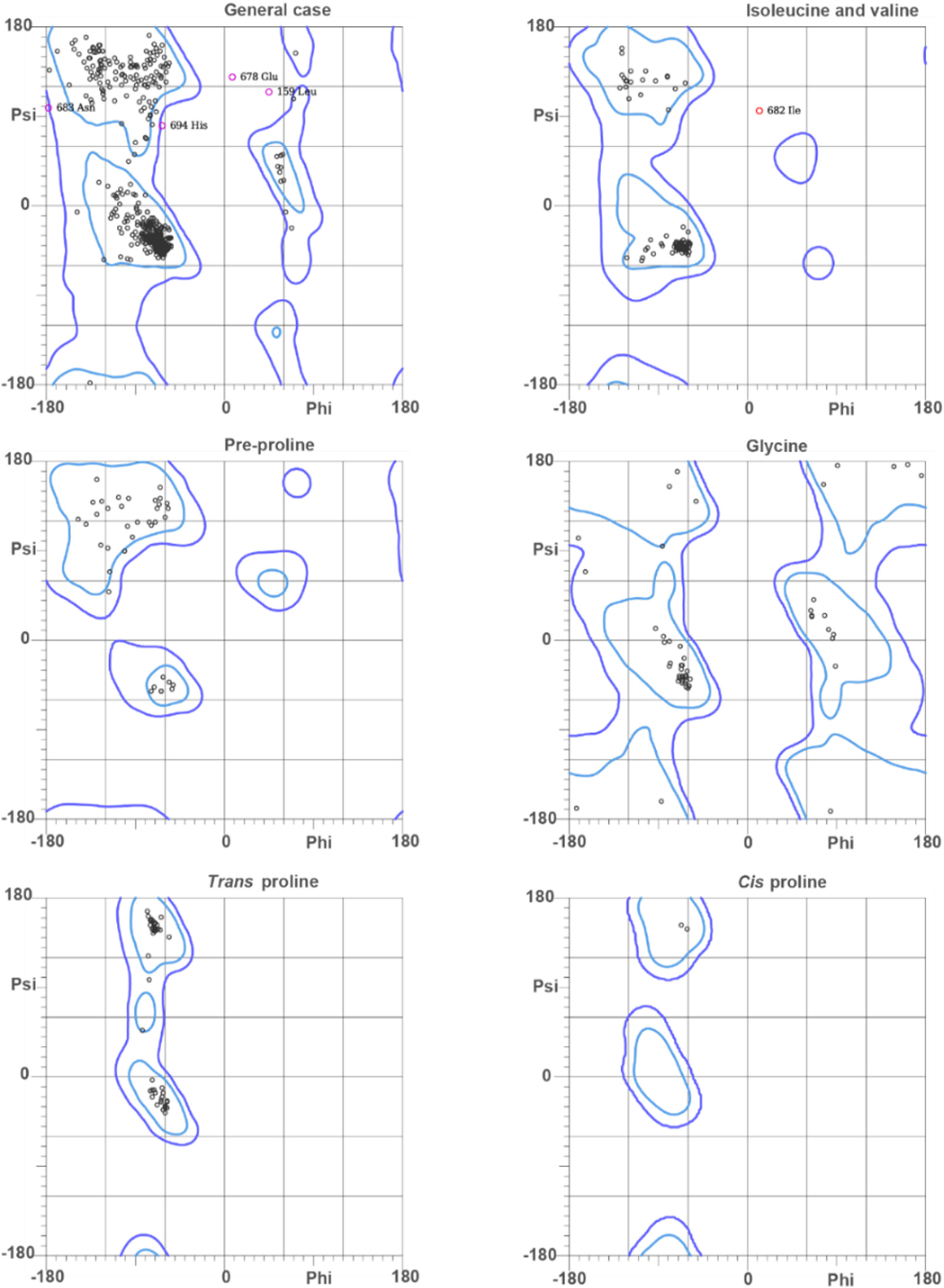
The MolProbity Ramachandran analysis of SsVHA-a3. 97.6% (728/747) of all residues were in favored (98%) regions. 99.4% (746/747) of all residues were in allowed (>99.8%) regions.

**Fig. S7.**
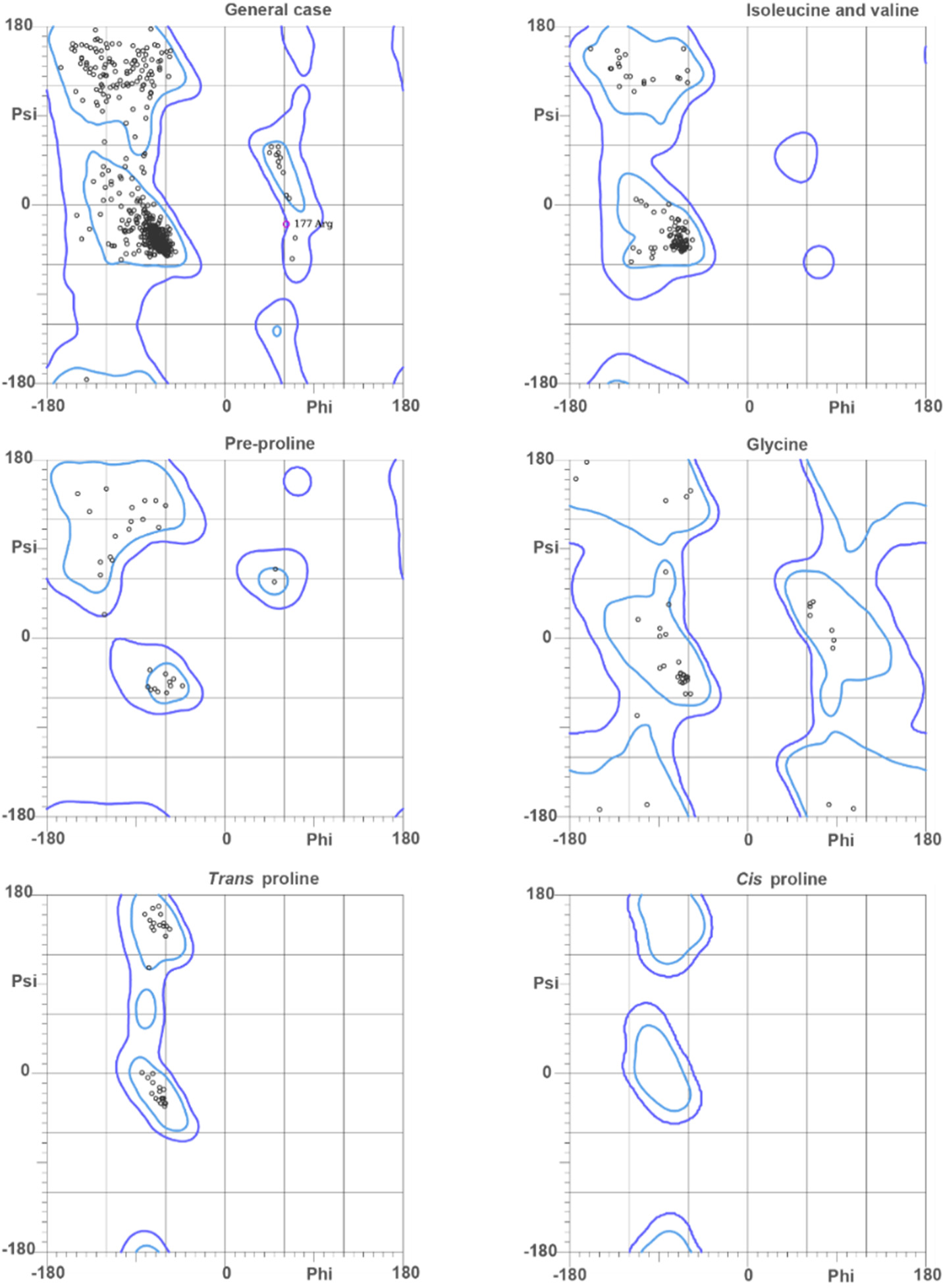
The MolProbity Ramachandran analysis of HsVHA-a1. 97.5% (728/747) of all residues were in favored (98%) regions. 99.9% (746/747) of all residues were in allowed (>99.8%) regions.

**Fig. S8.**
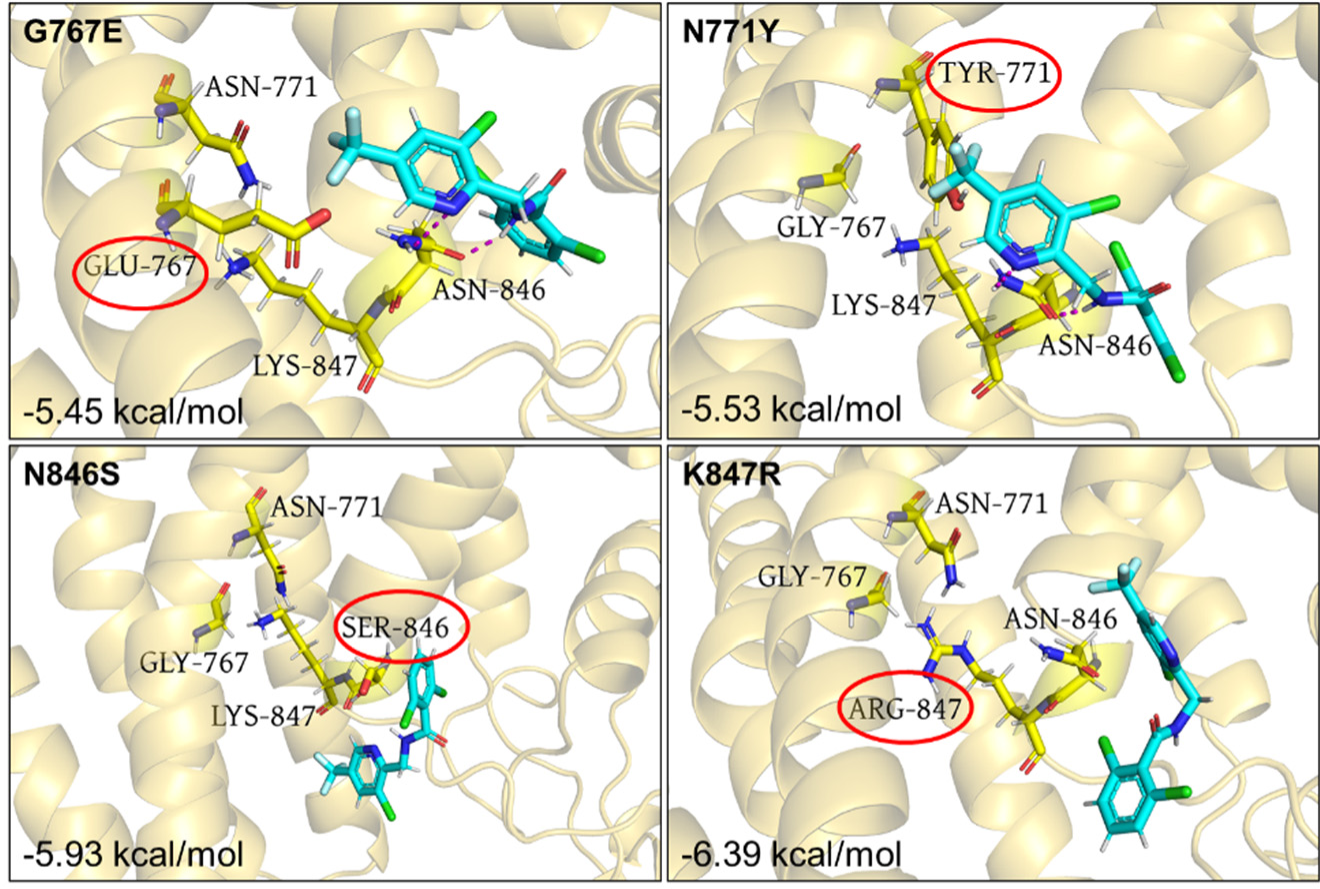
Molecular docking of fluopicolide to the binding sites V-ATPase subunit a from resistant mutants.

**Fig. S9.**
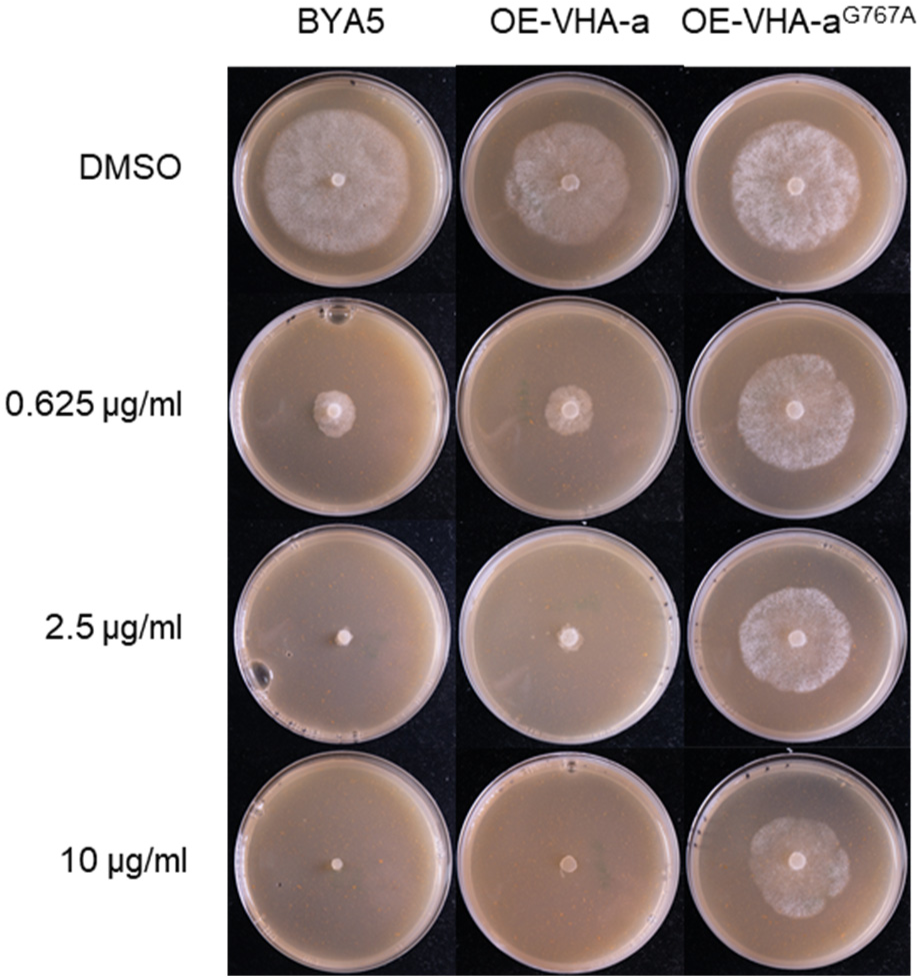
Sensitivity of G767A overexpression transformants to fluopicolide. The 767 site glycine was changed to the simple amino acid alanine to investigate the G767 on protein function, as well as on the ability to bind fluopicolide.

**Table S1.** Accession numbers of sequences listed in Figure 1A.

| Protein | accession number |
| --- | --- |
| *Homo sapiens* dystrophin | AAA53189.1 |
| *Homo sapiens* utrophin | NP_009055.2 |
| *Homo sapiens* α-actinin 1 | AAP36937.1 |
| *Homo sapiens* α-actinin 2 | NP_001094.1 |
| *Homo sapiens* α-actinin 3 | NP_001095.2 |
| *Homo sapiens* α-actinin 4 | NP_001308962.1 |
| *Homo sapiens* α-spectrin | XP_011508219.1 |
| *Homo sapiens* β-spectrin | NP_003119.2 |
| *Mus musculus* dystrophin | XP_006527831.1 |
| *Mus musculus* utrophin | NP_035812.3 |
| *Mus musculus* α-actinin 1 | NP_001333598.1 |
| *Mus musculus* α-actinin 2 | AAH89579.1 |
| *Mus musculus* α-actinin 3 | NP_038484.1 |
| *Mus musculus* α-actinin 4 | NP_001347478.1 |
| *Mus musculus* α-spectrin | NP_035595.2 |
| *Mus musculus* β-spectrin | NP_787030.2 |
| *Pc506502* | KAG1690892.1 |

**Table S2.** Significantly enriched genes by Euclidean Distance. All_Count, the number of SNPs on the Scaffold; Count, the number of SNPs obtained by the ED algorithm; P-value, the significance level; FDR, the probability after multiple test correction.

| Scaffold | AllCount | AssoCount | P-value | FDR |
| --- | --- | --- | --- | --- |
| PHYCAscaffold_12 | 9,542 | 292 | 0 | 0 |
| PHYCAscaffold_14 | 8,792 | 182 | 0 | 0 |
| PHYCAscaffold_184 | 66 | 36 | 0 | 0 |
| PHYCAscaffold_99 | 760 | 31 | 0 | 0 |
| PHYCAscaffold_245 | 50 | 9 | 2.22E-16 | 3.11E-14 |
| PHYCAscaffold_173 | 122 | 9 | 1.87E-12 | 2.18E-10 |
| PHYCAscaffold_585 | 8 | 4 | 7.54E-12 | 7.54E-10 |
| PHYCAscaffold_189 | 48 | 5 | 4.02E-09 | 3.52E-07 |
| PHYCAscaffold_541 | 6 | 2 | 3.80E-07 | 2.95E-05 |
| PHYCAscaffold_915 | 9 | 2 | 1.58E-06 | 0.00011091 |
| PHYCAscaffold_391 | 34 | 3 | 2.21E-06 | 0.00014089 |
| PHYCAscaffold_274 | 39 | 3 | 3.89E-06 | 0.00022663 |
| PHYCAscaffold_303 | 16 | 2 | 1.04E-05 | 0.00056086 |
| PHYCAscaffold_181 | 20 | 2 | 2.10E-05 | 0.00093701 |

**Table S3.** Significantly enriched genes by SNP-index.

| Scaffold | AllCount | AssoCount | P-value | FDR |
| --- | --- | --- | --- | --- |
| PHYCAscaffold_12 | 9,542 | 54 | 0 | 0 |
| PHYCAscaffold_14 | 8,792 | 18 | 2.56242E-09 | 9.4E-07 |
| PHYCAscaffold_585 | 8 | 2 | 3.51506E-09 | 9.4E-07 |
| PHYCAscaffold_199 | 59 | 3 | 1.09925E-08 | 2.2E-06 |
| PHYCAscaffold_245 | 50 | 2 | 1.22E-06 | 1.96E-04 |
| PHYCAscaffold_541 | 6 | 1 | 2.39E-06 | 3.21E-04 |
| PHYCAscaffold_441 | 13 | 1 | 1.24E-05 | 1.00E-03 |
| PHYCAscaffold_99 | 760 | 4 | 1.56E-05 | 2.00E-03 |
| PHYCAscaffold_565 | 21 | 1 | 3.33E-05 | 3.00E-03 |
| PHYCAscaffold_184 | 66 | 1 | 3.36E-04 | 0.005 |
| PHYCAscaffold_78 | 1777 | 5 | 8.91E-05 | 0.005 |
| PHYCAscaffold_136 | 85 | 1 | 5.57E-04 | 0.006 |
| PHYCAscaffold_126 | 108 | 1 | 8.96E-04 | 0.006 |
| PHYCAscaffold_73 | 243 | 1 | 4.00E-03 | 0.009 |

**Table S4.** Gene function annotation results of SNP and InDel in candidate regions. Annotated_Databases: Database applied in annotation; Gene Num: Annotated genes in candidate regions; Non_Syn Gene Num: Number of non-synonymous mutated genes between samples; FRAME_SHIFT Gene Num: Number of frameshift genes between samples.<colcnt=4>

| Annotated_databases | GeneNum | Non_Syn GeneNum | FRAME_SHIFT GeneNum |
| --- | --- | --- | --- |
| NR | 581 | 282 | 98 |
| NT | 585 | 284 | 98 |
| trEMBL | 583 | 284 | 98 |
| SwissProt | 137 | 66 | 19 |
| GO | 221 | 100 | 2.70E+01 |
| KEGG | 109 | 58 | 1.80E+01 |
| COG | 131 | 63 | 2.30E+01 |
| Total | 585 | 284 | 9.80E+01 |

**Table S5.** Determination of the sensitivity of Phytophthora capsici to fluopicolide. The EC_50_ of wild-type isolates BYA5, JA8, LP3,12-11, LT1534 and fluopicolide-resistant mutants J-7, J-8, J-10, J-12, 11-1, 11-2, 11-3, 11-5, 11-6, 11-7, L-2, L-3, B-1, B-12, LT-7, LT-8, LT-10 were was determined by Wu et al.(1).<colcnt=5>

| Isolate | Origin | Substitution | EC<sub>50</sub>(μg/ml) | Resistance factor |
| --- | --- | --- | --- | --- |
| BYA5 | Parent | - | 0.18/0.23 | - |
| B-1 | Mutant | N846S | 3.90 | 21.82 |
| B-12 | Mutant | N846S | 8.31 | 46.50 |
| 846-1 | Transformants | N846S | 4.34 | 18.87 |
| 846-2 | Transformants | N846S | 9.57 | 41.61 |
| 846-3 | Transformants | N846S | 2.99 | 13.00 |
| 767-1 | Transformants | G767E | >1000 | >2000 |
| 767-2 | Transformants | G767E | >1000 | >2000 |
| 767-3 | Transformants | G767E | >1000 | >2000 |
| 771-1 | Transformants | N771Y | >1000 | >2000 |
| 771-2 | Transformants | N771Y | 485.88 | >1000 |
| 771-3 | Transformants | N771Y | >1000 | >2000 |
| 847-1 | Transformants | K847R | 413.3 | >1000 |
| 847-2 | Transformants | K847R | >1000 | >2000 |
| 847-3 | Transformants | K847R | 538.22 | >1000 |
| 12-11 | Parent | - | 0.18 | - |
| 11-1 | Mutant | G767E | >100 | >500 |
| 11-2 | Mutant | G767E | >100 | >500 |
| 11-3 | Mutant | G767E | >100 | >500 |
| 11-5 | Mutant | G767E | >100 | >500 |
| 11-6 | Mutant | G767E | >100 | >500 |
| 11-7 | Mutant | G767E | >100 | >500 |
| LP3 | Parent | - | 0.22 | - |
| L-2 | Mutant | N771Y | >100 | >400 |
| L-3 | Mutant | N771Y | >100 | >400 |
| LT1534 | Parent | - | 0.08 | - |
| LT-7 | Mutant | K847R | >100 | >1000 |
| LT-8 | Mutant | K847R | >400 | >1000 |
| LT-10 | Mutant | K847R | >400 | >1000 |
| JA8 | Parent | - | 0.23 | - |
| J-7 | Mutant | G767E | >100 | >400 |
| J-8 | Mutant | N846S | 5.44 | 23.68 |
| J-10 | Mutant | G767E | >100 | >400 |
| J-12 | Mutant | G767E | >100 | >400 |

**Table S6.** VHA-a type and number of different species.

| Species name | Subunit classification |  |  |  |  |
| --- | --- | --- | --- | --- | --- |
|  | a | a1 | a2 | a3 | a4 |
| *Homo sapiens* | - | 37 | 1 | 2 | 1 |
| *Mus musculus* | - | 14 | 1 | 1 | 1 |
| *Danio rerio* | - | 2 | 1 | 1 | - |
| *Salmo salar* | - | - | - | 1 | - |
| *Anabarrilius grahami* | - | 2 | 2 | 2 | - |
| *Larimichthys crocea* | - | 2 | 1 | - | - |
| *Arabidopsis thaliana* | - | 1 | 1 | 1 | - |
| *Capsicum annuum* | - | 3 | - | 3 | - |
| *Phytophthora capsici* | 1 | - | - | - | - |
| *Phytophthora sojae* | 1 | - | - | - | - |
| *Phytophthora infestans* | 1 | - | - | - | - |
| *Phytophthora nicotianae* | 1 | - | - | - | - |
| *Pythium oligandrum* | 1 | - | - | - | - |
| *Plasmopara halstedii* | 1 | - | - | - | - |
| *Saccharomyces cerevisiae* | 2 | - | - | - | - |
| *Fusarium oxysporum* | 1 | - | - | - | - |
| *Colletotrichum fructicola* | 1 | - | - | - | - |
| *Verticillium longisporum* | 1 | - | - | - | - |
| *Trichoderma simmonsii* | 1 | - | - | - | - |

**Table S7.** Accession numbers of sequences listed in Figure 5A.

| Protein | Accession number |
| --- | --- |
| *Phytophthora sojae* VHA-a | EGZ19411.1 |
| *Phytophthora infestans* VHA-a | EEY62594.1 |
| *Phytophthora capsici* VHA-a | KAG1700981.1 |
| *Phytophthora nicotianae* VHA-a | KUF81858.1 |
| *Pythium aphanidermatum* VHA-a | PAG1_G008615 |
| *Plasmopara halstedii* VHA-a | XP_024571814.1 |
| *Homo sapiens* VHA-a1 | NP_001123493.1 |
| *Homo sapiens* VHA-a2 | NP_036595.2 |
| *Homo sapiens* VHA-a3 | NP_036595.2 |
| *Homo sapiens* VHA-a4 | NP_065683.2 |
| *Mus musculus* VHA-a1 | NP_001229979.1 |
| *Mus musculus* VHA-a2 | NP_035726.2 |
| *Mus musculus* VHA-a3 | NP_001129563.1 |
| *Mus musculus* VHA-a4 | NP_536715.3 |
| *Danio rerio* VHA-a1 | NP_997837.1 |
| *Danio rerio* VHA-a2 | NP_001116219.1 |
| *Danio rerio* VHA-a3 | NP_998234.1 |
| *Salmo salar* VHA-a3 | NP_001135285.1 |
| *Anabarrilius grahami* VHA-a1 | ROL51051.1 |
| *Anabarrilius grahami* VHA-a2 | ROI27727.1 |
| *Anabarrilius grahami* VHA-a3 | ROI47867.1 |
| *Larimichthys crocea* VHA-a1 | KAE8284578.1 |
| *Larimichthys crocea* VHA-a2 | KAE8292159.1 |
| *Saccharomyces cerevisiae* Stv1p | NP_013770.1 |
| *Saccharomyces cerevisiae* Vph1p | NP_014913.3 |
| *Fusarium oxysporum* VHA-a | KAG7421450.1 |
| *Colletotrichum fructicola* VHA-a | XP_031881304.1 |
| *Verticillium longisporum* VHA-a | KAG7105549.1 |
| *Trichoderma simmonsii* VHA-a | QYT04081.1 |

**Table S8.** The DNA of resistant pool.

| Isolate | 5 µg/ml inhibition<br>rate (%) | Concentration(ng/µL) | OD260/280 | OD260/230 |
| --- | --- | --- | --- | --- |
| 2-6 | < 50 | 687.8 | 1.90 | 2.20 |
| 2-8 | < 50 | 779.0 | 1.95 | 2.30 |
| 2-34 | < 50 | 170.7 | 1.95 | 2.13 |
| 2-20 | < 50 | 125.0 | 1.97 | 1.89 |
| 2-53 | < 50 | 254.0 | 2.05 | 2.13 |
| 2-82 | < 50 | 187.0 | 1.96 | 2.32 |
| 2-93 | < 50 | 448.0 | 1.94 | 2.16 |
| 2-38 | < 50 | 190.7 | 1.94 | 1.97 |
| 2-27 | < 50 | 862.9 | 1.97 | 2.20 |
| 2-31 | < 50 | 1270.0 | 1.98 | 2.35 |
| 2-313 | < 50 | 206.9 | 2.00 | 2.24 |
| 2-109 | < 50 | 121.0 | 1.90 | 2.09 |
| 2-111 | < 50 | 102.0 | 1.96 | 2.16 |
| 2-101 | < 50 | 181.0 | 1.97 | 2.47 |
| 2-135 | < 50 | 149.0 | 1.96 | 2.00 |
| 2-143 | < 50 | 1156.0 | 1.95 | 2.34 |
| 2-156 | < 50 | 219.0 | 1.93 | 2.07 |
| 2-153 | < 50 | 34.0 | 1.90 | 1.50 |
| 2-166 | < 50 | 966.7 | 1.70 | 2.02 |
| 2-171 | < 50 | 217.0 | 1.90 | 2.20 |
| 2-189 | < 50 | 137.0 | 1.89 | 1.84 |
| 2-149 | < 50 | 547.0 | 1.90 | 2.36 |
| 2-263 | < 50 | 131.0 | 1.95 | 2.17 |
| 2-229 | < 50 | 133.0 | 1.94 | 2.31 |
| 2-241 | < 50 | 117.0 | 2.00 | 2.30 |
| 2-251 | < 50 | 122.0 | 1.98 | 2.06 |
| 2-201 | < 50 | 123.8 | 1.97 | 2.21 |
| 2-228 | < 50 | 125.0 | 1.90 | 2.45 |
| 2-248 | < 50 | 160.0 | 1.97 | 2.10 |
| 2-223 | < 50 | 166.0 | 1.89 | 2.29 |
| 2-194 | < 50 | 122.6 | 1.97 | 1.64 |
| 2-244 | < 50 | 132.0 | 1.94 | 2.17 |
| 2-288 | < 50 | 1306.0 | 1.99 | 2.36 |
| 2-282 | < 50 | 220.0 | 1.98 | 2.22 |
| 2-286 | < 50 | 47.6 | 1.88 | 1.62 |
| 2-274 | < 50 | 212.0 | 1.97 | 2.27 |
| 2-268 | < 50 | 167.0 | 1.90 | 2.18 |
| 2-252 | < 50 | 184.0 | 1.92 | 2.45 |
| 2-243 | < 50 | 107.0 | 1.95 | 2.43 |
| 2-213 | < 50 | 98.6 | 1.95 | 2.23 |
| 2-297 | < 50 | 126.0 | 1.90 | 2.30 |
| 2-247 | < 50 | 110.0 | 1.98 | 2.40 |
| 2-204 | < 50 | 443.0 | 1.88 | 2.23 |
| 2-233 | < 50 | 135.0 | 1.91 | 2.30 |
| 2-203 | < 50 | 511.0 | 1.92 | 2.29 |
| 2-266 | < 50 | 137.0 | 1.85 | 2.39 |
| 2-214 | < 50 | 173.0 | 2.00 | 2.02 |
| 2-202 | < 50 | 252.0 | 1.95 | 2.15 |
| 2-273 | < 50 | 148.0 | 1.95 | 2.30 |
| 2-236 | < 50 | 164.0 | 1.90 | 2.30 |
| 2-224 | < 50 | 182.5 | 1.98 | 2.02 |

**Table S9.** The DNA of sensitive pool.

| Isolate | 5 µg/ml inhibition rate (%) | Concentration(ng/µL) | OD260/280 | OD260/230 |
| --- | --- | --- | --- | --- |
| 2-3 | 100 | 213.0 | 1.96 | 2.17 |
| 2-13 | 100 | 1570.0 | 1.94 | 2.47 |
| 2-26 | 100 | 168.0 | 1.98 | 2.08 |
| 2-28 | 100 | 68.0 | 1.90 | 2.00 |
| 2-35 | 100 | 163.0 | 2.08 | 2.36 |
| 2-47 | 100 | 97.0 | 2.04 | 1.87 |
| 2-49 | 100 | 1325.0 | 1.98 | 2.30 |
| 2-55 | 100 | 210.0 | 2.00 | 2.43 |
| 2-61 | 100 | 251.0 | 1.93 | 2.37 |
| 2-64 | 100 | 181.0 | 1.90 | 2.20 |
| 2-68 | 100 | 49.8 | 1.87 | 1.77 |
| 2-96 | 100 | 87.0 | 1.97 | 1.97 |
| 2-97 | 100 | 48.9 | 1.97 | 2.30 |
| 2-113 | 100 | 104.0 | 1.89 | 1.94 |
| 2-118 | 100 | 66.0 | 1.99 | 1.80 |
| 2-120 | 100 | 122.9 | 1.99 | 1.64 |
| 2-137 | 100 | 132.0 | 1.92 | 2.09 |
| 2-163 | 100 | 114.0 | 1.96 | 1.94 |
| 2-165 | 100 | 197.0 | 2.00 | 2.00 |
| 2-175 | 100 | 152.0 | 2.00 | 1.90 |
| 2-183 | 100 | 152.0 | 1.94 | 2.07 |
| 2-187 | 100 | 117.0 | 1.90 | 2.50 |
| 2-192 | 100 | 61.0 | 1.80 | 1.34 |
| 2-195 | 100 | 57.8 | 1.97 | 2.60 |
| 2-198 | 100 | 124.0 | 1.92 | 2.31 |
| 2-200 | 100 | 69.0 | 1.93 | 2.27 |
| 2-205 | 100 | 247.0 | 1.97 | 2.06 |
| 2-258 | 100 | 196.0 | 1.99 | 2.03 |
| 2-262 | 100 | 316.0 | 2.00 | 2.30 |
| 2-271 | 100 | 208.0 | 1.99 | 2.43 |
| 2-276 | 100 | 69.7 | 1.97 | 2.50 |
| 2-294 | 100 | 573.0 | 2.10 | 2.57 |
| 2-316 | 100 | 352.0 | 2.00 | 2.30 |
| 2-321 | 100 | 1166.0 | 2.02 | 2.04 |

**Table S10.**
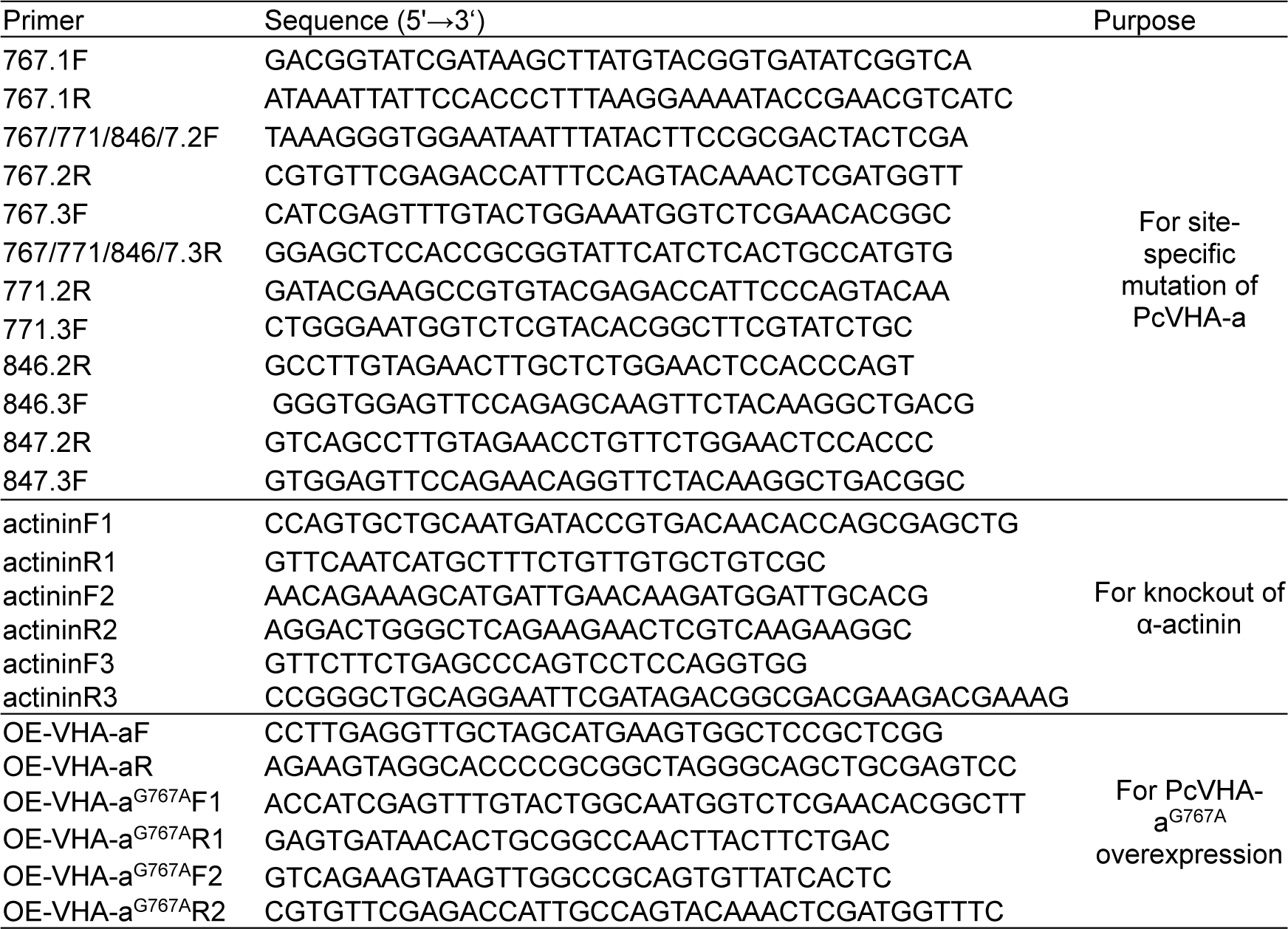
site-specific mutation and knockout Primers used for vector construction.

## REFERENCES AND NOTES

1 S. Kamoun, Molecular genetics of pathogenic oomycetes. Eukaryotic cell 2, 191-199 (2003).

1 A. Haverkort, P. Boonekamp, R. Hutten, E. Jacobsen, L. Lotz, G. Kessel, R. Visser, E. Van der Vossen, Societal costs of late blight in potato and prospects of durable resistance through cisgenic modification. Potato research 51, 47-57 (2008).

1 G. Jende, “Die Zellwand des Oomyceten *Phytophthora infestans* als Wirkort von Fungiziden”, thesis, Universitäts-und Landesbibliothek Bonn (2001).

1 B. Delvos, “Untersuchungen der Effekte von Iprovalicarb und Dimethomorph auf die Zellwand von *Phytophthora infestans*”, thesis, Dissertation, Mathematisch-Naturwissenschaftlichen Fakultät der Heinrich … (2009).

1 I. Andreassi, S. Gutteridge, S. Pember, J. Sweigard, E. Rehberg, Detection and screening method and materials useful in performance thereof. International Patent no. WO13/009971 Geneva: World Intellectual Property, (2013).

1 L. C. Davidse, A. E. Hofman, G. C. Velthuis, Specific interference of metalaxyl with endogenous RNA polymerase activity in isolated nuclei from *Phytophthora megasperma* f. sp. medicaginis. Experimental Mycology 7, 344-361 (1983).

1 R. Wollgiehn, E. Bräutigam, B. Schumann, D. Erge, Effect of metalaxyl on the synthesis of RNA, DNA and protein in Phytophthora nicotianae. Zeitschrift fûr allgemeine Mikrobiologie 24, 269-279 (1984).

1 Z. Zheng, Y. Hou, Y. Cai, Y. Zhang, Y. Li, M. Zhou, Whole-genome sequencing reveals that mutations in myosin-5 confer resistance to the fungicide phenamacril in Fusarium graminearum. Scientific reports 5, 1-9 (2015).

1 X. Qiao, X. Zhang, Z. Zhou, L. Guo, W. Wu, S. Ma, X. Zhang, C. Montell, J. Huang, An insecticide target in mechanoreceptor neurons. Science Advances 8, eabq3132 (2022).

1 J. Xu, X. Shao, Y. Li, Y. Wei, F. Xu, H. Wang, Metabolomic analysis and mode of action of metabolites of tea tree oil involved in the suppression of Botrytis cinerea. Frontiers in microbiology 8, 1017 (2017).

1 Z. Hu, T. Dai, L. Li, P. Liu, X. Liu, Use of GC–MS based metabolic fingerprinting for fast exploration of fungicide modes of action. BMC microbiology 19, 1-10 (2019).

1 J. Mu, S. Huang, S. Liu, Q. Zeng, M. Dai, Q. Wang, J. Wu, S. Yu, Z. Kang, D. Han, Genetic architecture of wheat stripe rust resistance revealed by combining QTL mapping using SNP-based genetic maps and bulked segregant analysis. Theoretical and Applied Genetics 132, 443-455 (2019).

1 A. Altinkut, N. Gozukirmizi, Search for microsatellite markers associated with water-stress tolerance in wheat through bulked segregant analysis. Molecular biotechnology 23, 97-106 (2003).

1 M. H. Bello, S. M. Moghaddam, M. Massoudi, P. E. McClean, P. B. Cregan, P. N. Miklas, Application of in silico bulked segregant analysis for rapid development of markers linked to Bean common mosaic virusresistance in common bean. BMC genomics 15, 1-13 (2014).

1 D. Shoba, N. Manivannan, P. Vindhiyavarman, S. Nigam, SSR markers associated for late leaf spot disease resistance by bulked segregant analysis in groundnut (Arachis hypogaea L.). Euphytica 188, 265-272 (2012).

1 B. Lomenick, R. Hao, N. Jonai, R. M. Chin, M. Aghajan, S. Warburton, J. Wang, R. P. Wu, F. Gomez, J. A. Loo, Target identification using drug affinity responsive target stability (DARTS). Proceedings of the National Academy of Sciences 106, 21984-21989 (2009).

1 Y. Cai, Y. Zheng, J. Gu, S. Wang, N. Wang, B. Yang, F. Zhang, D. Wang, W. Fu, Z. Wang, Betulinic acid chemosensitizes breast cancer by triggering ER stress-mediated apoptosis by directly targeting GRP78. Cell death & disease 9, 636 (2018).

1 R. M. Chin, X. Fu, M. Y. Pai, L. Vergnes, H. Hwang, G. Deng, S. Diep, B. Lomenick, V. S. Meli, G. C. Monsalve, The metabolite α-ketoglutarate extends lifespan by inhibiting ATP synthase and TOR. Nature 510, 397-401 (2014).

1 Y.-D. Park, W. Sun, A. Salas, A. Antia, C. Carvajal, A. Wang, X. Xu, Z. Meng, M. Zhou, G. J. Tawa, Identification of multiple cryptococcal fungicidal drug targets by combined gene dosing and drug affinity responsive target stability screening. MBio 7, e01073-01016 (2016).

1 V. Toquin, F. Barja, C. Sirven, S. Gamet, L. Mauprivez, P. Peret, M.-P. Latorse, J.-L. Zundel, F. Schmitt, M.-H. Lebrun, Novel tools to identify the mode of action of fungicides as exemplified with fluopicolide. Recent Developments in Management of Plant Diseases, 19-36 (2009).

1 M. Broderick, S. Winder, Spectrin, α-actinin, and dystrophin. Advances in protein chemistry 70, 203-246 (2005).

1 J. Wu, “The Resistance Risk and Resistant Mechanism of Phytophthora capsici to Fluopicolide”, thesis, China Agricultural University, China (2020).

1 J. Wu, Z. Xue, J. Miao, F. Zhang, X. Gao, X. Liu, Sensitivity of different developmental stages and resistance risk assessment of Phytophthora capsici to fluopicolide in China. Frontiers in Microbiology 11, 185 (2020).

1 V. Toquin, M. P. Latorse, R. Beffa, Fluopicolide: A New Anti-oomycete Fungicide. Modern Crop Protection Compounds 2, 871-878 (2019).

1 M. Cotado-Sampayo, P. O. Ramos, R. O. Perez, M. Ojha, F. Barja, Specificity of commercial anti-spectrin antibody in the study of fungal and Oomycete spectrin: cross-reaction with proteins other than spectrin. Fungal Genetics and Biology 45, 1008-1015 (2008).

1 M. Blum, M. Waldner, G. Olaya, Y. Cohen, U. Gisi, H. Sierotzki, Resistance mechanism to carboxylic acid amide fungicides in the cucurbit downy mildew pathogen Pseudoperonospora cubensis. Pest management science 67, 1211-1214 (2011).

1 Z. Pang, J. Shao, L. Chen, X. Lu, J. Hu, Z. Qin, X. Liu, Resistance to the novel fungicide pyrimorph in *Phytophthora capsici*: risk assessment and detection of point mutations in CesA3 that confer resistance. PLoS One 8, e56513 (2013).

1 Y. Zhou, L. Chen, J. Hu, H. Duan, D. Lin, P. Liu, Q. Meng, B. Li, N. Si, C. Liu, Resistance mechanisms and molecular docking studies of four novel QoI fungicides in *Peronophythora litchii*. Scientific Reports 5, 1-10 (2015).

1 M. Cai, J. Miao, X. Song, D. Lin, Y. Bi, L. Chen, X. Liu, B. M. Tyler, C239S mutation in the β-tubulin of *Phytophthora sojae* confers resistance to zoxamide. Frontiers in microbiology 7, 762 (2016).

1 J. Miao, M. Cai, X. Dong, L. Liu, D. Lin, C. Zhang, Z. Pang, X. Liu, Resistance assessment for oxathiapiprolin in *Phytophthora capsici* and the detection of a point mutation (G769W) in PcORP1 that confers resistance. Frontiers in Microbiology 7, 615 (2016).

1 D. Kim, H.-Y. Hwang, J. Y. Kim, J. Y. Lee, J. S. Yoo, G. r. Marko-Varga, H. J. Kwon, FK506, an immunosuppressive drug, induces autophagy by binding to the V-ATPase catalytic subunit A in neuronal cells. Journal of proteome research 16, 55-64 (2017).

1 M. Gao, Y. He, X. Yin, X. Zhong, B. Yan, Y. Wu, J. Chen, X. Li, K. Zhai, Y. Huang, Ca2+ sensor-mediated ROS scavenging suppresses rice immunity and is exploited by a fungal effector. Cell 184, 5391-5404. e5317 (2021).

1 W. Shao, R. Sharma, M. H. Clausen, H. V. Scheller, Microscale thermophoresis as a powerful tool for screening glycosyltransferases involved in cell wall biosynthesis. Plant methods 16, 1-12 (2020).

1 S. T. Chen, N. Y. He, J. H. Chen, F. Q. Guo, Identification of core subunits of photosystem II as action sites of HSP 21, which is activated by the GUN 5-mediated retrograde pathway in Arabidopsis. The Plant Journal 89, 1106-1118 (2017).

1 J. Jumper, R. Evans, A. Pritzel, T. Green, M. Figurnov, O. Ronneberger, K. Tunyasuvunakool, R. Bates, A. Žídek, A. Potapenko, Highly accurate protein structure prediction with AlphaFold. Nature 596, 583-589 (2021).

1 K. C. Jefferies, D. J. Cipriano, M. Forgac, Function, structure and regulation of the vacuolar (H+)-ATPases. Archives of biochemistry and biophysics 476, 33-42 (2008).

1 H. Wieczorek, M. Putzenlechner, W. Zeiske, U. Klein, A vacuolar-type proton pump energizes K+/H+ antiport in an animal plasma membrane. Journal of Biological Chemistry 266, 15340-15347 (1991).

1 M. Forgac, Vacuolar ATPases: rotary proton pumps in physiology and pathophysiology. Nature reviews Molecular cell biology 8, 917-929 (2007).

1 M. McConnell, S. Feng, W. Chen, G. Zhu, D. Shen, S. Ponnazhagan, L. Deng, Y.-P. Li, Osteoclast proton pump regulator Atp6v1c1 enhances breast cancer growth by activating the mTORC1 pathway and bone metastasis by increasing V-ATPase activity. Oncotarget 8, 47675 (2017).

1 L. Stransky, K. Cotter, M. Forgac, The function of V-ATPases in cancer. Physiological reviews 96, 1071-1091 (2016).

1 S. Fais, A. De Milito, H. You, W. Qin, Targeting vacuolar H+-ATPases as a new strategy against cancer. Cancer research 67, 10627-10630 (2007).

1 Ó. Ö. Hálfdánarson, K. Fall, M. H. Ogmundsdottir, S. H. Lund, E. Steingrímsson, H. M. Ogmundsdottir, H. Zoega, Proton pump inhibitor use and risk of breast cancer, prostate cancer, and malignant melanoma: an Icelandic population-based case-control study. Pharmacoepidemiology and Drug Safety 28, 471-478 (2019).

1 A. Kulshrestha, G. K. Katara, A. Kumar, K. D. Beaman, In silico identification of specific inhibitor (s) of tumor associated Vacuolar ATPase ‘a2’isoform: A critical target for anti-cancer drug development. Cancer Research 80, 5166-5166 (2020).

1 E. J. Bowman, L. A. Graham, T. H. Stevens, B. J. Bowman, The bafilomycin/concanamycin binding site in subunit c of the V-ATPases from *Neurospora crassa* and *Saccharomyces cerevisiae*. Journal of Biological Chemistry 279, 33131-33138 (2004).

1 L. Lu, Z. Qi, Q. Li, W. Wu, Validation of the target protein of insecticidal dihydroagarofuran sesquiterpene polyesters. Toxins 8, 79 (2016).

1 R. Wang, J. Wang, A. Hassan, C.-H. Lee, X.-S. Xie, X. Li, Molecular basis of V-ATPase inhibition by bafilomycin A1. Nature communications 12, 1782 (2021).

1 X. H. Lu, S. S. Zhu, Y. Bi, X. L. Liu, J. J. Hao, Baseline sensitivity and resistance-risk assessment of *Phytophthora capsici* to iprovalicarb. Phytopathology 100, 1162-1168 (2010).

1 L. Bower, M. Coffey, Development of laboratory tolerance to phosphorous acid, fosetyl-Al, and metalaxyl in *Phytophthora capsici*. Canadian Journal of Plant Pathology 7, 1-6 (1985).

1 P. Ann, W. Ko, Induction of oospore germination of Phytophthora parasitica. Phytopathology 78, 335-338 (1988).

1 Q. Meng, X. Cui, Y. Bi, Q. Wang, J. Hao, X. Liu, Biological and genetic characterization of *Phytophthora capsici* mutants resistant to flumorph. Plant Pathology 60, 957-966 (2011).

1 M. Uzunova, W. Ecke, K. Weissleder, G. Röbbelen, Mapping the genome of rapeseed (Brassica napus L.). I. Construction of an RFLP linkage map and localization of QTLs for seed glucosinolate content. Theoretical and Applied Genetics 90, 194-204 (1995).

1 J. T. Hill, B. L. Demarest, B. W. Bisgrove, B. Gorsi, Y.-C. Su, H. J. Yost, MMAPPR: mutation mapping analysis pipeline for pooled RNA-seq. Genome research 23, 687-697 (2013).

1 Y. Deng, J. Li, S. Wu, Y. Zhu, Y. Chen, F. He, Integrated nr database in protein annotation system and its localization. Comput Eng 32, 71-74 (2006).

1 M. Ashburner, C. A. Ball, J. A. Blake, D. Botstein, H. Butler, J. M. Cherry, A. P. Davis, K. Dolinski, S. S. Dwight, J. T. Eppig, Gene ontology: tool for the unification of biology. Nature genetics 25, 25-29 (2000).

1 H. Takagi, A. Abe, K. Yoshida, S. Kosugi, S. Natsume, C. Mitsuoka, A. Uemura, H. Utsushi, M. Tamiru, S. Takuno, QTL-seq: rapid mapping of quantitative trait loci in rice by whole genome resequencing of DNA from two bulked populations. The Plant Journal 74, 174-183 (2013).

1 Y. Fang, B. M. Tyler, Efficient disruption and replacement of an effector gene in the oomycete P hytophthora sojae using CRISPR/C as9. Molecular plant pathology 17, 127-139 (2016).

1 J. Krucinska, E. Falcone, H. Erlandsen, A. Hazeen, M. N. Lombardo, A. Estrada, V. L. Robinson, A. C. Anderson, D.L. Wright, Structural and functional studies of bacterial enolase, a potential target against Gram-negative pathogens. Biochemistry 58, 1188-1197 (2019).

1 V. F. Lazarev, D. V. Sverchinsky, E. R. Mikhaylova, P. I. Semenyuk, E. Y. Komarova, S. A. Niskanen, A. D. Nikotina, A. V. Burakov, V. G. Kartsev, I. V. Guzhova, Sensitizing tumor cells to conventional drugs: HSP70 chaperone inhibitors, their selection and application in cancer models. Cell death & disease 9, 41 (2018).

1 M. Mirdita, K. Schütze, Y. Moriwaki, L. Heo, S. Ovchinnikov, M. Steinegger, ColabFold: making protein folding accessible to all. Nature methods 19, 679-682 (2022).

1 J. Liu, X. Yang, Y. Zhang, Characterization of a lambda-cyhalothrin metabolizing glutathione S-transferase CpGSTd1 from Cydia pomonella (L.). Applied microbiology and biotechnology 98, 8947-8962 (2014).

1 V. B. Chen, W. B. Arendall, J. J. Headd, D. A. Keedy, R. M. Immormino, G. J. Kapral, L. W. Murray, J. S. Richardson, D. C. Richardson, MolProbity: all-atom structure validation for macromolecular crystallography. Acta Crystallographica Section D: Biological Crystallography 66, 12-21 (2010).

