## Supplementary Materials 2-F5h for "Vacuolar H^+^-ATPase subunit a was identified as the target protein of the oomycete inhibitor fluopicolide"

| Family | Member | Protein ID | Score | Mass | Num. of matches | Num. of significant matches | Num. of sequences | Num. of significant sequences | emPAI | Gene ID |
| --- | --- | --- | --- | --- | --- | --- | --- | --- | --- | --- |
| 3 | 1 | 558578 | 1176 | 138453 | 55 | 55 | 28 | 28 | 1.53 | >jgi\|Phyca11\|558578\|estExt2_Genewise1.C_PHYCAscaffold_20133 |
| 4 | 2 | 509820 | 1155 | 86980 | 47 | 47 | 23 | 23 | 2.25 | >jgi\|Phyca11\|509820\|fgenesh2_kg.PHYCAscaffold_50_#_21_#_Contig158.1 |
| 5 | 3 | 569720 | 814 | 58735 | 40 | 40 | 24 | 24 | 5.13 | >jgi\|Phyca11\|569720\|estExt2_Genewise1.C_PHYCAscaffold_330350 |
| 6 | 4 | 525392 | 788 | 94498 | 37 | 37 | 22 | 22 | 1.83 | >jgi\|Phyca11\|525392\|estExt2_fgenesh1_pm.C_PHYCAscaffold_30158 |
| 7 | 5 | 577453 | 696 | 46059 | 28 | 28 | 18 | 18 | 4.26 | >jgi\|Phyca11\|577453\|estExt2_Genewise1.C_PHYCAscaffold_1190011 |
| 8 | 6 | 116367 | 692 | 96518 | 33 | 33 | 20 | 20 | 1.42 | >jgi\|Phyca11\|116367\|e_gw1.30.3.1 |
| 9 | 7 | 548602 | 679 | 69141 | 30 | 30 | 20 | 20 | 2.43 | >jgi\|Phyca11\|548602\|estExt2_Genewise1Plus.C_PHYCAscaffold_290268 |
| 10 | 8 | 98041 | 611 | 98789 | 24 | 24 | 20 | 20 | 1.37 | >jgi\|Phyca11\|98041\|e_gw1.2.524.1 |
| 11 | 9 | 565311 | 584 | 60786 | 33 | 33 | 19 | 19 | 3.35 | >jgi\|Phyca11\|565311\|estExt2_Genewise1.C_PHYCAscaffold_170498 |
| 12 | 10 | 505974 | 507 | 81161 | 21 | 21 | 14 | 14 | 1.09 | >jgi\|Phyca11\|505974\|fgenesh2_kg.PHYCAscaffold_17_#_59_#_4098047:1 |
| 14 | 11 | 555787 | 461 | 90055 | 26 | 26 | 17 | 17 | 1.24 | >jgi\|Phyca11\|555787\|estExt2_Genewise1Plus.C_PHYCAscaffold_780059 |
| 15 | 12 | 562095 | 443 | 195387 | 22 | 22 | 19 | 19 | 0.52 | >jgi\|Phyca11\|562095\|estExt2_Genewise1.C_PHYCAscaffold_80628 |
| 16 | 13 | 503562 | 427 | 116908 | 21 | 21 | 16 | 16 | 0.8 | >jgi\|Phyca11\|503562\|fgenesh2_kg.PHYCAscaffold_4_#_101_#_4100091:2 |
| 17 | 14 | 8926 | 409 | 59018 | 25 | 25 | 16 | 16 | 2.41 | >jgi\|Phyca11\|8926\|fgenesh1_pm.PHYCAscaffold_32_#_61 |
| 18 | 15 | 577365 | 383 | 67399 | 17 | 17 | 10 | 10 | 0.88 | >jgi\|Phyca11\|577365\|estExt2_Genewise1.C_PHYCAscaffold_1090022 |
| 19 | 16 | 108059 | 379 | 102760 | 19 | 19 | 14 | 14 | 0.79 | >jgi\|Phyca11\|108059\|e_gw1.14.147.1 |
| 20 | 17 | 576775 | 340 | 78845 | 13 | 13 | 10 | 10 | 0.72 | >jgi\|Phyca11\|576775\|estExt2_Genewise1.C_PHYCAscaffold_950063 |
| 21 | 18 | 568375 | 333 | 60822 | 15 | 15 | 8 | 8 | 0.75 | >jgi\|Phyca11\|568375\|estExt2_Genewise1.C_PHYCAscaffold_280427 |
| 22 | 19 | 511385 | 329 | 115018 | 13 | 13 | 10 | 10 | 0.45 | >jgi\|Phyca11\|511385\|fgenesh2_kg.PHYCAscaffold_84_#_1_#_Contig68.1 |
| 23 | 20 | 506531 | 328 | 68497 | 17 | 17 | 13 | 13 | 1.39 | >jgi\|Phyca11\|506531\|fgenesh2_kg.PHYCAscaffold_20_#_77_#_Contig273.1 |
| 24 | 1 | 537721 | 310 | 706535 | 17 | 17 | 14 | 14 | 0.09 | >jgi\|Phyca11\|537721\|estExt2_fgenesh1_pg.C_PHYCAscaffold_1090004 |
| 25 | 1 | 511907 | 300 | 96155 | 15 | 15 | 12 | 12 | 0.78 | >jgi\|Phyca11\|511907\|fgenesh2_kg.PHYCAscaffold_105_#_5_#_4098187:2 |
| 26 | 1 | 506857 | 296 | 75869 | 14 | 14 | 11 | 11 | 0.86 | >jgi\|Phyca11\|506857\|fgenesh2_kg.PHYCAscaffold_22_#_96_#_gi\|189084480\|gb\|BT031996.1\| |
| 27 | 1 | 509774 | 295 | 85263 | 14 | 14 | 11 | 11 | 0.73 | >jgi\|Phyca11\|509774\|fgenesh2_kg.PHYCAscaffold_49_#_66_#_Contig174.1 |
| 28 | 1 | 508653 | 288 | 74131 | 14 | 14 | 10 | 10 | 0.78 | >jgi\|Phyca11\|508653\|fgenesh2_kg.PHYCAscaffold_37_#_28_#_Contig1670.1 |
| 29 | 1 | 509101 | 288 | 125590 | 12 | 12 | 8 | 8 | 0.31 | >jgi\|Phyca11\|509101\|fgenesh2_kg.PHYCAscaffold_42_#_8_#_Contig2284.1 |
| 30 | 1 | 530396 | 275 | 90922 | 11 | 11 | 10 | 10 | 0.6 | >jgi\|Phyca11\|530396\|estExt2_fgenesh1_pm.C_PHYCAscaffold_630021 |
| 28 | 2 | 550410 | 269 | 73974 | 11 | 11 | 7 | 7 | 0.5 | >jgi\|Phyca11\|550410\|estExt2_Genewise1Plus.C_PHYCAscaffold_370115 |
| 31 | 1 | 532207 | 269 | 123366 | 13 | 13 | 7 | 7 | 0.27 | >jgi\|Phyca11\|532207\|estExt2_fgenesh1_pg.C_PHYCAscaffold_40153 |
| 32 | 1 | 505109 | 262 | 108406 | 7 | 7 | 6 | 6 | 0.27 | >jgi\|Phyca11\|505109\|fgenesh2_kg.PHYCAscaffold_11_#_55_#_4100846:2 |
| 33 | 1 | 529283 | 252 | 60530 | 16 | 16 | 13 | 13 | 1.5 | >jgi\|Phyca11\|529283\|estExt2_fgenesh1_pm.C_PHYCAscaffold_410034 |
| 34 | 1 | 503258 | 251 | 42657 | 10 | 10 | 7 | 7 | 1.01 | >jgi\|Phyca11\|503258\|fgenesh2_kg.PHYCAscaffold_3_#_123_#_Contig19.2 |
| 35 | 1 | 534200 | 247 | 152568 | 11 | 11 | 8 | 8 | 0.29 | >jgi\|Phyca11\|534200\|estExt2_fgenesh1_pg.C_PHYCAscaffold_210041 |
| 36 | 1 | 505046 | 246 | 80584 | 16 | 16 | 10 | 10 | 0.79 | >jgi\|Phyca11\|505046\|fgenesh2_kg.PHYCAscaffold_10_#_206_#_4097622:1 |
| 37 | 1 | 507039 | 236 | 57928 | 9 | 9 | 6 | 6 | 0.55 | >jgi\|Phyca11\|507039\|fgenesh2_kg.PHYCAscaffold_24_#_31_#_Contig1904.1 |
| 38 | 1 | 6113 | 228 | 57855 | 9 | 9 | 9 | 9 | 0.94 | >jgi\|Phyca11\|6113\|fgenesh1_pm.PHYCAscaffold_9_#_173 |
| 39 | 1 | 510859 | 220 | 48785 | 10 | 10 | 7 | 7 | 1.01 | >jgi\|Phyca11\|510859\|fgenesh2_kg.PHYCAscaffold_70_#_5_#_Contig3.1 |
| 40 | 1 | 107407 | 215 | 25619 | 9 | 9 | 7 | 7 | 2.16 | >jgi\|Phyca11\|107407\|e_gw1.13.765.1 |
| 41 | 1 |  | 209 | 39193 | 3 | 3 | 2 | 2 | 0.24 | SWISS-PROT:P12763 (Bos taurus) Alpha-2-HS-glycoprotein precursor |
| 42 | 1 | 529555 | 202 | 66199 | 10 | 10 | 8 | 8 | 0.67 | >jgi\|Phyca11\|529555\|estExt2_fgenesh1_pm.C_PHYCAscaffold_460008 |
| 43 | 1 | 104429 | 198 | 70424 | 14 | 14 | 12 | 12 | 1.07 | >jgi\|Phyca11\|104429\|e_gw1.9.149.1 |
| 44 | 1 | 545685 | 197 | 107998 | 13 | 13 | 13 | 13 | 0.67 | >jgi\|Phyca11\|545685\|estExt2_Genewise1Plus.C_PHYCAscaffold_180528 |
| 45 | 1 |  | 196 | 283140 | 5 | 5 | 4 | 4 | 0.06 | SWISS-PROT:Q86YZ3 Tax_Id=9606 Gene_Symbol=HRNR Hornerin |
| 46 | 1 | 511144 | 193 | 56958 | 11 | 11 | 11 | 11 | 1.28 | >jgi\|Phyca11\|511144\|fgenesh2_kg.PHYCAscaffold_76_#_45_#_4101945:241 |
| 47 | 1 | 506235 | 191 | 56205 | 13 | 13 | 10 | 10 | 1.13 | >jgi\|Phyca11\|506235\|fgenesh2_kg.PHYCAscaffold_18_#_108_#_Contig553.1 |
| 48 | 1 | 505083 | 190 | 69264 | 9 | 9 | 7 | 7 | 0.54 | >jgi\|Phyca11\|505083\|fgenesh2_kg.PHYCAscaffold_11_#_29_#_gi\|189084466\|gb\|BT031982.1\| |
| 49 | 1 | 17480 | 190 | 250999 | 13 | 13 | 12 | 12 | 0.23 | >jgi\|Phyca11\|17480\|fgenesh1_pg.PHYCAscaffold_28_#_15 |
| 50 | 1 | 570119 | 189 | 40022 | 10 | 10 | 7 | 7 | 1.1 | >jgi\|Phyca11\|570119\|estExt2_Genewise1.C_PHYCAscaffold_350311 |
| 51 | 1 | 509625 | 187 | 56331 | 10 | 10 | 9 | 9 | 0.98 | >jgi\|Phyca11\|509625\|fgenesh2_kg.PHYCAscaffold_48_#_28_#_Contig2714.1 |
| 52 | 1 | 565873 | 184 | 64634 | 7 | 7 | 7 | 7 | 0.59 | >jgi\|Phyca11\|565873\|estExt2_Genewise1.C_PHYCAscaffold_190295 |
| 53 | 1 | 6173 | 183 | 109251 | 11 | 11 | 9 | 9 | 0.42 | >jgi\|Phyca11\|6173\|fgenesh1_pm.PHYCAscaffold_10_#_20 |
| 35 | 2 | 530299 | 178 | 142871 | 8 | 8 | 6 | 6 | 0.23 | >jgi\|Phyca11\|530299\|estExt2_fgenesh1_pm.C_PHYCAscaffold_610001 |
| 35 | 3 | 557455 | 173 | 149278 | 8 | 8 | 6 | 6 | 0.22 | >jgi\|Phyca11\|557455\|estExt2_Genewise1Plus.C_PHYCAscaffold_1170012 |
| 54 | 1 | 98712 | 170 | 60979 | 12 | 12 | 10 | 10 | 1.01 | >jgi\|Phyca11\|98712\|e_gw1.3.703.1 |
| 55 | 1 | 503123 | 170 | 44559 | 12 | 12 | 10 | 10 | 1.86 | >jgi\|Phyca11\|503123\|fgenesh2_kg.PHYCAscaffold_2_#_239_#_Contig21.3 |
| 56 | 1 | 502783 | 169 | 83473 | 8 | 8 | 7 | 7 | 0.43 | >jgi\|Phyca11\|502783\|fgenesh2_kg.PHYCAscaffold_1_#_243_#_Contig762.1 |
| 40 | 2 | 70742 | 168 | 19036 | 6 | 6 | 5 | 5 | 2.02 | >jgi\|Phyca11\|70742\|gw1.13.784.1 |
| 57 | 1 | 503983 | 168 | 63243 | 9 | 9 | 7 | 7 | 0.6 | >jgi\|Phyca11\|503983\|fgenesh2_kg.PHYCAscaffold_5_#_147_#_4099435:1 |
| 58 | 1 | 574695 | 165 | 66331 | 9 | 9 | 7 | 7 | 0.57 | >jgi\|Phyca11\|574695\|estExt2_Genewise1.C_PHYCAscaffold_650023 |
| 59 | 1 | 534535 | 164 | 130373 | 11 | 11 | 10 | 10 | 0.39 | >jgi\|Phyca11\|534535\|estExt2_fgenesh1_pg.C_PHYCAscaffold_250008 |
| 35 | 4 | 504139 | 158 | 154076 | 6 | 6 | 4 | 4 | 0.15 | >jgi\|Phyca11\|504139\|fgenesh2_kg.PHYCAscaffold_6_#_56_#_Contig215.1 |
| 60 | 1 |  | 158 | 25078 | 10 | 10 | 2 | 2 | 0.4 | SWISS-PROT:P00761\|TRYP_PIG Trypsin - Sus scrofa (Pig). |
| 61 | 1 | 507331 | 158 | 29680 | 9 | 9 | 7 | 7 | 1.71 | >jgi\|Phyca11\|507331\|fgenesh2_kg.PHYCAscaffold_27_#_5_#_gi\|189084902\|gb\|BT032433.1\| |
| 62 | 1 | 510618 | 156 | 108675 | 11 | 11 | 10 | 10 | 0.54 | >jgi\|Phyca11\|510618\|fgenesh2_kg.PHYCAscaffold_64_#_11_#_Contig39.1 |
| 63 | 1 | 504155 | 154 | 20467 | 7 | 7 | 5 | 5 | 1.8 | >jgi\|Phyca11\|504155\|fgenesh2_kg.PHYCAscaffold_6_#_72_#_4101945:207 |
| 64 | 1 | 508065 | 151 | 61901 | 5 | 5 | 5 | 5 | 0.41 | >jgi\|Phyca11\|508065\|fgenesh2_kg.PHYCAscaffold_32_#_24_#_Contig137.1 |
| 65 | 1 | 132086 | 150 | 42182 | 7 | 7 | 6 | 6 | 0.83 | >jgi\|Phyca11\|132086\|e_gw1.132.17.1 |
| 66 | 1 | 569273 | 147 | 59485 | 8 | 8 | 8 | 8 | 0.77 | >jgi\|Phyca11\|569273\|estExt2_Genewise1.C_PHYCAscaffold_320071 |
| 67 | 1 | 565179 | 143 | 57244 | 8 | 8 | 7 | 7 | 0.68 | >jgi\|Phyca11\|565179\|estExt2_Genewise1.C_PHYCAscaffold_170294 |
| 68 | 1 | 536934 | 141 | 109579 | 6 | 6 | 5 | 5 | 0.22 | >jgi\|Phyca11\|536934\|estExt2_fgenesh1_pg.C_PHYCAscaffold_670026 |
| 69 | 1 | 503431 | 139 | 62538 | 7 | 7 | 6 | 6 | 0.51 | >jgi\|Phyca11\|503431\|fgenesh2_kg.PHYCAscaffold_3_#_296_#_4097660:2 |
| 70 | 1 | 506742 | 139 | 79728 | 8 | 8 | 6 | 6 | 0.38 | >jgi\|Phyca11\|506742\|fgenesh2_kg.PHYCAscaffold_21_#_114_#_Contig1308.1 |
| 71 | 1 | 563206 | 138 | 77231 | 5 | 5 | 5 | 5 | 0.32 | >jgi\|Phyca11\|563206\|estExt2_Genewise1.C_PHYCAscaffold_110375 |
| 72 | 1 | 566774 | 137 | 51922 | 7 | 7 | 6 | 6 | 0.63 | >jgi\|Phyca11\|566774\|estExt2_Genewise1.C_PHYCAscaffold_220429 |
| 65 | 2 |  | 134 | 42052 | 5 | 5 | 4 | 4 | 0.5 | SWISS-PROT:P60712 (Bos taurus) Actin, cytoplasmic 1 |
| 73 | 1 | 34163 | 132 | 90948 | 9 | 9 | 7 | 7 | 0.39 | >jgi\|Phyca11\|34163\|gw1.2.21.1 |
| 74 | 1 | 549506 | 132 | 69618 | 5 | 5 | 4 | 4 | 0.28 | >jgi\|Phyca11\|549506\|estExt2_Genewise1Plus.C_PHYCAscaffold_330019 |
| 75 | 1 | 544983 | 131 | 32876 | 3 | 3 | 2 | 2 | 0.29 | >jgi\|Phyca11\|544983\|estExt2_Genewise1Plus.C_PHYCAscaffold_160492 |
| 76 | 1 | 508289 | 130 | 17173 | 4 | 4 | 4 | 4 | 1.64 | >jgi\|Phyca11\|508289\|fgenesh2_kg.PHYCAscaffold_33_#_148_#_4101647:5 |
| 77 | 1 | 503727 | 129 | 64500 | 6 | 6 | 6 | 6 | 0.49 | >jgi\|Phyca11\|503727\|fgenesh2_kg.PHYCAscaffold_4_#_266_#_Contig7.1 |
| 78 | 1 | 510571 | 127 | 124747 | 7 | 7 | 6 | 6 | 0.23 | >jgi\|Phyca11\|510571\|fgenesh2_kg.PHYCAscaffold_63_#_10_#_gi\|189084688\|gb\|BT032204.1\| |
| 79 | 1 | 510504 | 127 | 38812 | 5 | 5 | 2 | 2 | 0.24 | >jgi\|Phyca11\|510504\|fgenesh2_kg.PHYCAscaffold_61_#_37_#_4101945:41 |
| 43 | 2 | 104335 | 126 | 71675 | 10 | 10 | 8 | 8 | 0.61 | >jgi\|Phyca11\|104335\|e_gw1.9.161.1 |
| 52 | 2 | 110703 | 126 | 66887 | 8 | 8 | 5 | 5 | 0.47 | >jgi\|Phyca11\|110703\|e_gw1.19.482.1 |
| 80 | 1 | 9669 | 126 | 23181 | 4 | 4 | 3 | 3 | 0.72 | >jgi\|Phyca11\|9669\|fgenesh1_pm.PHYCAscaffold_40_#_57 |
| 81 | 1 | 538324 | 125 | 80360 | 5 | 5 | 5 | 5 | 0.3 | >jgi\|Phyca11\|538324\|estExt2_Genewise1Plus.C_PHYCAscaffold_10911 |
| 82 | 1 | 577084 | 123 | 61665 | 5 | 5 | 5 | 5 | 0.41 | >jgi\|Phyca11\|577084\|estExt2_Genewise1.C_PHYCAscaffold_1020014 |
| 83 | 1 | 21374 | 122 | 16113 | 5 | 5 | 3 | 3 | 1.18 | >jgi\|Phyca11\|21374\|fgenesh1_pg.PHYCAscaffold_93_#_8 |
| 84 | 1 | 575784 | 116 | 65395 | 6 | 6 | 5 | 5 | 0.39 | >jgi\|Phyca11\|575784\|estExt2_Genewise1.C_PHYCAscaffold_790031 |
| 85 | 1 | 565192 | 114 | 58334 | 8 | 8 | 7 | 7 | 0.67 | >jgi\|Phyca11\|565192\|estExt2_Genewise1.C_PHYCAscaffold_170319 |
| 86 | 1 | 553954 | 113 | 125096 | 7 | 7 | 6 | 6 | 0.23 | >jgi\|Phyca11\|553954\|estExt2_Genewise1Plus.C_PHYCAscaffold_580009 |
| 87 | 1 | 503898 | 112 | 14775 | 5 | 5 | 4 | 4 | 2.09 | >jgi\|Phyca11\|503898\|fgenesh2_kg.PHYCAscaffold_5_#_62_#_gi\|189083883\|gb\|BT031399.1\| |
| 88 | 1 | 20647 | 112 | 32057 | 6 | 6 | 5 | 5 | 0.93 | >jgi\|Phyca11\|20647\|fgenesh1_pg.PHYCAscaffold_70_#_2 |
| 89 | 1 | 527152 | 108 | 69652 | 7 | 7 | 7 | 7 | 0.54 | >jgi\|Phyca11\|527152\|estExt2_fgenesh1_pm.C_PHYCAscaffold_170020 |
| 90 | 1 | 502852 | 107 | 23768 | 3 | 3 | 3 | 3 | 0.7 | >jgi\|Phyca11\|502852\|fgenesh2_kg.PHYCAscaffold_1_#_312_#_Contig157.1 |
| 91 | 1 | 15447 | 104 | 248333 | 7 | 7 | 7 | 7 | 0.13 | >jgi\|Phyca11\|15447\|fgenesh1_pg.PHYCAscaffold_13_#_135 |
| 92 | 1 | 544083 | 104 | 62563 | 4 | 4 | 3 | 3 | 0.23 | >jgi\|Phyca11\|544083\|estExt2_Genewise1Plus.C_PHYCAscaffold_130608 |
| 30 | 2 | 506056 | 103 | 94357 | 2 | 2 | 2 | 2 | 0.09 | >jgi\|Phyca11\|506056\|fgenesh2_kg.PHYCAscaffold_17_#_141_#_gi\|189084230\|gb\|BT031746.1\| |
| 93 | 1 | 503602 | 103 | 83544 | 7 | 7 | 7 | 7 | 0.43 | >jgi\|Phyca11\|503602\|fgenesh2_kg.PHYCAscaffold_4_#_141_#_gi\|189084094\|gb\|BT031610.1\| |
| 94 | 1 | 507901 | 103 | 42076 | 4 | 4 | 3 | 3 | 0.35 | >jgi\|Phyca11\|507901\|fgenesh2_kg.PHYCAscaffold_30_#_170_#_Contig686.1 |
| 95 | 1 | 533987 | 103 | 136648 | 9 | 9 | 9 | 9 | 0.33 | >jgi\|Phyca11\|533987\|estExt2_fgenesh1_pg.C_PHYCAscaffold_190079 |
| 96 | 1 | 504262 | 103 | 89471 | 6 | 6 | 6 | 6 | 0.33 | >jgi\|Phyca11\|504262\|fgenesh2_kg.PHYCAscaffold_7_#_14_#_4101715:1 |
| 97 | 1 | 508572 | 103 | 73583 | 5 | 5 | 5 | 5 | 0.34 | >jgi\|Phyca11\|508572\|fgenesh2_kg.PHYCAscaffold_36_#_25_#_gi\|189084323\|gb\|BT031839.1\| |
| 98 | 1 | 577748 | 98 | 59749 | 4 | 4 | 3 | 3 | 0.24 | >jgi\|Phyca11\|577748\|estExt2_Genewise1.C_PHYCAscaffold_3510003 |
| 99 | 1 | 568110 | 96 | 74431 | 7 | 7 | 6 | 6 | 0.41 | >jgi\|Phyca11\|568110\|estExt2_Genewise1.C_PHYCAscaffold_270349 |
| 100 | 1 | 507760 | 95 | 27768 | 2 | 2 | 2 | 2 | 0.36 | >jgi\|Phyca11\|507760\|fgenesh2_kg.PHYCAscaffold_30_#_29_#_Contig16.1 |
| 101 | 1 | 504348 | 95 | 29796 | 4 | 4 | 4 | 4 | 0.77 | >jgi\|Phyca11\|504348\|fgenesh2_kg.PHYCAscaffold_7_#_100_#_gi\|189084964\|gb\|BT032496.1\| |
| 102 | 1 | 117520 | 94 | 52656 | 4 | 4 | 4 | 4 | 0.38 | >jgi\|Phyca11\|117520\|e_gw1.33.435.1 |
| 103 | 1 | 510736 | 94 | 65119 | 4 | 4 | 4 | 4 | 0.3 | >jgi\|Phyca11\|510736\|fgenesh2_kg.PHYCAscaffold_66_#_10_#_4101945:56 |
| 104 | 1 | 574456 | 93 | 85938 | 8 | 8 | 8 | 8 | 0.49 | >jgi\|Phyca11\|574456\|estExt2_Genewise1.C_PHYCAscaffold_620176 |
| 105 | 1 | 562791 | 93 | 62172 | 6 | 6 | 4 | 4 | 0.32 | >jgi\|Phyca11\|562791\|estExt2_Genewise1.C_PHYCAscaffold_100242 |
| 106 | 1 | 21290 | 92 | 151885 | 7 | 7 | 7 | 7 | 0.22 | >jgi\|Phyca11\|21290\|fgenesh1_pg.PHYCAscaffold_89_#_4 |
| 107 | 1 | 512063 | 91 | 74585 | 5 | 5 | 5 | 5 | 0.33 | >jgi\|Phyca11\|512063\|fgenesh2_kg.PHYCAscaffold_121_#_12_#_gi\|189084018\|gb\|BT031534.1\| |
| 108 | 1 | 531039 | 90 | 101658 | 6 | 6 | 5 | 5 | 0.23 | >jgi\|Phyca11\|531039\|estExt2_fgenesh1_pm.C_PHYCAscaffold_910003 |
| 109 | 1 | 509300 | 90 | 80654 | 4 | 4 | 4 | 4 | 0.24 | >jgi\|Phyca11\|509300\|fgenesh2_kg.PHYCAscaffold_43_#_98_#_Contig479.1 |
| 110 | 1 | 540959 | 89 | 21538 | 6 | 6 | 4 | 4 | 1.18 | >jgi\|Phyca11\|540959\|estExt2_Genewise1Plus.C_PHYCAscaffold_50907 |
| 111 | 1 | 123621 | 89 | 52506 | 6 | 6 | 6 | 6 | 0.63 | >jgi\|Phyca11\|123621\|e_gw1.51.113.1 |
| 112 | 1 | 511305 | 88 | 88501 | 5 | 5 | 5 | 5 | 0.27 | >jgi\|Phyca11\|511305\|fgenesh2_kg.PHYCAscaffold_80_#_49_#_4098993:3 |
| 113 | 1 | 503425 | 86 | 29978 | 6 | 6 | 5 | 5 | 1.02 | >jgi\|Phyca11\|503425\|fgenesh2_kg.PHYCAscaffold_3_#_290_#_4101945:196 |
| 35 | 5 | 127018 | 85 | 108376 | 5 | 5 | 5 | 5 | 0.22 | >jgi\|Phyca11\|127018\|e_gw1.66.38.1 |
| 114 | 1 | 527083 | 84 | 43530 | 7 | 7 | 5 | 5 | 0.63 | >jgi\|Phyca11\|527083\|estExt2_fgenesh1_pm.C_PHYCAscaffold_150137 |
| 115 | 1 | 511466 | 83 | 78712 | 4 | 4 | 4 | 4 | 0.24 | >jgi\|Phyca11\|511466\|fgenesh2_kg.PHYCAscaffold_86_#_3_#_gi\|189084813\|gb\|BT032344.1\| |
| 116 | 1 |  | 83 | 71244 | 5 | 5 | 5 | 5 | 0.35 | SWISS-PROT:P02769 (Bos taurus) Bovine serum albumin precursor |
| 117 | 1 | 504369 | 82 | 76494 | 4 | 4 | 3 | 3 | 0.18 | >jgi\|Phyca11\|504369\|fgenesh2_kg.PHYCAscaffold_7_#_121_#_gi\|189084575\|gb\|BT032091.1\| |
| 118 | 1 | 504053 | 81 | 71164 | 3 | 3 | 3 | 3 | 0.2 | >jgi\|Phyca11\|504053\|fgenesh2_kg.PHYCAscaffold_5_#_217_#_Contig796.1 |
| 119 | 1 | 525553 | 80 | 64751 | 6 | 6 | 6 | 6 | 0.48 | >jgi\|Phyca11\|525553\|estExt2_fgenesh1_pm.C_PHYCAscaffold_40090 |
| 120 | 1 | 118185 | 79 | 222850 | 3 | 3 | 3 | 3 | 0.06 | >jgi\|Phyca11\|118185\|e_gw1.35.65.1 |
| 121 | 1 | 111740 | 79 | 90538 | 7 | 7 | 7 | 7 | 0.39 | >jgi\|Phyca11\|111740\|e_gw1.20.14.1 |
| 122 | 1 | 118996 | 79 | 76653 | 3 | 3 | 3 | 3 | 0.18 | >jgi\|Phyca11\|118996\|e_gw1.37.459.1 |
| 123 | 1 | 542260 | 78 | 89224 | 4 | 4 | 4 | 4 | 0.21 | >jgi\|Phyca11\|542260\|estExt2_Genewise1Plus.C_PHYCAscaffold_90046 |
| 124 | 1 | 530787 | 78 | 60685 | 2 | 2 | 2 | 2 | 0.15 | >jgi\|Phyca11\|530787\|estExt2_fgenesh1_pm.C_PHYCAscaffold_780004 |
| 125 | 1 | 510359 | 77 | 70048 | 5 | 5 | 5 | 5 | 0.36 | >jgi\|Phyca11\|510359\|fgenesh2_kg.PHYCAscaffold_58_#_72_#_Contig2242.1 |
| 126 | 1 | 503358 | 77 | 51975 | 3 | 3 | 3 | 3 | 0.28 | >jgi\|Phyca11\|503358\|fgenesh2_kg.PHYCAscaffold_3_#_223_#_Contig173.1 |
| 127 | 1 | 74521 | 76 | 64501 | 4 | 4 | 3 | 3 | 0.22 | >jgi\|Phyca11\|74521\|gw1.15.657.1 |
| 128 | 1 | 504469 | 76 | 23161 | 4 | 4 | 4 | 4 | 1.07 | >jgi\|Phyca11\|504469\|fgenesh2_kg.PHYCAscaffold_8_#_38_#_4101945:176 |
| 129 | 1 | 504594 | 76 | 48837 | 3 | 3 | 3 | 3 | 0.3 | >jgi\|Phyca11\|504594\|fgenesh2_kg.PHYCAscaffold_8_#_163_#_Contig1207.1 |
| 130 | 1 | 542625 | 75 | 62190 | 2 | 2 | 2 | 2 | 0.15 | >jgi\|Phyca11\|542625\|estExt2_Genewise1Plus.C_PHYCAscaffold_90614 |
| 131 | 1 | 504894 | 75 | 121029 | 6 | 6 | 6 | 6 | 0.24 | >jgi\|Phyca11\|504894\|fgenesh2_kg.PHYCAscaffold_10_#_54_#_Contig182.1 |
| 132 | 1 | 109154 | 75 | 22119 | 2 | 2 | 2 | 2 | 0.46 | >jgi\|Phyca11\|109154\|e_gw1.16.590.1 |
| 117 | 2 | 532433 | 74 | 85584 | 5 | 5 | 3 | 3 | 0.16 | >jgi\|Phyca11\|532433\|estExt2_fgenesh1_pg.C_PHYCAscaffold_50170 |
| 133 | 1 | 504612 | 74 | 117382 | 5 | 5 | 5 | 5 | 0.2 | >jgi\|Phyca11\|504612\|fgenesh2_kg.PHYCAscaffold_9_#_3_#_Contig947.1 |
| 134 | 1 | 124610 | 74 | 89043 | 6 | 6 | 6 | 6 | 0.33 | >jgi\|Phyca11\|124610\|e_gw1.54.84.1 |
| 135 | 1 |  | 74 | 80499 | 1 | 1 | 1 | 1 | 0.05 |  |
| 136 | 1 | 503447 | 73 | 57286 | 2 | 2 | 2 | 2 | 0.16 | >jgi\|Phyca11\|503447\|fgenesh2_kg.PHYCAscaffold_3_#_312_#_4097098:2 |
| 137 | 1 | 507472 | 73 | 24177 | 4 | 4 | 4 | 4 | 1.01 | >jgi\|Phyca11\|507472\|fgenesh2_kg.PHYCAscaffold_27_#_146_#_Contig84.1 |
| 138 | 1 | 122536 | 73 | 85692 | 2 | 2 | 2 | 2 | 0.1 | >jgi\|Phyca11\|122536\|e_gw1.48.64.1 |
| 139 | 1 | 538943 | 73 | 167258 | 5 | 5 | 5 | 5 | 0.14 | >jgi\|Phyca11\|538943\|estExt2_Genewise1Plus.C_PHYCAscaffold_20665 |
| 140 | 1 | 563829 | 73 | 32177 | 3 | 3 | 2 | 2 | 0.3 | >jgi\|Phyca11\|563829\|estExt2_Genewise1.C_PHYCAscaffold_130199 |
| 141 | 1 | 509051 | 72 | 69322 | 7 | 7 | 6 | 6 | 0.45 | >jgi\|Phyca11\|509051\|fgenesh2_kg.PHYCAscaffold_41_#_47_#_Contig254.1 |
| 142 | 1 | 19592 | 72 | 60727 | 4 | 4 | 4 | 4 | 0.32 | >jgi\|Phyca11\|19592\|fgenesh1_pg.PHYCAscaffold_50_#_14 |
| 143 | 1 | 504276 | 72 | 76273 | 2 | 2 | 2 | 2 | 0.12 | >jgi\|Phyca11\|504276\|fgenesh2_kg.PHYCAscaffold_7_#_28_#_Contig229.1 |
| 43 | 3 | 510696 | 71 | 72798 | 5 | 5 | 5 | 5 | 0.34 | >jgi\|Phyca11\|510696\|fgenesh2_kg.PHYCAscaffold_65_#_36_#_4098202:1 |
| 144 | 1 | 122065 | 71 | 53767 | 5 | 5 | 4 | 4 | 0.37 | >jgi\|Phyca11\|122065\|e_gw1.47.14.1 |
| 139 | 2 | 526928 | 70 | 161520 | 3 | 3 | 3 | 3 | 0.08 | >jgi\|Phyca11\|526928\|estExt2_fgenesh1_pm.C_PHYCAscaffold_140061 |
| 145 | 1 | 559116 | 70 | 84985 | 5 | 5 | 5 | 5 | 0.29 | >jgi\|Phyca11\|559116\|estExt2_Genewise1.C_PHYCAscaffold_21127 |
| 146 | 1 | 502943 | 70 | 16614 | 2 | 2 | 2 | 2 | 0.65 | >jgi\|Phyca11\|502943\|fgenesh2_kg.PHYCAscaffold_2_#_59_#_gi\|189083898\|gb\|BT031414.1\| |
| 147 | 1 | 531982 | 69 | 130522 | 2 | 2 | 2 | 2 | 0.07 | >jgi\|Phyca11\|531982\|estExt2_fgenesh1_pg.C_PHYCAscaffold_30148 |
| 148 | 1 | 560995 | 69 | 86993 | 3 | 3 | 3 | 3 | 0.16 | >jgi\|Phyca11\|560995\|estExt2_Genewise1.C_PHYCAscaffold_51129 |
| 149 | 1 | 508801 | 69 | 20272 | 4 | 4 | 2 | 2 | 0.51 | >jgi\|Phyca11\|508801\|fgenesh2_kg.PHYCAscaffold_38_#_53_#_Contig95.1 |
| 150 | 1 | 506902 | 69 | 27776 | 4 | 4 | 4 | 4 | 0.84 | >jgi\|Phyca11\|506902\|fgenesh2_kg.PHYCAscaffold_23_#_33_#_gi\|189084893\|gb\|BT032424.1\| |
| 151 | 1 | 542873 | 68 | 55889 | 4 | 4 | 3 | 3 | 0.26 | >jgi\|Phyca11\|542873\|estExt2_Genewise1Plus.C_PHYCAscaffold_100329 |
| 152 | 1 | 535097 | 67 | 77097 | 5 | 5 | 5 | 5 | 0.32 | >jgi\|Phyca11\|535097\|estExt2_fgenesh1_pg.C_PHYCAscaffold_310096 |
| 153 | 1 | 572227 | 66 | 57321 | 2 | 2 | 2 | 2 | 0.16 | >jgi\|Phyca11\|572227\|estExt2_Genewise1.C_PHYCAscaffold_470089 |
| 154 | 1 | 561150 | 66 | 65782 | 2 | 2 | 2 | 2 | 0.14 | >jgi\|Phyca11\|561150\|estExt2_Genewise1.C_PHYCAscaffold_60312 |
| 155 | 1 | 6748 | 65 | 140772 | 3 | 3 | 3 | 3 | 0.1 | >jgi\|Phyca11\|6748\|fgenesh1_pm.PHYCAscaffold_14_#_52 |
| 156 | 1 | 574490 | 65 | 129810 | 7 | 7 | 6 | 6 | 0.22 | >jgi\|Phyca11\|574490\|estExt2_Genewise1.C_PHYCAscaffold_630028 |
| 157 | 1 | 533489 | 64 | 176235 | 2 | 2 | 2 | 2 | 0.05 | >jgi\|Phyca11\|533489\|estExt2_fgenesh1_pg.C_PHYCAscaffold_140069 |
| 158 | 1 | 509308 | 64 | 35212 | 2 | 2 | 2 | 2 | 0.27 | >jgi\|Phyca11\|509308\|fgenesh2_kg.PHYCAscaffold_43_#_106_#_Contig105.1 |
| 159 | 1 |  | 64 | 249296 | 1 | 1 | 1 | 1 | 0.02 | SWISS-PROT:Q5D862 Tax_Id=9606 Gene_Symbol=FLG2 Filaggrin-2 |
| 160 | 1 | 507516 | 63 | 34484 | 1 | 1 | 1 | 1 | 0.13 | >jgi\|Phyca11\|507516\|fgenesh2_kg.PHYCAscaffold_28_#_23_#_Contig24.1 |
| 161 | 1 | 525308 | 63 | 73526 | 6 | 6 | 5 | 5 | 0.34 | >jgi\|Phyca11\|525308\|estExt2_fgenesh1_pm.C_PHYCAscaffold_30053 |
| 162 | 1 | 21359 | 63 | 63609 | 1 | 1 | 1 | 1 | 0.07 | >jgi\|Phyca11\|21359\|fgenesh1_pg.PHYCAscaffold_92_#_14 |
| 163 | 1 | 571594 | 63 | 59774 | 6 | 6 | 5 | 5 | 0.43 | >jgi\|Phyca11\|571594\|estExt2_Genewise1.C_PHYCAscaffold_430074 |
| 164 | 1 | 561936 | 62 | 61077 | 5 | 5 | 5 | 5 | 0.42 | >jgi\|Phyca11\|561936\|estExt2_Genewise1.C_PHYCAscaffold_80266 |
| 165 | 1 | 558277 | 62 | 130749 | 3 | 3 | 3 | 3 | 0.1 | >jgi\|Phyca11\|558277\|estExt2_Genewise1.C_PHYCAscaffold_10916 |
| 166 | 1 | 527927 | 62 | 74101 | 4 | 4 | 4 | 4 | 0.26 | >jgi\|Phyca11\|527927\|estExt2_fgenesh1_pm.C_PHYCAscaffold_240054 |
| 167 | 1 | 510361 | 61 | 140057 | 4 | 4 | 4 | 4 | 0.13 | >jgi\|Phyca11\|510361\|fgenesh2_kg.PHYCAscaffold_58_#_74_#_gi\|189084453\|gb\|BT031969.1\| |
| 168 | 1 | 510582 | 60 | 52105 | 2 | 2 | 2 | 2 | 0.18 | >jgi\|Phyca11\|510582\|fgenesh2_kg.PHYCAscaffold_63_#_21_#_Contig2224.1 |
| 153 | 2 | 122092 | 59 | 47524 | 2 | 2 | 2 | 2 | 0.2 | >jgi\|Phyca11\|122092\|e_gw1.47.100.1 |
| 169 | 1 | 542884 | 58 | 97887 | 3 | 3 | 3 | 3 | 0.14 | >jgi\|Phyca11\|542884\|estExt2_Genewise1Plus.C_PHYCAscaffold_100379 |
| 170 | 1 | 127542 | 58 | 56132 | 2 | 2 | 2 | 2 | 0.16 | >jgi\|Phyca11\|127542\|e_gw1.70.87.1 |
| 52 | 3 | 506373 | 57 | 65226 | 2 | 2 | 2 | 2 | 0.14 | >jgi\|Phyca11\|506373\|fgenesh2_kg.PHYCAscaffold_19_#_79_#_Contig441.1 |
| 171 | 1 | 530845 | 57 | 61153 | 2 | 2 | 2 | 2 | 0.15 | >jgi\|Phyca11\|530845\|estExt2_fgenesh1_pm.C_PHYCAscaffold_800021 |
| 172 | 1 | 558677 | 57 | 99339 | 2 | 2 | 2 | 2 | 0.09 | >jgi\|Phyca11\|558677\|estExt2_Genewise1.C_PHYCAscaffold_20357 |
| 173 | 1 | 509239 | 57 | 61394 | 6 | 6 | 5 | 5 | 0.41 | >jgi\|Phyca11\|509239\|fgenesh2_kg.PHYCAscaffold_43_#_37_#_4101945:59 |
| 174 | 1 | 100378 | 56 | 58668 | 2 | 2 | 2 | 2 | 0.16 | >jgi\|Phyca11\|100378\|e_gw1.4.499.1 |
| 175 | 1 | 576097 | 55 | 49574 | 3 | 3 | 3 | 3 | 0.29 | >jgi\|Phyca11\|576097\|estExt2_Genewise1.C_PHYCAscaffold_840019 |
| 176 | 1 | 505375 | 55 | 17847 | 4 | 4 | 4 | 4 | 1.55 | >jgi\|Phyca11\|505375\|fgenesh2_kg.PHYCAscaffold_13_#_25_#_4098936:2 |
| 177 | 1 | 545771 | 55 | 116345 | 3 | 3 | 3 | 3 | 0.12 | >jgi\|Phyca11\|545771\|estExt2_Genewise1Plus.C_PHYCAscaffold_190083 |
| 178 | 1 | 506502 | 55 | 105481 | 3 | 3 | 3 | 3 | 0.13 | >jgi\|Phyca11\|506502\|fgenesh2_kg.PHYCAscaffold_20_#_48_#_gi\|189084416\|gb\|BT031932.1\| |
| 179 | 1 | 505188 | 55 | 27120 | 2 | 2 | 2 | 2 | 0.37 | >jgi\|Phyca11\|505188\|fgenesh2_kg.PHYCAscaffold_11_#_134_#_Contig42.1 |
| 180 | 1 | 525091 | 54 | 19610 | 1 | 1 | 1 | 1 | 0.24 | >jgi\|Phyca11\|525091\|estExt2_fgenesh1_pm.C_PHYCAscaffold_20022 |
| 181 | 1 | 508721 | 54 | 90477 | 3 | 3 | 3 | 3 | 0.15 | >jgi\|Phyca11\|508721\|fgenesh2_kg.PHYCAscaffold_37_#_96_#_4098316:1 |
| 182 | 1 | 506596 | 54 | 63682 | 4 | 4 | 4 | 4 | 0.31 | >jgi\|Phyca11\|506596\|fgenesh2_kg.PHYCAscaffold_20_#_142_#_gi\|189084101\|gb\|BT031617.1\| |
| 183 | 1 | 502973 | 54 | 22064 | 3 | 3 | 3 | 3 | 0.77 | >jgi\|Phyca11\|502973\|fgenesh2_kg.PHYCAscaffold_2_#_89_#_4101945:287 |
| 184 | 1 | 502708 | 53 | 26043 | 2 | 2 | 2 | 2 | 0.38 | >jgi\|Phyca11\|502708\|fgenesh2_kg.PHYCAscaffold_1_#_168_#_Contig881.1 |
| 35 | 6 | 20570 | 52 | 66384 | 4 | 4 | 4 | 4 | 0.29 | >jgi\|Phyca11\|20570\|fgenesh1_pg.PHYCAscaffold_66_#_46 |
| 49 | 2 | 571405 | 52 | 163648 | 2 | 2 | 2 | 2 | 0.05 | >jgi\|Phyca11\|571405\|estExt2_Genewise1.C_PHYCAscaffold_420119 |
| 185 | 1 | 16638 | 52 | 29644 | 1 | 1 | 1 | 1 | 0.15 | >jgi\|Phyca11\|16638\|fgenesh1_pg.PHYCAscaffold_21_#_55 |
| 186 | 1 | 575101 | 52 | 84264 | 3 | 3 | 3 | 3 | 0.16 | >jgi\|Phyca11\|575101\|estExt2_Genewise1.C_PHYCAscaffold_700114 |
| 187 | 1 | 573771 | 52 | 91543 | 2 | 2 | 1 | 1 | 0.05 | >jgi\|Phyca11\|573771\|estExt2_Genewise1.C_PHYCAscaffold_550212 |
| 188 | 1 | 576285 | 52 | 61508 | 3 | 3 | 3 | 3 | 0.23 | >jgi\|Phyca11\|576285\|estExt2_Genewise1.C_PHYCAscaffold_860075 |
| 189 | 1 | 571228 | 52 | 47998 | 2 | 2 | 2 | 2 | 0.19 | >jgi\|Phyca11\|571228\|estExt2_Genewise1.C_PHYCAscaffold_410155 |
| 190 | 1 | 535506 | 52 | 74784 | 1 | 1 | 1 | 1 | 0.06 | >jgi\|Phyca11\|535506\|estExt2_fgenesh1_pg.C_PHYCAscaffold_370049 |
| 126 | 2 | 503356 | 51 | 49536 | 2 | 2 | 2 | 2 | 0.19 | >jgi\|Phyca11\|503356\|fgenesh2_kg.PHYCAscaffold_3_#_221_#_4099680:2 |
| 191 | 1 | 508077 | 51 | 152340 | 3 | 3 | 3 | 3 | 0.09 | >jgi\|Phyca11\|508077\|fgenesh2_kg.PHYCAscaffold_32_#_36_#_Contig2846.1 |
| 192 | 1 | 106526 | 51 | 99721 | 1 | 1 | 1 | 1 | 0.04 | >jgi\|Phyca11\|106526\|e_gw1.12.724.1 |
| 193 | 1 | 506178 | 51 | 86004 | 1 | 1 | 1 | 1 | 0.05 | >jgi\|Phyca11\|506178\|fgenesh2_kg.PHYCAscaffold_18_#_51_#_Contig938.1 |
| 194 | 1 | 538776 | 51 | 17625 | 1 | 1 | 1 | 1 | 0.27 | >jgi\|Phyca11\|538776\|estExt2_Genewise1Plus.C_PHYCAscaffold_20418 |
| 195 | 1 | 20438 | 51 | 80714 | 2 | 2 | 2 | 2 | 0.11 | >jgi\|Phyca11\|20438\|fgenesh1_pg.PHYCAscaffold_64_#_24 |
| 196 | 1 | 535286 | 50 | 57740 | 1 | 1 | 1 | 1 | 0.08 | >jgi\|Phyca11\|535286\|estExt2_fgenesh1_pg.C_PHYCAscaffold_330120 |
| 197 | 1 | 535142 | 49 | 56284 | 4 | 4 | 4 | 4 | 0.35 | >jgi\|Phyca11\|535142\|estExt2_fgenesh1_pg.C_PHYCAscaffold_320051 |
| 198 | 1 | 120320 | 49 | 124349 | 3 | 3 | 3 | 3 | 0.11 | >jgi\|Phyca11\|120320\|e_gw1.41.116.1 |
| 199 | 1 | 535625 | 49 | 223563 | 2 | 2 | 2 | 2 | 0.04 | >jgi\|Phyca11\|535625\|estExt2_fgenesh1_pg.C_PHYCAscaffold_390039 |
| 200 | 1 | 504590 | 48 | 23475 | 1 | 1 | 1 | 1 | 0.2 | >jgi\|Phyca11\|504590\|fgenesh2_kg.PHYCAscaffold_8_#_159_#_Contig101.1 |
| 201 | 1 | 508983 | 48 | 33906 | 4 | 4 | 3 | 3 | 0.46 | >jgi\|Phyca11\|508983\|fgenesh2_kg.PHYCAscaffold_40_#_72_#_Contig64.1 |
| 202 | 1 | 118234 | 48 | 14132 | 1 | 1 | 1 | 1 | 0.34 | >jgi\|Phyca11\|118234\|e_gw1.35.157.1 |
| 203 | 1 | 6376 | 48 | 90763 | 2 | 2 | 2 | 2 | 0.1 | >jgi\|Phyca11\|6376\|fgenesh1_pm.PHYCAscaffold_11_#_92 |
| 204 | 1 | 545320 | 48 | 95221 | 3 | 3 | 3 | 3 | 0.14 | >jgi\|Phyca11\|545320\|estExt2_Genewise1Plus.C_PHYCAscaffold_170465 |
| 205 | 1 | 506722 | 47 | 127848 | 3 | 3 | 3 | 3 | 0.11 | >jgi\|Phyca11\|506722\|fgenesh2_kg.PHYCAscaffold_21_#_94_#_4096595:1 |
| 206 | 1 | 504052 | 47 | 69526 | 3 | 3 | 2 | 2 | 0.13 | >jgi\|Phyca11\|504052\|fgenesh2_kg.PHYCAscaffold_5_#_216_#_Contig1023.1 |
| 207 | 1 | 503568 | 47 | 49308 | 1 | 1 | 1 | 1 | 0.09 | >jgi\|Phyca11\|503568\|fgenesh2_kg.PHYCAscaffold_4_#_107_#_4097947:1 |
| 208 | 1 | 507518 | 47 | 73710 | 2 | 2 | 2 | 2 | 0.12 | >jgi\|Phyca11\|507518\|fgenesh2_kg.PHYCAscaffold_28_#_25_#_4101945:43 |
| 209 | 1 | 507227 | 47 | 53724 | 2 | 2 | 2 | 2 | 0.17 | >jgi\|Phyca11\|507227\|fgenesh2_kg.PHYCAscaffold_26_#_30_#_Contig197.1 |
| 210 | 1 | 6127 | 46 | 63943 | 3 | 3 | 3 | 3 | 0.22 | >jgi\|Phyca11\|6127\|fgenesh1_pm.PHYCAscaffold_9_#_187 |
| 211 | 1 | 118056 | 46 | 69434 | 3 | 3 | 3 | 3 | 0.2 | >jgi\|Phyca11\|118056\|e_gw1.35.12.1 |
| 212 | 1 | 541200 | 46 | 15076 | 2 | 2 | 2 | 2 | 0.74 | >jgi\|Phyca11\|541200\|estExt2_Genewise1Plus.C_PHYCAscaffold_60305 |
| 213 | 1 | 112758 | 46 | 16983 | 1 | 1 | 1 | 1 | 0.28 | >jgi\|Phyca11\|112758\|e_gw1.22.548.1 |
| 214 | 1 | 56419 | 45 | 41111 | 1 | 1 | 1 | 1 | 0.11 | >jgi\|Phyca11\|56419\|gw1.27.248.1 |
| 215 | 1 | 505505 | 45 | 36287 | 5 | 5 | 4 | 4 | 0.6 | >jgi\|Phyca11\|505505\|fgenesh2_kg.PHYCAscaffold_13_#_155_#_4100209:1 |
| 216 | 1 | 547672 | 44 | 59367 | 2 | 2 | 2 | 2 | 0.15 | >jgi\|Phyca11\|547672\|estExt2_Genewise1Plus.C_PHYCAscaffold_260082 |
| 217 | 1 | 531975 | 44 | 72070 | 2 | 2 | 2 | 2 | 0.13 | >jgi\|Phyca11\|531975\|estExt2_fgenesh1_pg.C_PHYCAscaffold_30139 |
| 218 | 1 | 528052 | 44 | 132419 | 2 | 2 | 2 | 2 | 0.07 | >jgi\|Phyca11\|528052\|estExt2_fgenesh1_pm.C_PHYCAscaffold_260044 |
| 219 | 1 | 510914 | 43 | 17277 | 3 | 3 | 2 | 2 | 0.62 | >jgi\|Phyca11\|510914\|fgenesh2_kg.PHYCAscaffold_71_#_14_#_gi\|189083867\|gb\|BT031383.1\| |
| 220 | 1 | 509055 | 43 | 96457 | 3 | 3 | 3 | 3 | 0.14 | >jgi\|Phyca11\|509055\|fgenesh2_kg.PHYCAscaffold_41_#_51_#_4101815:2 |
| 221 | 1 | 121918 | 43 | 104722 | 1 | 1 | 1 | 1 | 0.04 | >jgi\|Phyca11\|121918\|e_gw1.46.425.1 |
| 222 | 1 | 504277 | 43 | 78116 | 1 | 1 | 1 | 1 | 0.06 | >jgi\|Phyca11\|504277\|fgenesh2_kg.PHYCAscaffold_7_#_29_#_Contig133.1 |
| 223 | 1 | 507368 | 42 | 63627 | 2 | 2 | 2 | 2 | 0.14 | >jgi\|Phyca11\|507368\|fgenesh2_kg.PHYCAscaffold_27_#_42_#_Contig268.1 |
| 224 | 1 | 505013 | 42 | 27838 | 2 | 2 | 2 | 2 | 0.35 | >jgi\|Phyca11\|505013\|fgenesh2_kg.PHYCAscaffold_10_#_173_#_gi\|189084885\|gb\|BT032416.1\| |
| 225 | 1 | 561387 | 42 | 44291 | 1 | 1 | 1 | 1 | 0.1 | >jgi\|Phyca11\|561387\|estExt2_Genewise1.C_PHYCAscaffold_60808 |
| 226 | 1 | 525609 | 42 | 99796 | 1 | 1 | 1 | 1 | 0.04 | >jgi\|Phyca11\|525609\|estExt2_fgenesh1_pm.C_PHYCAscaffold_40150 |
| 227 | 1 | 545598 | 42 | 111192 | 6 | 6 | 6 | 6 | 0.26 | >jgi\|Phyca11\|545598\|estExt2_Genewise1Plus.C_PHYCAscaffold_180327 |
| 228 | 1 | 507425 | 42 | 21112 | 2 | 2 | 2 | 2 | 0.49 | >jgi\|Phyca11\|507425\|fgenesh2_kg.PHYCAscaffold_27_#_99_#_4096835:1 |
| 229 | 1 | 116914 | 42 | 30361 | 3 | 3 | 3 | 3 | 0.52 | >jgi\|Phyca11\|116914\|e_gw1.32.324.1 |
| 230 | 1 | 111011 | 42 | 47792 | 1 | 1 | 1 | 1 | 0.09 | >jgi\|Phyca11\|111011\|e_gw1.19.39.1 |
| 231 | 1 | 511938 | 41 | 46144 | 2 | 2 | 2 | 2 | 0.2 | >jgi\|Phyca11\|511938\|fgenesh2_kg.PHYCAscaffold_105_#_36_#_4097069:3 |
| 232 | 1 | 508441 | 41 | 64542 | 1 | 1 | 1 | 1 | 0.07 | >jgi\|Phyca11\|508441\|fgenesh2_kg.PHYCAscaffold_35_#_24_#_Contig211.1 |
| 233 | 1 | 9769 | 41 | 21549 | 3 | 3 | 3 | 3 | 0.8 | >jgi\|Phyca11\|9769\|fgenesh1_pm.PHYCAscaffold_41_#_63 |
| 234 | 1 |  | 40 | 118096 | 3 | 3 | 3 | 3 | 0.11 |  |
| 235 | 1 | 10281 | 40 | 51491 | 2 | 2 | 2 | 2 | 0.18 | >jgi\|Phyca11\|10281\|fgenesh1_pm.PHYCAscaffold_48_#_18 |
| 236 | 1 | 124163 | 40 | 11399 | 1 | 1 | 1 | 1 | 0.44 | >jgi\|Phyca11\|124163\|e_gw1.53.42.1 |
| 237 | 1 | 507239 | 40 | 71157 | 1 | 1 | 1 | 1 | 0.06 | >jgi\|Phyca11\|507239\|fgenesh2_kg.PHYCAscaffold_26_#_42_#_Contig494.1 |
| 238 | 1 | 116925 | 40 | 68303 | 1 | 1 | 1 | 1 | 0.06 | >jgi\|Phyca11\|116925\|e_gw1.32.161.1 |
| 239 | 1 | 503199 | 39 | 51104 | 3 | 3 | 3 | 3 | 0.28 | >jgi\|Phyca11\|503199\|fgenesh2_kg.PHYCAscaffold_3_#_64_#_gi\|189084616\|gb\|BT032132.1\| |
| 240 | 1 | 575550 | 39 | 29762 | 2 | 2 | 2 | 2 | 0.33 | >jgi\|Phyca11\|575550\|estExt2_Genewise1.C_PHYCAscaffold_750112 |
| 241 | 1 | 554070 | 39 | 135803 | 1 | 1 | 1 | 1 | 0.03 | >jgi\|Phyca11\|554070\|estExt2_Genewise1Plus.C_PHYCAscaffold_580178 |
| 242 | 1 | 507479 | 39 | 44888 | 1 | 1 | 1 | 1 | 0.1 | >jgi\|Phyca11\|507479\|fgenesh2_kg.PHYCAscaffold_27_#_153_#_4098657:2 |
| 243 | 1 | 542607 | 39 | 51338 | 2 | 2 | 2 | 2 | 0.18 | >jgi\|Phyca11\|542607\|estExt2_Genewise1Plus.C_PHYCAscaffold_90588 |
| 244 | 1 | 527698 | 39 | 60965 | 2 | 2 | 2 | 2 | 0.15 | >jgi\|Phyca11\|527698\|estExt2_fgenesh1_pm.C_PHYCAscaffold_220003 |
| 245 | 1 | 507461 | 39 | 62650 | 1 | 1 | 1 | 1 | 0.07 | >jgi\|Phyca11\|507461\|fgenesh2_kg.PHYCAscaffold_27_#_135_#_Contig2698.1 |
| 205 | 2 | 54680 | 38 | 83997 | 2 | 2 | 2 | 2 | 0.11 | >jgi\|Phyca11\|54680\|gw1.21.144.1 |
| 246 | 1 | 505233 | 38 | 20821 | 2 | 2 | 1 | 1 | 0.22 | >jgi\|Phyca11\|505233\|fgenesh2_kg.PHYCAscaffold_12_#_21_#_4100316:9 |
| 247 | 1 | 558170 | 38 | 47229 | 2 | 2 | 2 | 2 | 0.2 | >jgi\|Phyca11\|558170\|estExt2_Genewise1.C_PHYCAscaffold_10741 |
| 248 | 1 | 15064 | 38 | 94252 | 2 | 2 | 1 | 1 | 0.05 | >jgi\|Phyca11\|15064\|fgenesh1_pg.PHYCAscaffold_11_#_50 |
| 249 | 1 | 568901 | 38 | 159351 | 3 | 3 | 3 | 3 | 0.08 | >jgi\|Phyca11\|568901\|estExt2_Genewise1.C_PHYCAscaffold_300383 |
| 250 | 1 | 510725 | 38 | 33122 | 1 | 1 | 1 | 1 | 0.14 | >jgi\|Phyca11\|510725\|fgenesh2_kg.PHYCAscaffold_65_#_65_#_Contig957.1 |
| 251 | 1 | 503542 | 37 | 53948 | 1 | 1 | 1 | 1 | 0.08 | >jgi\|Phyca11\|503542\|fgenesh2_kg.PHYCAscaffold_4_#_81_#_Contig1894.1 |
| 252 | 1 | 541046 | 37 | 17198 | 1 | 1 | 1 | 1 | 0.28 | >jgi\|Phyca11\|541046\|estExt2_Genewise1Plus.C_PHYCAscaffold_51078 |
| 253 | 1 | 5622 | 37 | 193584 | 2 | 2 | 1 | 1 | 0.02 | >jgi\|Phyca11\|5622\|fgenesh1_pm.PHYCAscaffold_6_#_142 |
| 254 | 1 | 503496 | 37 | 41637 | 2 | 2 | 1 | 1 | 0.11 | >jgi\|Phyca11\|503496\|fgenesh2_kg.PHYCAscaffold_4_#_35_#_Contig2825.1 |
| 255 | 1 | 113812 | 37 | 20248 | 2 | 2 | 2 | 2 | 0.52 | >jgi\|Phyca11\|113812\|e_gw1.25.423.1 |
| 256 | 1 | 509497 | 36 | 64598 | 2 | 2 | 2 | 2 | 0.14 | >jgi\|Phyca11\|509497\|fgenesh2_kg.PHYCAscaffold_46_#_82_#_Contig380.1 |
| 257 | 1 | 561666 | 36 | 29051 | 1 | 1 | 1 | 1 | 0.16 | >jgi\|Phyca11\|561666\|estExt2_Genewise1.C_PHYCAscaffold_70606 |
| 258 | 1 | 541961 | 36 | 64086 | 1 | 1 | 1 | 1 | 0.07 | >jgi\|Phyca11\|541961\|estExt2_Genewise1Plus.C_PHYCAscaffold_80181 |
| 259 | 1 | 7637 | 36 | 214844 | 2 | 2 | 2 | 2 | 0.04 | >jgi\|Phyca11\|7637\|fgenesh1_pm.PHYCAscaffold_21_#_10 |
| 260 | 1 | 505702 | 36 | 33335 | 2 | 2 | 2 | 2 | 0.29 | >jgi\|Phyca11\|505702\|fgenesh2_kg.PHYCAscaffold_15_#_9_#_4096891:2 |
| 261 | 1 | 511639 | 36 | 39884 | 1 | 1 | 1 | 1 | 0.11 | >jgi\|Phyca11\|511639\|fgenesh2_kg.PHYCAscaffold_93_#_10_#_Contig2592.1 |
| 262 | 1 | 553648 | 36 | 54839 | 1 | 1 | 1 | 1 | 0.08 | >jgi\|Phyca11\|553648\|estExt2_Genewise1Plus.C_PHYCAscaffold_540171 |
| 263 | 1 | 504902 | 36 | 24490 | 1 | 1 | 1 | 1 | 0.19 | >jgi\|Phyca11\|504902\|fgenesh2_kg.PHYCAscaffold_10_#_62_#_4096919:1 |
| 264 | 1 | 564153 | 35 | 41377 | 1 | 1 | 1 | 1 | 0.11 | >jgi\|Phyca11\|564153\|estExt2_Genewise1.C_PHYCAscaffold_140075 |
| 265 | 1 | 17937 | 35 | 24458 | 1 | 1 | 1 | 1 | 0.19 | >jgi\|Phyca11\|17937\|fgenesh1_pg.PHYCAscaffold_32_#_12 |
| 266 | 1 | 538160 | 35 | 6551 | 3 | 3 | 2 | 2 | 2.41 | >jgi\|Phyca11\|538160\|estExt2_Genewise1Plus.C_PHYCAscaffold_10551 |
| 267 | 1 | 536029 | 34 | 80643 | 1 | 1 | 1 | 1 | 0.05 | >jgi\|Phyca11\|536029\|estExt2_fgenesh1_pg.C_PHYCAscaffold_460057 |
| 268 | 1 | 503853 | 34 | 56122 | 1 | 1 | 1 | 1 | 0.08 | >jgi\|Phyca11\|503853\|fgenesh2_kg.PHYCAscaffold_5_#_17_#_Contig1448.1 |
| 269 | 1 | 53999 | 34 | 109401 | 1 | 1 | 1 | 1 | 0.04 | >jgi\|Phyca11\|53999\|gw1.13.151.1 |
| 270 | 1 | 504146 | 34 | 24868 | 2 | 2 | 2 | 2 | 0.4 | >jgi\|Phyca11\|504146\|fgenesh2_kg.PHYCAscaffold_6_#_63_#_Contig1083.1 |
| 271 | 1 | 503902 | 34 | 35315 | 2 | 2 | 2 | 2 | 0.27 | >jgi\|Phyca11\|503902\|fgenesh2_kg.PHYCAscaffold_5_#_66_#_Contig14.1 |
| 272 | 1 | 507207 | 34 | 62676 | 1 | 1 | 1 | 1 | 0.07 | >jgi\|Phyca11\|507207\|fgenesh2_kg.PHYCAscaffold_26_#_10_#_Contig4811.1 |
| 273 | 1 | 111127 | 34 | 66053 | 1 | 1 | 1 | 1 | 0.07 | >jgi\|Phyca11\|111127\|e_gw1.19.476.1 |
| 274 | 1 | 509252 | 33 | 60690 | 1 | 1 | 1 | 1 | 0.07 | >jgi\|Phyca11\|509252\|fgenesh2_kg.PHYCAscaffold_43_#_50_#_gi\|189083975\|gb\|BT031491.1\| |
| 275 | 1 |  | 33 | 26543 | 1 | 1 | 1 | 1 | 0.17 | 223 aa P33049 |
| 276 | 1 | 127049 | 33 | 123862 | 1 | 1 | 1 | 1 | 0.04 | >jgi\|Phyca11\|127049\|e_gw1.66.2.1 |
| 277 | 1 | 546081 | 33 | 59607 | 1 | 1 | 1 | 1 | 0.07 | >jgi\|Phyca11\|546081\|estExt2_Genewise1Plus.C_PHYCAscaffold_200073 |
| 278 | 1 | 504523 | 33 | 36578 | 3 | 3 | 2 | 2 | 0.42 | >jgi\|Phyca11\|504523\|fgenesh2_kg.PHYCAscaffold_8_#_92_#_4101945:54 |
| 279 | 1 | 507292 | 33 | 28550 | 1 | 1 | 1 | 1 | 0.16 | >jgi\|Phyca11\|507292\|fgenesh2_kg.PHYCAscaffold_26_#_95_#_4096992:5 |
| 234 | 2 |  | 32 | 117499 | 3 | 3 | 3 | 3 | 0.12 |  |
| 280 | 1 | 505024 | 32 | 60059 | 2 | 2 | 2 | 2 | 0.15 | >jgi\|Phyca11\|505024\|fgenesh2_kg.PHYCAscaffold_10_#_184_#_4097260:2 |
| 281 | 1 | 577399 | 32 | 82532 | 1 | 1 | 1 | 1 | 0.05 | >jgi\|Phyca11\|577399\|estExt2_Genewise1.C_PHYCAscaffold_1120030 |
| 282 | 1 | 564873 | 32 | 45614 | 1 | 1 | 1 | 1 | 0.1 | >jgi\|Phyca11\|564873\|estExt2_Genewise1.C_PHYCAscaffold_160358 |
| 283 | 1 | 9410 | 32 | 146089 | 1 | 1 | 1 | 1 | 0.03 | >jgi\|Phyca11\|9410\|fgenesh1_pm.PHYCAscaffold_37_#_61 |
| 284 | 1 | 509041 | 32 | 22172 | 1 | 1 | 1 | 1 | 0.21 | >jgi\|Phyca11\|509041\|fgenesh2_kg.PHYCAscaffold_41_#_37_#_Contig2305.1 |
| 285 | 1 | 566838 | 32 | 72303 | 1 | 1 | 1 | 1 | 0.06 | >jgi\|Phyca11\|566838\|estExt2_Genewise1.C_PHYCAscaffold_220535 |
| 286 | 1 | 126045 | 32 | 82427 | 2 | 2 | 2 | 2 | 0.11 | >jgi\|Phyca11\|126045\|e_gw1.61.41.1 |
| 287 | 1 | 503888 | 31 | 22688 | 1 | 1 | 1 | 1 | 0.2 | >jgi\|Phyca11\|503888\|fgenesh2_kg.PHYCAscaffold_5_#_52_#_Contig36.1 |
| 288 | 1 | 511117 | 31 | 18412 | 1 | 1 | 1 | 1 | 0.26 | >jgi\|Phyca11\|511117\|fgenesh2_kg.PHYCAscaffold_76_#_18_#_Contig2045.1 |
| 289 | 1 | 525574 | 31 | 114459 | 1 | 1 | 1 | 1 | 0.04 | >jgi\|Phyca11\|525574\|estExt2_fgenesh1_pm.C_PHYCAscaffold_40114 |
| 290 | 1 | 99958 | 31 | 49559 | 1 | 1 | 1 | 1 | 0.09 | >jgi\|Phyca11\|99958\|e_gw1.4.612.1 |
| 291 | 1 | 119215 | 31 | 20026 | 1 | 1 | 1 | 1 | 0.23 | >jgi\|Phyca11\|119215\|e_gw1.38.51.1 |
| 292 | 1 | 542594 | 31 | 88794 | 1 | 1 | 1 | 1 | 0.05 | >jgi\|Phyca11\|542594\|estExt2_Genewise1Plus.C_PHYCAscaffold_90568 |
| 293 | 1 | 536136 | 31 | 100898 | 1 | 1 | 1 | 1 | 0.04 | >jgi\|Phyca11\|536136\|estExt2_fgenesh1_pg.C_PHYCAscaffold_480011 |
| 294 | 1 | 572181 | 31 | 73595 | 2 | 2 | 2 | 2 | 0.12 | >jgi\|Phyca11\|572181\|estExt2_Genewise1.C_PHYCAscaffold_460371 |
| 295 | 1 | 504259 | 31 | 129675 | 1 | 1 | 1 | 1 | 0.03 | >jgi\|Phyca11\|504259\|fgenesh2_kg.PHYCAscaffold_7_#_11_#_Contig1696.1 |
| 296 | 1 | 543145 | 31 | 143603 | 1 | 1 | 1 | 1 | 0.03 | >jgi\|Phyca11\|543145\|estExt2_Genewise1Plus.C_PHYCAscaffold_110219 |
| 297 | 1 | 556160 | 31 | 71452 | 1 | 1 | 1 | 1 | 0.06 | >jgi\|Phyca11\|556160\|estExt2_Genewise1Plus.C_PHYCAscaffold_840066 |
| 298 | 1 | 509548 | 30 | 90904 | 2 | 2 | 2 | 2 | 0.1 | >jgi\|Phyca11\|509548\|fgenesh2_kg.PHYCAscaffold_47_#_39_#_gi\|189084335\|gb\|BT031851.1\| |
| 299 | 1 | 52778 | 30 | 21158 | 4 | 4 | 1 | 1 | 0.22 | >jgi\|Phyca11\|52778\|gw1.859.2.1 |
| 300 | 1 | 509665 | 30 | 80777 | 1 | 1 | 1 | 1 | 0.05 | >jgi\|Phyca11\|509665\|fgenesh2_kg.PHYCAscaffold_48_#_68_#_Contig411.1 |
| 301 | 1 | 8284 | 30 | 59975 | 1 | 1 | 1 | 1 | 0.07 | >jgi\|Phyca11\|8284\|fgenesh1_pm.PHYCAscaffold_27_#_30 |
| 302 | 1 | 507065 | 30 | 53551 | 2 | 2 | 1 | 1 | 0.08 | >jgi\|Phyca11\|507065\|fgenesh2_kg.PHYCAscaffold_24_#_57_#_Contig26.1 |
| 303 | 1 | 21145 | 30 | 68289 | 2 | 2 | 2 | 2 | 0.13 | >jgi\|Phyca11\|21145\|fgenesh1_pg.PHYCAscaffold_84_#_5 |
| 304 | 1 | 508485 | 30 | 100124 | 1 | 1 | 1 | 1 | 0.04 | >jgi\|Phyca11\|508485\|fgenesh2_kg.PHYCAscaffold_35_#_68_#_Contig1957.1 |
| 305 | 1 | 7598 | 30 | 17660 | 1 | 1 | 1 | 1 | 0.27 | >jgi\|Phyca11\|7598\|fgenesh1_pm.PHYCAscaffold_20_#_113 |
| 306 | 1 | 509550 | 30 | 39887 | 1 | 1 | 1 | 1 | 0.11 | >jgi\|Phyca11\|509550\|fgenesh2_kg.PHYCAscaffold_47_#_41_#_4101945:185 |
| 307 | 1 | 13754 | 30 | 66853 | 2 | 2 | 2 | 2 | 0.14 | >jgi\|Phyca11\|13754\|fgenesh1_pg.PHYCAscaffold_4_#_277 |
| 308 | 1 | 508915 | 29 | 146454 | 1 | 1 | 1 | 1 | 0.03 | >jgi\|Phyca11\|508915\|fgenesh2_kg.PHYCAscaffold_40_#_4_#_Contig2843.1 |
| 309 | 1 | 573714 | 29 | 55596 | 3 | 3 | 3 | 3 | 0.26 | >jgi\|Phyca11\|573714\|estExt2_Genewise1.C_PHYCAscaffold_550085 |
| 310 | 1 | 20316 | 29 | 90212 | 1 | 1 | 1 | 1 | 0.05 | >jgi\|Phyca11\|20316\|fgenesh1_pg.PHYCAscaffold_61_#_44 |
| 311 | 1 | 509735 | 29 | 28698 | 1 | 1 | 1 | 1 | 0.16 | >jgi\|Phyca11\|509735\|fgenesh2_kg.PHYCAscaffold_49_#_27_#_4097066:1 |
| 312 | 1 | 534485 | 29 | 47695 | 2 | 2 | 1 | 1 | 0.09 | >jgi\|Phyca11\|534485\|estExt2_fgenesh1_pg.C_PHYCAscaffold_240045 |
| 313 | 1 |  | 29 | 59053 | 1 | 1 | 1 | 1 | 0.07 |  |
| 314 | 1 | 102840 | 29 | 137993 | 1 | 1 | 1 | 1 | 0.03 | >jgi\|Phyca11\|102840\|e_gw1.7.158.1 |
| 315 | 1 | 540015 | 29 | 12816 | 1 | 1 | 1 | 1 | 0.38 | >jgi\|Phyca11\|540015\|estExt2_Genewise1Plus.C_PHYCAscaffold_40346 |
| 316 | 1 | 502842 | 29 | 47240 | 2 | 2 | 2 | 2 | 0.2 | >jgi\|Phyca11\|502842\|fgenesh2_kg.PHYCAscaffold_1_#_302_#_4096672:1 |
| 317 | 1 | 124543 | 29 | 42728 | 1 | 1 | 1 | 1 | 0.1 | >jgi\|Phyca11\|124543\|e_gw1.54.292.1 |
| 318 | 1 | 104177 | 29 | 38748 | 1 | 1 | 1 | 1 | 0.12 | >jgi\|Phyca11\|104177\|e_gw1.9.229.1 |
| 319 | 1 | 511005 | 28 | 9386 | 1 | 1 | 1 | 1 | 0.54 | >jgi\|Phyca11\|511005\|fgenesh2_kg.PHYCAscaffold_72_#_50_#_gi\|189084915\|gb\|BT032446.1\| |
| 320 | 1 | 558708 | 28 | 12415 | 1 | 1 | 1 | 1 | 0.4 | >jgi\|Phyca11\|558708\|estExt2_Genewise1.C_PHYCAscaffold_20399 |
| 321 | 1 | 12926 | 28 | 54173 | 1 | 1 | 1 | 1 | 0.08 | >jgi\|Phyca11\|12926\|fgenesh1_pg.PHYCAscaffold_2_#_22 |
| 322 | 1 | 108394 | 28 | 31659 | 1 | 1 | 1 | 1 | 0.14 | >jgi\|Phyca11\|108394\|e_gw1.15.711.1 |
| 323 | 1 | 10181 | 28 | 205349 | 1 | 1 | 1 | 1 | 0.02 | >jgi\|Phyca11\|10181\|fgenesh1_pm.PHYCAscaffold_47_#_10 |
| 324 | 1 | 570034 | 28 | 115203 | 2 | 2 | 2 | 2 | 0.08 | >jgi\|Phyca11\|570034\|estExt2_Genewise1.C_PHYCAscaffold_350131 |
| 325 | 1 | 503099 | 28 | 52799 | 1 | 1 | 1 | 1 | 0.08 | >jgi\|Phyca11\|503099\|fgenesh2_kg.PHYCAscaffold_2_#_215_#_4096614:2 |
| 326 | 1 | 116963 | 28 | 95001 | 1 | 1 | 1 | 1 | 0.05 | >jgi\|Phyca11\|116963\|e_gw1.32.12.1 |
| 327 | 1 | 505827 | 28 | 87715 | 1 | 1 | 1 | 1 | 0.05 | >jgi\|Phyca11\|505827\|fgenesh2_kg.PHYCAscaffold_15_#_134_#_Contig345.1 |
| 328 | 1 | 568292 | 28 | 49736 | 1 | 1 | 1 | 1 | 0.09 | >jgi\|Phyca11\|568292\|estExt2_Genewise1.C_PHYCAscaffold_280248 |
| 329 | 1 | 575588 | 28 | 37419 | 2 | 2 | 2 | 2 | 0.25 | >jgi\|Phyca11\|575588\|estExt2_Genewise1.C_PHYCAscaffold_760052 |
| 330 | 1 | 503263 | 28 | 59510 | 1 | 1 | 1 | 1 | 0.07 | >jgi\|Phyca11\|503263\|fgenesh2_kg.PHYCAscaffold_3_#_128_#_Contig1288.1 |
| 331 | 1 | 511439 | 28 | 114572 | 1 | 1 | 1 | 1 | 0.04 | >jgi\|Phyca11\|511439\|fgenesh2_kg.PHYCAscaffold_85_#_7_#_gi\|189084557\|gb\|BT032073.1\| |
| 332 | 1 | 548494 | 27 | 119701 | 2 | 2 | 2 | 2 | 0.07 | >jgi\|Phyca11\|548494\|estExt2_Genewise1Plus.C_PHYCAscaffold_290086 |
| 333 | 1 | 505935 | 27 | 44622 | 2 | 2 | 2 | 2 | 0.21 | >jgi\|Phyca11\|505935\|fgenesh2_kg.PHYCAscaffold_17_#_20_#_4096803:2 |
| 334 | 1 | 566170 | 27 | 30514 | 1 | 1 | 1 | 1 | 0.15 | >jgi\|Phyca11\|566170\|estExt2_Genewise1.C_PHYCAscaffold_200232 |
| 335 | 1 | 536047 | 27 | 80179 | 1 | 1 | 1 | 1 | 0.05 | >jgi\|Phyca11\|536047\|estExt2_fgenesh1_pg.C_PHYCAscaffold_460089 |
| 336 | 1 | 17928 | 27 | 61996 | 2 | 2 | 2 | 2 | 0.15 | >jgi\|Phyca11\|17928\|fgenesh1_pg.PHYCAscaffold_32_#_3 |
| 337 | 1 | 502585 | 27 | 36150 | 1 | 1 | 1 | 1 | 0.12 | >jgi\|Phyca11\|502585\|fgenesh2_kg.PHYCAscaffold_1_#_45_#_Contig972.1 |
| 338 | 1 | 506576 | 27 | 72048 | 2 | 2 | 2 | 2 | 0.13 | >jgi\|Phyca11\|506576\|fgenesh2_kg.PHYCAscaffold_20_#_122_#_Contig362.1 |
| 339 | 1 | 509713 | 27 | 7861 | 1 | 1 | 1 | 1 | 0.67 | >jgi\|Phyca11\|509713\|fgenesh2_kg.PHYCAscaffold_49_#_5_#_gi\|189084943\|gb\|BT032475.1\| |
| 340 | 1 | 507105 | 27 | 72746 | 1 | 1 | 1 | 1 | 0.06 | >jgi\|Phyca11\|507105\|fgenesh2_kg.PHYCAscaffold_24_#_97_#_gi\|189084274\|gb\|BT031790.1\| |
| 341 | 1 | 557319 | 27 | 49059 | 2 | 2 | 1 | 1 | 0.09 | >jgi\|Phyca11\|557319\|estExt2_Genewise1Plus.C_PHYCAscaffold_1070014 |
| 342 | 1 | 503852 | 27 | 17433 | 1 | 1 | 1 | 1 | 0.27 | >jgi\|Phyca11\|503852\|fgenesh2_kg.PHYCAscaffold_5_#_16_#_4101529:5 |
| 343 | 1 | 510172 | 26 | 43700 | 1 | 1 | 1 | 1 | 0.1 | >jgi\|Phyca11\|510172\|fgenesh2_kg.PHYCAscaffold_54_#_66_#_4101945:150 |
| 344 | 1 | 504142 | 26 | 34596 | 2 | 2 | 2 | 2 | 0.28 | >jgi\|Phyca11\|504142\|fgenesh2_kg.PHYCAscaffold_6_#_59_#_4101529:2 |
| 345 | 1 | 507525 | 26 | 56350 | 1 | 1 | 1 | 1 | 0.08 | >jgi\|Phyca11\|507525\|fgenesh2_kg.PHYCAscaffold_28_#_32_#_Contig2244.1 |
| 346 | 1 | 545106 | 26 | 105339 | 1 | 1 | 1 | 1 | 0.04 | >jgi\|Phyca11\|545106\|estExt2_Genewise1Plus.C_PHYCAscaffold_170111 |
| 347 | 1 | 564962 | 26 | 75452 | 3 | 3 | 3 | 3 | 0.18 | >jgi\|Phyca11\|564962\|estExt2_Genewise1.C_PHYCAscaffold_160543 |
| 348 | 1 | 545294 | 26 | 63679 | 1 | 1 | 1 | 1 | 0.07 | >jgi\|Phyca11\|545294\|estExt2_Genewise1Plus.C_PHYCAscaffold_170411 |
| 349 | 1 | 507030 | 26 | 41223 | 1 | 1 | 1 | 1 | 0.11 | >jgi\|Phyca11\|507030\|fgenesh2_kg.PHYCAscaffold_24_#_22_#_Contig274.1 |
| 350 | 1 | 573749 | 26 | 86552 | 1 | 1 | 1 | 1 | 0.05 | >jgi\|Phyca11\|573749\|estExt2_Genewise1.C_PHYCAscaffold_550157 |
| 351 | 1 | 508807 | 26 | 62683 | 1 | 1 | 1 | 1 | 0.07 | >jgi\|Phyca11\|508807\|fgenesh2_kg.PHYCAscaffold_38_#_59_#_Contig2014.1 |
| 352 | 1 | 129924 | 26 | 37470 | 1 | 1 | 1 | 1 | 0.12 | >jgi\|Phyca11\|129924\|e_gw1.89.53.1 |
| 353 | 1 | 507870 | 25 | 16465 | 1 | 1 | 1 | 1 | 0.29 | >jgi\|Phyca11\|507870\|fgenesh2_kg.PHYCAscaffold_30_#_139_#_4101945:254 |
| 354 | 1 | 534395 | 25 | 79780 | 1 | 1 | 1 | 1 | 0.05 | >jgi\|Phyca11\|534395\|estExt2_fgenesh1_pg.C_PHYCAscaffold_230052 |
| 355 | 1 | 527618 | 25 | 121256 | 1 | 1 | 1 | 1 | 0.04 | >jgi\|Phyca11\|527618\|estExt2_fgenesh1_pm.C_PHYCAscaffold_210020 |
| 356 | 1 | 13027 | 25 | 16189 | 1 | 1 | 1 | 1 | 0.29 | >jgi\|Phyca11\|13027\|fgenesh1_pg.PHYCAscaffold_2_#_123 |
| 357 | 1 | 133548 | 25 | 8927 | 1 | 1 | 1 | 1 | 0.58 | >jgi\|Phyca11\|133548\|e_gw1.548.9.1 |
| 358 | 1 | 506049 | 25 | 68815 | 1 | 1 | 1 | 1 | 0.06 | >jgi\|Phyca11\|506049\|fgenesh2_kg.PHYCAscaffold_17_#_134_#_gi\|189084701\|gb\|BT032217.1\| |
| 359 | 1 | 12657 | 25 | 100222 | 1 | 1 | 1 | 1 | 0.04 | >jgi\|Phyca11\|12657\|fgenesh1_pg.PHYCAscaffold_1_#_109 |
| 360 | 1 | 508791 | 25 | 98965 | 1 | 1 | 1 | 1 | 0.04 | >jgi\|Phyca11\|508791\|fgenesh2_kg.PHYCAscaffold_38_#_43_#_Contig2859.1 |
| 361 | 1 | 106472 | 25 | 96852 | 1 | 1 | 1 | 1 | 0.05 | >jgi\|Phyca11\|106472\|e_gw1.12.486.1 |
| 362 | 1 | 508067 | 25 | 24183 | 1 | 1 | 1 | 1 | 0.19 | >jgi\|Phyca11\|508067\|fgenesh2_kg.PHYCAscaffold_32_#_26_#_Contig1225.1 |
| 363 | 1 | 6938 | 25 | 137493 | 1 | 1 | 1 | 1 | 0.03 | >jgi\|Phyca11\|6938\|fgenesh1_pm.PHYCAscaffold_15_#_99 |
| 364 | 1 | 559923 | 25 | 77845 | 1 | 1 | 1 | 1 | 0.06 | >jgi\|Phyca11\|559923\|estExt2_Genewise1.C_PHYCAscaffold_40275 |
| 365 | 1 | 103654 | 24 | 467596 | 1 | 1 | 1 | 1 | 0.01 | >jgi\|Phyca11\|103654\|e_gw1.8.913.1 |
| 366 | 1 | 558527 | 24 | 28603 | 1 | 1 | 1 | 1 | 0.16 | >jgi\|Phyca11\|558527\|estExt2_Genewise1.C_PHYCAscaffold_11463 |
| 367 | 1 | 510978 | 24 | 59574 | 1 | 1 | 1 | 1 | 0.07 | >jgi\|Phyca11\|510978\|fgenesh2_kg.PHYCAscaffold_72_#_23_#_Contig4731.1 |
| 368 | 1 |  | 24 | 24331 | 1 | 1 | 1 | 1 | 0.19 | 214 aa P18626 |
| 369 | 1 | 130473 | 24 | 64751 | 1 | 1 | 1 | 1 | 0.07 | >jgi\|Phyca11\|130473\|e_gw1.94.12.1 |
| 370 | 1 | 566280 | 24 | 72812 | 1 | 1 | 1 | 1 | 0.06 | >jgi\|Phyca11\|566280\|estExt2_Genewise1.C_PHYCAscaffold_200416 |
| 371 | 1 | 20099 | 24 | 26945 | 1 | 1 | 1 | 1 | 0.17 | >jgi\|Phyca11\|20099\|fgenesh1_pg.PHYCAscaffold_57_#_50 |
| 372 | 1 | 537210 | 24 | 38195 | 1 | 1 | 1 | 1 | 0.12 | >jgi\|Phyca11\|537210\|estExt2_fgenesh1_pg.C_PHYCAscaffold_790005 |
| 373 | 1 | 21516 | 23 | 40377 | 1 | 1 | 1 | 1 | 0.11 | >jgi\|Phyca11\|21516\|fgenesh1_pg.PHYCAscaffold_99_#_12 |
| 374 | 1 | 556109 | 23 | 52075 | 1 | 1 | 1 | 1 | 0.09 | >jgi\|Phyca11\|556109\|estExt2_Genewise1Plus.C_PHYCAscaffold_820135 |
| 375 | 1 | 13219 | 23 | 57130 | 1 | 1 | 1 | 1 | 0.08 | >jgi\|Phyca11\|13219\|fgenesh1_pg.PHYCAscaffold_3_#_32 |
| 376 | 1 | 114358 | 23 | 144400 | 1 | 1 | 1 | 1 | 0.03 | >jgi\|Phyca11\|114358\|e_gw1.26.296.1 |
| 377 | 1 | 551419 | 23 | 32513 | 2 | 2 | 2 | 2 | 0.3 | >jgi\|Phyca11\|551419\|estExt2_Genewise1Plus.C_PHYCAscaffold_420070 |
| 378 | 1 | 556069 | 23 | 81718 | 1 | 1 | 1 | 1 | 0.05 | >jgi\|Phyca11\|556069\|estExt2_Genewise1Plus.C_PHYCAscaffold_820056 |
| 379 | 1 | 503383 | 23 | 53444 | 1 | 1 | 1 | 1 | 0.08 | >jgi\|Phyca11\|503383\|fgenesh2_kg.PHYCAscaffold_3_#_248_#_Contig106.1 |
| 380 | 1 | 19832 | 23 | 37148 | 1 | 1 | 1 | 1 | 0.12 | >jgi\|Phyca11\|19832\|fgenesh1_pg.PHYCAscaffold_52_#_84 |
| 381 | 1 | 548475 | 23 | 62936 | 1 | 1 | 1 | 1 | 0.07 | >jgi\|Phyca11\|548475\|estExt2_Genewise1Plus.C_PHYCAscaffold_290043 |
| 382 | 1 | 506158 | 23 | 71254 | 1 | 1 | 1 | 1 | 0.06 | >jgi\|Phyca11\|506158\|fgenesh2_kg.PHYCAscaffold_18_#_31_#_Contig643.1 |
| 383 | 1 | 509232 | 23 | 20790 | 1 | 1 | 1 | 1 | 0.22 | >jgi\|Phyca11\|509232\|fgenesh2_kg.PHYCAscaffold_43_#_30_#_4101945:190 |
| 384 | 1 | 40694 | 23 | 16728 | 1 | 1 | 1 | 1 | 0.28 | >jgi\|Phyca11\|40694\|gw1.1.182.1 |
| 385 | 1 | 570824 | 22 | 52801 | 1 | 1 | 1 | 1 | 0.08 | >jgi\|Phyca11\|570824\|estExt2_Genewise1.C_PHYCAscaffold_390124 |
| 386 | 1 | 573119 | 22 | 52311 | 1 | 1 | 1 | 1 | 0.08 | >jgi\|Phyca11\|573119\|estExt2_Genewise1.C_PHYCAscaffold_510194 |
| 387 | 1 | 528272 | 22 | 65345 | 2 | 2 | 2 | 2 | 0.14 | >jgi\|Phyca11\|528272\|estExt2_fgenesh1_pm.C_PHYCAscaffold_280032 |
| 388 | 1 | 548878 | 22 | 69725 | 1 | 1 | 1 | 1 | 0.06 | >jgi\|Phyca11\|548878\|estExt2_Genewise1Plus.C_PHYCAscaffold_300299 |
| 389 | 1 | 539189 | 22 | 16536 | 2 | 2 | 2 | 2 | 0.66 | >jgi\|Phyca11\|539189\|estExt2_Genewise1Plus.C_PHYCAscaffold_30046 |
| 390 | 1 | 553991 | 22 | 125118 | 1 | 1 | 1 | 1 | 0.03 | >jgi\|Phyca11\|553991\|estExt2_Genewise1Plus.C_PHYCAscaffold_580074 |
| 391 | 1 | 508456 | 22 | 52157 | 1 | 1 | 1 | 1 | 0.08 | >jgi\|Phyca11\|508456\|fgenesh2_kg.PHYCAscaffold_35_#_39_#_gi\|189084740\|gb\|BT032258.1\| |
| 392 | 1 | 535762 | 22 | 61857 | 1 | 1 | 1 | 1 | 0.07 | >jgi\|Phyca11\|535762\|estExt2_fgenesh1_pg.C_PHYCAscaffold_410062 |
| 393 | 1 | 511162 | 22 | 28566 | 1 | 1 | 1 | 1 | 0.16 | >jgi\|Phyca11\|511162\|fgenesh2_kg.PHYCAscaffold_77_#_17_#_4098516:1 |
| 394 | 1 | 565012 | 22 | 59015 | 1 | 1 | 1 | 1 | 0.07 | >jgi\|Phyca11\|565012\|estExt2_Genewise1.C_PHYCAscaffold_170013 |
| 395 | 1 | 506552 | 22 | 30387 | 1 | 1 | 1 | 1 | 0.15 | >jgi\|Phyca11\|506552\|fgenesh2_kg.PHYCAscaffold_20_#_98_#_Contig1804.1 |
| 396 | 1 | 108388 | 21 | 62053 | 1 | 1 | 1 | 1 | 0.07 | >jgi\|Phyca11\|108388\|e_gw1.15.262.1 |
| 397 | 1 | 567679 | 21 | 90155 | 1 | 1 | 1 | 1 | 0.05 | >jgi\|Phyca11\|567679\|estExt2_Genewise1.C_PHYCAscaffold_260120 |
| 398 | 1 | 8949 | 21 | 85750 | 2 | 2 | 1 | 1 | 0.05 | >jgi\|Phyca11\|8949\|fgenesh1_pm.PHYCAscaffold_32_#_84 |
| 399 | 1 | 511694 | 21 | 60273 | 1 | 1 | 1 | 1 | 0.07 | >jgi\|Phyca11\|511694\|fgenesh2_kg.PHYCAscaffold_95_#_23_#_gi\|189084146\|gb\|BT031662.1\| |
| 400 | 1 | 562582 | 21 | 76243 | 2 | 2 | 2 | 2 | 0.12 | >jgi\|Phyca11\|562582\|estExt2_Genewise1.C_PHYCAscaffold_90636 |
| 401 | 1 | 571799 | 21 | 65659 | 1 | 1 | 1 | 1 | 0.07 | >jgi\|Phyca11\|571799\|estExt2_Genewise1.C_PHYCAscaffold_430351 |
| 402 | 1 | 539116 | 21 | 56269 | 1 | 1 | 1 | 1 | 0.08 | >jgi\|Phyca11\|539116\|estExt2_Genewise1Plus.C_PHYCAscaffold_20989 |
| 403 | 1 | 114899 | 21 | 191346 | 1 | 1 | 1 | 1 | 0.02 | >jgi\|Phyca11\|114899\|e_gw1.27.1.1 |
| 404 | 1 | 40390 | 20 | 20746 | 1 | 1 | 1 | 1 | 0.22 | >jgi\|Phyca11\|40390\|gw1.5.116.1 |
| 405 | 1 | 503545 | 20 | 98917 | 1 | 1 | 1 | 1 | 0.04 | >jgi\|Phyca11\|503545\|fgenesh2_kg.PHYCAscaffold_4_#_84_#_Contig5.2 |
| 406 | 1 | 508386 | 20 | 51919 | 1 | 1 | 1 | 1 | 0.09 | >jgi\|Phyca11\|508386\|fgenesh2_kg.PHYCAscaffold_34_#_82_#_4101676:1 |
| 407 | 1 | 504193 | 20 | 150606 | 1 | 1 | 1 | 1 | 0.03 | >jgi\|Phyca11\|504193\|fgenesh2_kg.PHYCAscaffold_6_#_110_#_gi\|189084605\|gb\|BT032121.1\| |
| 408 | 1 | 505405 | 20 | 46756 | 1 | 1 | 1 | 1 | 0.1 | >jgi\|Phyca11\|505405\|fgenesh2_kg.PHYCAscaffold_13_#_55_#_4101849:4 |
| 409 | 1 | 115422 | 20 | 24913 | 1 | 1 | 1 | 1 | 0.18 | >jgi\|Phyca11\|115422\|e_gw1.28.453.1 |
| 410 | 1 | 79205 | 19 | 25929 | 1 | 1 | 1 | 1 | 0.18 | >jgi\|Phyca11\|79205\|gw1.28.598.1 |
| 411 | 1 | 552811 | 19 | 61340 | 1 | 1 | 1 | 1 | 0.07 | >jgi\|Phyca11\|552811\|estExt2_Genewise1Plus.C_PHYCAscaffold_490242 |
| 412 | 1 | 534052 | 19 | 110330 | 2 | 2 | 1 | 1 | 0.04 | >jgi\|Phyca11\|534052\|estExt2_fgenesh1_pg.C_PHYCAscaffold_200013 |
| 413 | 1 | 544178 | 19 | 71000 | 1 | 1 | 1 | 1 | 0.06 | >jgi\|Phyca11\|544178\|estExt2_Genewise1Plus.C_PHYCAscaffold_140074 |
| 414 | 1 | 103847 | 19 | 21913 | 1 | 1 | 1 | 1 | 0.21 | >jgi\|Phyca11\|103847\|e_gw1.8.798.1 |
| 415 | 1 | 506922 | 19 | 14320 | 1 | 1 | 1 | 1 | 0.34 | >jgi\|Phyca11\|506922\|fgenesh2_kg.PHYCAscaffold_23_#_53_#_4101134:1 |
| 416 | 1 | 529934 | 19 | 121660 | 2 | 2 | 2 | 2 | 0.07 | >jgi\|Phyca11\|529934\|estExt2_fgenesh1_pm.C_PHYCAscaffold_510045 |
| 417 | 1 | 529559 | 19 | 122629 | 1 | 1 | 1 | 1 | 0.04 | >jgi\|Phyca11\|529559\|estExt2_fgenesh1_pm.C_PHYCAscaffold_460012 |
| 418 | 1 | 528038 | 19 | 193379 | 2 | 2 | 2 | 2 | 0.05 | >jgi\|Phyca11\|528038\|estExt2_fgenesh1_pm.C_PHYCAscaffold_260026 |
| 419 | 1 | 503926 | 19 | 17112 | 1 | 1 | 1 | 1 | 0.28 | >jgi\|Phyca11\|503926\|fgenesh2_kg.PHYCAscaffold_5_#_90_#_4097723:1 |
| 420 | 1 | 553558 | 19 | 156357 | 1 | 1 | 1 | 1 | 0.03 | >jgi\|Phyca11\|553558\|estExt2_Genewise1Plus.C_PHYCAscaffold_540017 |
| 421 | 1 | 551325 | 18 | 50894 | 1 | 1 | 1 | 1 | 0.09 | >jgi\|Phyca11\|551325\|estExt2_Genewise1Plus.C_PHYCAscaffold_410240 |
| 422 | 1 | 538731 | 18 | 45074 | 1 | 1 | 1 | 1 | 0.1 | >jgi\|Phyca11\|538731\|estExt2_Genewise1Plus.C_PHYCAscaffold_20342 |
| 423 | 1 | 528255 | 18 | 94441 | 1 | 1 | 1 | 1 | 0.05 | >jgi\|Phyca11\|528255\|estExt2_fgenesh1_pm.C_PHYCAscaffold_280010 |
| 424 | 1 | 507961 | 18 | 82552 | 1 | 1 | 1 | 1 | 0.05 | >jgi\|Phyca11\|507961\|fgenesh2_kg.PHYCAscaffold_31_#_42_#_gi\|189092330\|gb\|BT032284.1\| |
| 425 | 1 | 510434 | 18 | 27753 | 2 | 2 | 1 | 1 | 0.16 | >jgi\|Phyca11\|510434\|fgenesh2_kg.PHYCAscaffold_60_#_22_#_gi\|189084804\|gb\|BT032334.1\| |
| 426 | 1 | 507252 | 18 | 46760 | 1 | 1 | 1 | 1 | 0.1 | >jgi\|Phyca11\|507252\|fgenesh2_kg.PHYCAscaffold_26_#_55_#_4098046:1 |
| 427 | 1 | 507278 | 18 | 52801 | 1 | 1 | 1 | 1 | 0.08 | >jgi\|Phyca11\|507278\|fgenesh2_kg.PHYCAscaffold_26_#_81_#_gi\|189084092\|gb\|BT031608.1\| |
| 428 | 1 | 133727 | 18 | 30614 | 1 | 1 | 1 | 1 | 0.15 | >jgi\|Phyca11\|133727\|e_gw1.683.1.1 |
| 429 | 1 | 502589 | 18 | 46213 | 1 | 1 | 1 | 1 | 0.1 | >jgi\|Phyca11\|502589\|fgenesh2_kg.PHYCAscaffold_1_#_49_#_Contig2680.1 |
| 430 | 1 | 81118 | 18 | 12821 | 1 | 1 | 1 | 1 | 0.38 | >jgi\|Phyca11\|81118\|gw1.49.421.1 |
| 431 | 1 | 108292 | 18 | 103095 | 1 | 1 | 1 | 1 | 0.04 | >jgi\|Phyca11\|108292\|e_gw1.15.153.1 |
| 432 | 1 | 110509 | 18 | 54395 | 1 | 1 | 1 | 1 | 0.08 | >jgi\|Phyca11\|110509\|e_gw1.18.170.1 |
| 433 | 1 | 528840 | 18 | 52324 | 1 | 1 | 1 | 1 | 0.08 | >jgi\|Phyca11\|528840\|estExt2_fgenesh1_pm.C_PHYCAscaffold_340073 |
| 434 | 1 | 113930 | 18 | 39501 | 1 | 1 | 1 | 1 | 0.11 | >jgi\|Phyca11\|113930\|e_gw1.25.661.1 |
| 435 | 1 | 114721 | 17 | 45209 | 1 | 1 | 1 | 1 | 0.1 | >jgi\|Phyca11\|114721\|e_gw1.27.288.1 |
| 436 | 1 | 511910 | 17 | 25014 | 2 | 2 | 1 | 1 | 0.18 | >jgi\|Phyca11\|511910\|fgenesh2_kg.PHYCAscaffold_105_#_8_#_4098198:1 |
| 437 | 1 | 74711 | 17 | 51341 | 1 | 1 | 1 | 1 | 0.09 | >jgi\|Phyca11\|74711\|gw1.8.558.1 |
| 438 | 1 | 14100 | 17 | 15164 | 1 | 1 | 1 | 1 | 0.32 | >jgi\|Phyca11\|14100\|fgenesh1_pg.PHYCAscaffold_6_#_22 |
| 439 | 1 | 532470 | 17 | 16315 | 1 | 1 | 1 | 1 | 0.29 | >jgi\|Phyca11\|532470\|estExt2_fgenesh1_pg.C_PHYCAscaffold_50218 |
| 440 | 1 | 11017 | 17 | 29284 | 1 | 1 | 1 | 1 | 0.16 | >jgi\|Phyca11\|11017\|fgenesh1_pm.PHYCAscaffold_60_#_25 |
| 441 | 1 |  | 17 | 50832 | 1 | 1 | 1 | 1 | 0.09 | (Bos taurus) similar to alpha-tubulin I isoform 1 |
| 442 | 1 | 505595 | 17 | 37961 | 1 | 1 | 1 | 1 | 0.12 | >jgi\|Phyca11\|505595\|fgenesh2_kg.PHYCAscaffold_14_#_76_#_Contig4404.1 |
| 443 | 1 | 551747 | 17 | 61827 | 1 | 1 | 1 | 1 | 0.07 | >jgi\|Phyca11\|551747\|estExt2_Genewise1Plus.C_PHYCAscaffold_430233 |
| 444 | 1 | 133399 | 16 | 33163 | 1 | 1 | 1 | 1 | 0.14 | >jgi\|Phyca11\|133399\|e_gw1.442.4.1 |
| 445 | 1 | 532804 | 16 | 73700 | 1 | 1 | 1 | 1 | 0.06 | >jgi\|Phyca11\|532804\|estExt2_fgenesh1_pg.C_PHYCAscaffold_80096 |
| 446 | 1 | 122330 | 16 | 63337 | 1 | 1 | 1 | 1 | 0.07 | >jgi\|Phyca11\|122330\|e_gw1.47.420.1 |
| 447 | 1 | 127727 | 16 | 113405 | 1 | 1 | 1 | 1 | 0.04 | >jgi\|Phyca11\|127727\|e_gw1.71.37.1 |
| 448 | 1 | 565056 | 16 | 54364 | 1 | 1 | 1 | 1 | 0.08 | >jgi\|Phyca11\|565056\|estExt2_Genewise1.C_PHYCAscaffold_170094 |
| 449 | 1 | 573718 | 16 | 50006 | 1 | 1 | 1 | 1 | 0.09 | >jgi\|Phyca11\|573718\|estExt2_Genewise1.C_PHYCAscaffold_550091 |
| 450 | 1 | 505864 | 16 | 59687 | 1 | 1 | 1 | 1 | 0.07 | >jgi\|Phyca11\|505864\|fgenesh2_kg.PHYCAscaffold_16_#_23_#_4101827:1 |
| 451 | 1 | 535447 | 16 | 86282 | 1 | 1 | 1 | 1 | 0.05 | >jgi\|Phyca11\|535447\|estExt2_fgenesh1_pg.C_PHYCAscaffold_360034 |
| 452 | 1 | 10726 | 16 | 18077 | 1 | 1 | 1 | 1 | 0.26 | >jgi\|Phyca11\|10726\|fgenesh1_pm.PHYCAscaffold_54_#_9 |
| 453 | 1 | 117612 | 16 | 62422 | 1 | 1 | 1 | 1 | 0.07 | >jgi\|Phyca11\|117612\|e_gw1.33.228.1 |
| 454 | 1 | 508846 | 16 | 51821 | 1 | 1 | 1 | 1 | 0.09 | >jgi\|Phyca11\|508846\|fgenesh2_kg.PHYCAscaffold_39_#_13_#_gi\|189084611\|gb\|BT032127.1\| |
| 455 | 1 | 509028 | 15 | 55156 | 1 | 1 | 1 | 1 | 0.08 | >jgi\|Phyca11\|509028\|fgenesh2_kg.PHYCAscaffold_41_#_24_#_Contig2661.1 |
| 456 | 1 | 534319 | 15 | 60205 | 1 | 1 | 1 | 1 | 0.07 | >jgi\|Phyca11\|534319\|estExt2_fgenesh1_pg.C_PHYCAscaffold_220087 |
| 457 | 1 | 118683 | 15 | 17340 | 1 | 1 | 1 | 1 | 0.27 | >jgi\|Phyca11\|118683\|e_gw1.36.498.1 |
| 458 | 1 | 506642 | 15 | 9094 | 1 | 1 | 1 | 1 | 0.57 | >jgi\|Phyca11\|506642\|fgenesh2_kg.PHYCAscaffold_21_#_14_#_Contig5387.1 |
| 459 | 1 | 107310 | 15 | 84217 | 1 | 1 | 1 | 1 | 0.05 | >jgi\|Phyca11\|107310\|e_gw1.13.173.1 |
| 460 | 1 |  | 15 | 36431 | 1 | 1 | 1 | 1 | 0.12 | hypothetical protein RBSDVs7gp2 [Rice black streaked dwarf virus] |
| 461 | 1 | 97325 | 15 | 9553 | 1 | 1 | 1 | 1 | 0.54 | >jgi\|Phyca11\|97325\|e_gw1.1.235.1 |
| 462 | 1 | 511388 | 15 | 76448 | 1 | 1 | 1 | 1 | 0.06 | >jgi\|Phyca11\|511388\|fgenesh2_kg.PHYCAscaffold_84_#_4_#_Contig937.1 |
| 463 | 1 | 546033 | 15 | 52534 | 1 | 1 | 1 | 1 | 0.08 | >jgi\|Phyca11\|546033\|estExt2_Genewise1Plus.C_PHYCAscaffold_190540 |
| 464 | 1 | 550581 | 15 | 72617 | 1 | 1 | 1 | 1 | 0.06 | >jgi\|Phyca11\|550581\|estExt2_Genewise1Plus.C_PHYCAscaffold_370357 |
| 465 | 1 | 113258 | 15 | 34927 | 1 | 1 | 1 | 1 | 0.13 | >jgi\|Phyca11\|113258\|e_gw1.23.56.1 |
| 466 | 1 | 7028 | 15 | 13525 | 1 | 1 | 1 | 1 | 0.36 | >jgi\|Phyca11\|7028\|fgenesh1_pm.PHYCAscaffold_16_#_49 |
| 467 | 1 | 508503 | 15 | 44812 | 1 | 1 | 1 | 1 | 0.1 | >jgi\|Phyca11\|508503\|fgenesh2_kg.PHYCAscaffold_35_#_86_#_Contig1273.1 |
| 468 | 1 | 96024 | 14 | 36676 | 1 | 1 | 1 | 1 | 0.12 | >jgi\|Phyca11\|96024\|e_gw1.1.1208.1 |
| 469 | 1 | 125348 | 14 | 139047 | 1 | 1 | 1 | 1 | 0.03 | >jgi\|Phyca11\|125348\|e_gw1.58.66.1 |
| 470 | 1 | 113987 | 14 | 49064 | 1 | 1 | 1 | 1 | 0.09 | >jgi\|Phyca11\|113987\|e_gw1.25.320.1 |
| 471 | 1 | 558600 | 14 | 43585 | 1 | 1 | 1 | 1 | 0.1 | >jgi\|Phyca11\|558600\|estExt2_Genewise1.C_PHYCAscaffold_20201 |
| 472 | 1 | 571830 | 14 | 66836 | 1 | 1 | 1 | 1 | 0.07 | >jgi\|Phyca11\|571830\|estExt2_Genewise1.C_PHYCAscaffold_430388 |
| 473 | 1 | 16825 | 14 | 366643 | 1 | 1 | 1 | 1 | 0.01 | >jgi\|Phyca11\|16825\|fgenesh1_pg.PHYCAscaffold_22_#_104 |
| 474 | 1 | 104701 | 14 | 37214 | 1 | 1 | 1 | 1 | 0.12 | >jgi\|Phyca11\|104701\|e_gw1.9.628.1 |
| 475 | 1 | 527873 | 14 | 19228 | 1 | 1 | 1 | 1 | 0.24 | >jgi\|Phyca11\|527873\|estExt2_fgenesh1_pm.C_PHYCAscaffold_230104 |
| 476 | 1 | 533060 | 14 | 90951 | 1 | 1 | 1 | 1 | 0.05 | >jgi\|Phyca11\|533060\|estExt2_fgenesh1_pg.C_PHYCAscaffold_100047 |
| 477 | 1 | 108538 | 14 | 24183 | 1 | 1 | 1 | 1 | 0.19 | >jgi\|Phyca11\|108538\|e_gw1.15.770.1 |
| 478 | 1 | 120599 | 14 | 35948 | 1 | 1 | 1 | 1 | 0.13 | >jgi\|Phyca11\|120599\|e_gw1.42.174.1 |
| 479 | 1 | 560279 | 14 | 47305 | 1 | 1 | 1 | 1 | 0.09 | >jgi\|Phyca11\|560279\|estExt2_Genewise1.C_PHYCAscaffold_40823 |
