## Supplementary Materials 1-F1t for "Vacuolar H^+^-ATPase subunit a was identified as the target protein of the oomycete inhibitor fluopicolide"

| Family | Member | Protein ID | Score | Mass | Num. of matches | Num. of significant matches | Num. of sequences | Num. of significant sequences | emPAI | Gene ID |
| --- | --- | --- | --- | --- | --- | --- | --- | --- | --- | --- |
| 1 | 1 | 525392 | 1516 | 94498 | 57 | 57 | 24 | 24 | 2.24 | >jgi\|Phyca11\|525392\|estExt2_fgenesh1_pm.C_PHYCAscaffold_30158 |
| 2 | 2 | 558578 | 1131 | 138453 | 51 | 51 | 28 | 28 | 1.53 | >jgi\|Phyca11\|558578\|estExt2_Genewise1.C_PHYCAscaffold_20133 |
| 3 | 3 | 108059 | 871 | 102760 | 33 | 33 | 19 | 19 | 1.2 | >jgi\|Phyca11\|108059\|e_gw1.14.147.1 |
| 5 | 4 | 545685 | 589 | 107998 | 30 | 30 | 19 | 19 | 1.12 | >jgi\|Phyca11\|545685\|estExt2_Genewise1Plus.C_PHYCAscaffold_180528 |
| 6 | 5 | 98712 | 544 | 60979 | 27 | 27 | 16 | 16 | 2.52 | >jgi\|Phyca11\|98712\|e_gw1.3.703.1 |
| 7 | 6 | 509820 | 529 | 86980 | 23 | 23 | 15 | 15 | 1.09 | >jgi\|Phyca11\|509820\|fgenesh2_kg.PHYCAscaffold_50_#_21_#_Contig158.1 |
| 9 | 7 | 511907 | 516 | 96155 | 25 | 25 | 16 | 16 | 1.33 | >jgi\|Phyca11\|511907\|fgenesh2_kg.PHYCAscaffold_105_#_5_#_4098187:2 |
| 10 | 8 | 562095 | 496 | 195387 | 25 | 25 | 21 | 21 | 0.58 | >jgi\|Phyca11\|562095\|estExt2_Genewise1.C_PHYCAscaffold_80628 |
| 11 | 9 | 503562 | 443 | 116908 | 21 | 21 | 14 | 14 | 0.67 | >jgi\|Phyca11\|503562\|fgenesh2_kg.PHYCAscaffold_4_#_101_#_4100091:2 |
| 13 | 10 | 537721 | 440 | 706535 | 29 | 29 | 23 | 23 | 0.15 | >jgi\|Phyca11\|537721\|estExt2_fgenesh1_pg.C_PHYCAscaffold_1090004 |
| 14 | 11 | 511385 | 415 | 115018 | 19 | 19 | 13 | 13 | 0.62 | >jgi\|Phyca11\|511385\|fgenesh2_kg.PHYCAscaffold_84_#_1_#_Contig68.1 |
| 15 | 12 | 509101 | 414 | 125590 | 16 | 16 | 10 | 10 | 0.41 | >jgi\|Phyca11\|509101\|fgenesh2_kg.PHYCAscaffold_42_#_8_#_Contig2284.1 |
| 5 | 13 | 545682 | 389 | 107341 | 20 | 20 | 16 | 16 | 0.89 | >jgi\|Phyca11\|545682\|estExt2_Genewise1Plus.C_PHYCAscaffold_180525 |
| 16 | 14 | 534535 | 372 | 130373 | 22 | 22 | 15 | 15 | 0.64 | >jgi\|Phyca11\|534535\|estExt2_fgenesh1_pg.C_PHYCAscaffold_250008 |
| 17 | 15 | 510618 | 356 | 108675 | 17 | 17 | 12 | 12 | 0.73 | >jgi\|Phyca11\|510618\|fgenesh2_kg.PHYCAscaffold_64_#_11_#_Contig39.1 |
| 17 | 16 | 20426 | 344 | 125040 | 17 | 17 | 12 | 12 | 0.61 | >jgi\|Phyca11\|20426\|fgenesh1_pg.PHYCAscaffold_64_#_12 |
| 18 | 17 | 107407 | 334 | 25619 | 15 | 15 | 10 | 10 | 5.11 | >jgi\|Phyca11\|107407\|e_gw1.13.765.1 |
| 19 | 18 | 17480 | 324 | 250999 | 17 | 17 | 14 | 14 | 0.27 | >jgi\|Phyca11\|17480\|fgenesh1_pg.PHYCAscaffold_28_#_15 |
| 20 | 19 | 502787 | 294 | 108569 | 16 | 16 | 13 | 13 | 0.67 | >jgi\|Phyca11\|502787\|fgenesh2_kg.PHYCAscaffold_1_#_247_#_gi\|189084186\|gb\|BT031702.1\| |
| 21 | 20 | 505109 | 279 | 108406 | 9 | 9 | 7 | 7 | 0.32 | >jgi\|Phyca11\|505109\|fgenesh2_kg.PHYCAscaffold_11_#_55_#_4100846:2 |
| 18 | 2 | 70742 | 278 | 19036 | 9 | 9 | 7 | 7 | 3.69 | >jgi\|Phyca11\|70742\|gw1.13.784.1 |
| 22 | 1 | 548602 | 264 | 69141 | 12 | 12 | 8 | 8 | 0.64 | >jgi\|Phyca11\|548602\|estExt2_Genewise1Plus.C_PHYCAscaffold_290268 |
| 23 | 1 | 505974 | 254 | 81161 | 12 | 12 | 7 | 7 | 0.44 | >jgi\|Phyca11\|505974\|fgenesh2_kg.PHYCAscaffold_17_#_59_#_4098047:1 |
| 24 | 1 | 116367 | 242 | 96518 | 18 | 18 | 13 | 13 | 0.86 | >jgi\|Phyca11\|116367\|e_gw1.30.3.1 |
| 25 | 1 | 506502 | 230 | 105481 | 12 | 12 | 11 | 11 | 0.56 | >jgi\|Phyca11\|506502\|fgenesh2_kg.PHYCAscaffold_20_#_48_#_gi\|189084416\|gb\|BT031932.1\| |
| 27 | 1 | 572341 | 226 | 71559 | 11 | 11 | 9 | 9 | 0.71 | >jgi\|Phyca11\|572341\|estExt2_Genewise1.C_PHYCAscaffold_470331 |
| 28 | 1 | 534200 | 225 | 152568 | 12 | 12 | 8 | 8 | 0.25 | >jgi\|Phyca11\|534200\|estExt2_fgenesh1_pg.C_PHYCAscaffold_210041 |
| 29 | 1 | 34163 | 224 | 90948 | 13 | 13 | 10 | 10 | 0.6 | >jgi\|Phyca11\|34163\|gw1.2.21.1 |
| 30 | 1 | 531039 | 221 | 101658 | 13 | 13 | 11 | 11 | 0.66 | >jgi\|Phyca11\|531039\|estExt2_fgenesh1_pm.C_PHYCAscaffold_910003 |
| 31 | 1 | 15447 | 220 | 248333 | 17 | 17 | 14 | 14 | 0.27 | >jgi\|Phyca11\|15447\|fgenesh1_pg.PHYCAscaffold_13_#_135 |
| 32 | 1 | 536136 | 205 | 100898 | 10 | 10 | 7 | 7 | 0.34 | >jgi\|Phyca11\|536136\|estExt2_fgenesh1_pg.C_PHYCAscaffold_480011 |
| 33 | 1 | 509055 | 201 | 96457 | 11 | 11 | 11 | 11 | 0.63 | >jgi\|Phyca11\|509055\|fgenesh2_kg.PHYCAscaffold_41_#_51_#_4101815:2 |
| 34 | 1 | 6748 | 200 | 140772 | 9 | 9 | 7 | 7 | 0.24 | >jgi\|Phyca11\|6748\|fgenesh1_pm.PHYCAscaffold_14_#_52 |
| 35 | 1 | 525609 | 200 | 99796 | 6 | 6 | 6 | 6 | 0.29 | >jgi\|Phyca11\|525609\|estExt2_fgenesh1_pm.C_PHYCAscaffold_40150 |
| 36 | 1 | 506056 | 196 | 94357 | 8 | 8 | 7 | 7 | 0.37 | >jgi\|Phyca11\|506056\|fgenesh2_kg.PHYCAscaffold_17_#_141_#_gi\|189084230\|gb\|BT031746.1\| |
| 28 | 2 | 512023 | 190 | 119916 | 15 | 15 | 10 | 10 | 0.48 | >jgi\|Phyca11\|512023\|fgenesh2_kg.PHYCAscaffold_117_#_4_#_Contig156.1 |
| 37 | 1 | 510361 | 190 | 140057 | 10 | 10 | 8 | 8 | 0.28 | >jgi\|Phyca11\|510361\|fgenesh2_kg.PHYCAscaffold_58_#_74_#_gi\|189084453\|gb\|BT031969.1\| |
| 38 | 1 | 542884 | 190 | 97887 | 10 | 10 | 9 | 9 | 0.48 | >jgi\|Phyca11\|542884\|estExt2_Genewise1Plus.C_PHYCAscaffold_100379 |
| 39 | 1 | 558277 | 188 | 130749 | 8 | 8 | 7 | 7 | 0.3 | >jgi\|Phyca11\|558277\|estExt2_Genewise1.C_PHYCAscaffold_10916 |
| 40 | 1 | 99561 | 184 | 110284 | 9 | 9 | 7 | 7 | 0.31 | >jgi\|Phyca11\|99561\|e_gw1.4.365.1 |
| 41 | 1 | 21374 | 169 | 16113 | 5 | 5 | 3 | 3 | 1.18 | >jgi\|Phyca11\|21374\|fgenesh1_pg.PHYCAscaffold_93_#_8 |
| 42 | 1 | 107310 | 169 | 84217 | 9 | 9 | 7 | 7 | 0.43 | >jgi\|Phyca11\|107310\|e_gw1.13.173.1 |
| 43 | 1 | 510859 | 167 | 48785 | 11 | 11 | 8 | 8 | 1.19 | >jgi\|Phyca11\|510859\|fgenesh2_kg.PHYCAscaffold_70_#_5_#_Contig3.1 |
| 28 | 3 | 557455 | 166 | 149278 | 13 | 13 | 8 | 8 | 0.29 | >jgi\|Phyca11\|557455\|estExt2_Genewise1Plus.C_PHYCAscaffold_1170012 |
| 44 | 1 | 503425 | 166 | 29978 | 7 | 7 | 5 | 5 | 1.02 | >jgi\|Phyca11\|503425\|fgenesh2_kg.PHYCAscaffold_3_#_290_#_4101945:196 |
| 45 | 1 | 98041 | 162 | 98789 | 6 | 6 | 6 | 6 | 0.3 | >jgi\|Phyca11\|98041\|e_gw1.2.524.1 |
| 46 | 1 | 575784 | 161 | 65395 | 9 | 9 | 7 | 7 | 0.58 | >jgi\|Phyca11\|575784\|estExt2_Genewise1.C_PHYCAscaffold_790031 |
| 47 | 1 | 538943 | 159 | 167258 | 9 | 9 | 9 | 9 | 0.26 | >jgi\|Phyca11\|538943\|estExt2_Genewise1Plus.C_PHYCAscaffold_20665 |
| 48 | 1 | 504894 | 157 | 121029 | 11 | 11 | 9 | 9 | 0.37 | >jgi\|Phyca11\|504894\|fgenesh2_kg.PHYCAscaffold_10_#_54_#_Contig182.1 |
| 49 | 1 | 553954 | 154 | 125096 | 8 | 8 | 6 | 6 | 0.23 | >jgi\|Phyca11\|553954\|estExt2_Genewise1Plus.C_PHYCAscaffold_580009 |
| 50 | 1 | 503602 | 153 | 83544 | 6 | 6 | 6 | 6 | 0.36 | >jgi\|Phyca11\|503602\|fgenesh2_kg.PHYCAscaffold_4_#_141_#_gi\|189084094\|gb\|BT031610.1\| |
| 51 | 1 | 507207 | 151 | 62676 | 8 | 8 | 5 | 5 | 0.4 | >jgi\|Phyca11\|507207\|fgenesh2_kg.PHYCAscaffold_26_#_10_#_Contig4811.1 |
| 52 | 1 | 504262 | 149 | 89471 | 8 | 8 | 7 | 7 | 0.4 | >jgi\|Phyca11\|504262\|fgenesh2_kg.PHYCAscaffold_7_#_14_#_4101715:1 |
| 53 | 1 | 506722 | 148 | 127848 | 6 | 6 | 6 | 6 | 0.22 | >jgi\|Phyca11\|506722\|fgenesh2_kg.PHYCAscaffold_21_#_94_#_4096595:1 |
| 54 | 1 | 507278 | 147 | 52801 | 7 | 7 | 7 | 7 | 0.76 | >jgi\|Phyca11\|507278\|fgenesh2_kg.PHYCAscaffold_26_#_81_#_gi\|189084092\|gb\|BT031608.1\| |
| 55 | 1 | 503383 | 147 | 53444 | 6 | 6 | 3 | 3 | 0.38 | >jgi\|Phyca11\|503383\|fgenesh2_kg.PHYCAscaffold_3_#_248_#_Contig106.1 |
| 56 | 1 | 563829 | 144 | 32177 | 3 | 3 | 2 | 2 | 0.3 | >jgi\|Phyca11\|563829\|estExt2_Genewise1.C_PHYCAscaffold_130199 |
| 28 | 4 | 20570 | 142 | 66384 | 11 | 11 | 7 | 7 | 0.67 | >jgi\|Phyca11\|20570\|fgenesh1_pg.PHYCAscaffold_66_#_46 |
| 58 | 1 | 548494 | 141 | 119701 | 5 | 5 | 5 | 5 | 0.2 | >jgi\|Phyca11\|548494\|estExt2_Genewise1Plus.C_PHYCAscaffold_290086 |
| 59 | 1 | 511439 | 138 | 114572 | 8 | 8 | 8 | 8 | 0.35 | >jgi\|Phyca11\|511439\|fgenesh2_kg.PHYCAscaffold_85_#_7_#_gi\|189084557\|gb\|BT032073.1\| |
| 60 | 1 | 511305 | 137 | 88501 | 6 | 6 | 6 | 6 | 0.34 | >jgi\|Phyca11\|511305\|fgenesh2_kg.PHYCAscaffold_80_#_49_#_4098993:3 |
| 19 | 2 | 533489 | 134 | 176235 | 6 | 6 | 5 | 5 | 0.13 | >jgi\|Phyca11\|533489\|estExt2_fgenesh1_pg.C_PHYCAscaffold_140069 |
| 28 | 5 | 127018 | 133 | 108376 | 10 | 10 | 8 | 8 | 0.43 | >jgi\|Phyca11\|127018\|e_gw1.66.38.1 |
| 61 | 1 | 127049 | 132 | 123862 | 7 | 7 | 6 | 6 | 0.27 | >jgi\|Phyca11\|127049\|e_gw1.66.2.1 |
| 62 | 1 | 504612 | 132 | 117382 | 12 | 12 | 10 | 10 | 0.49 | >jgi\|Phyca11\|504612\|fgenesh2_kg.PHYCAscaffold_9_#_3_#_Contig947.1 |
| 63 | 1 | 505188 | 132 | 27120 | 3 | 3 | 3 | 3 | 0.6 | >jgi\|Phyca11\|505188\|fgenesh2_kg.PHYCAscaffold_11_#_134_#_Contig42.1 |
| 64 | 1 | 531982 | 130 | 130522 | 8 | 8 | 8 | 8 | 0.3 | >jgi\|Phyca11\|531982\|estExt2_fgenesh1_pg.C_PHYCAscaffold_30148 |
| 65 | 1 | 502852 | 129 | 23768 | 6 | 6 | 5 | 5 | 1.43 | >jgi\|Phyca11\|502852\|fgenesh2_kg.PHYCAscaffold_1_#_312_#_Contig157.1 |
| 67 | 1 | 503727 | 128 | 64500 | 5 | 5 | 5 | 5 | 0.39 | >jgi\|Phyca11\|503727\|fgenesh2_kg.PHYCAscaffold_4_#_266_#_Contig7.1 |
| 68 | 1 | 131909 | 128 | 58808 | 2 | 2 | 2 | 2 | 0.16 | >jgi\|Phyca11\|131909\|e_gw1.121.14.1 |
| 69 | 1 | 504155 | 127 | 20467 | 6 | 6 | 5 | 5 | 1.8 | >jgi\|Phyca11\|504155\|fgenesh2_kg.PHYCAscaffold_6_#_72_#_4101945:207 |
| 70 | 1 | 125348 | 126 | 139047 | 7 | 7 | 6 | 6 | 0.2 | >jgi\|Phyca11\|125348\|e_gw1.58.66.1 |
| 71 | 1 | 555787 | 123 | 90055 | 5 | 5 | 5 | 5 | 0.27 | >jgi\|Phyca11\|555787\|estExt2_Genewise1Plus.C_PHYCAscaffold_780059 |
| 72 | 1 | 564009 | 123 | 112552 | 7 | 7 | 7 | 7 | 0.3 | >jgi\|Phyca11\|564009\|estExt2_Genewise1.C_PHYCAscaffold_130507 |
| 73 | 1 | 124610 | 122 | 89043 | 9 | 9 | 8 | 8 | 0.47 | >jgi\|Phyca11\|124610\|e_gw1.54.84.1 |
| 74 | 1 | 503258 | 122 | 42657 | 7 | 7 | 5 | 5 | 0.65 | >jgi\|Phyca11\|503258\|fgenesh2_kg.PHYCAscaffold_3_#_123_#_Contig19.2 |
| 28 | 6 | 504139 | 121 | 154076 | 7 | 7 | 5 | 5 | 0.15 | >jgi\|Phyca11\|504139\|fgenesh2_kg.PHYCAscaffold_6_#_56_#_Contig215.1 |
| 75 | 1 | 533987 | 121 | 136648 | 8 | 8 | 6 | 6 | 0.21 | >jgi\|Phyca11\|533987\|estExt2_fgenesh1_pg.C_PHYCAscaffold_190079 |
| 76 | 1 | 534052 | 120 | 110330 | 6 | 6 | 5 | 5 | 0.21 | >jgi\|Phyca11\|534052\|estExt2_fgenesh1_pg.C_PHYCAscaffold_200013 |
| 77 | 1 | 568901 | 119 | 159351 | 5 | 5 | 5 | 5 | 0.14 | >jgi\|Phyca11\|568901\|estExt2_Genewise1.C_PHYCAscaffold_300383 |
| 78 | 1 | 528052 | 119 | 132419 | 6 | 6 | 5 | 5 | 0.18 | >jgi\|Phyca11\|528052\|estExt2_fgenesh1_pm.C_PHYCAscaffold_260044 |
| 79 | 1 | 552387 | 119 | 194533 | 2 | 2 | 2 | 2 | 0.04 | >jgi\|Phyca11\|552387\|estExt2_Genewise1Plus.C_PHYCAscaffold_470321 |
| 19 | 3 | 571405 | 116 | 163648 | 4 | 4 | 4 | 4 | 0.11 | >jgi\|Phyca11\|571405\|estExt2_Genewise1.C_PHYCAscaffold_420119 |
| 80 | 1 | 569541 | 116 | 68789 | 7 | 7 | 6 | 6 | 0.45 | >jgi\|Phyca11\|569541\|estExt2_Genewise1.C_PHYCAscaffold_330109 |
| 82 | 1 | 21290 | 114 | 151885 | 10 | 10 | 8 | 8 | 0.25 | >jgi\|Phyca11\|21290\|fgenesh1_pg.PHYCAscaffold_89_#_4 |
| 83 | 1 | 9669 | 113 | 23181 | 5 | 5 | 4 | 4 | 1.07 | >jgi\|Phyca11\|9669\|fgenesh1_pm.PHYCAscaffold_40_#_57 |
| 84 | 1 | 504348 | 113 | 29796 | 4 | 4 | 3 | 3 | 0.53 | >jgi\|Phyca11\|504348\|fgenesh2_kg.PHYCAscaffold_7_#_100_#_gi\|189084964\|gb\|BT032496.1\| |
| 36 | 2 | 530396 | 111 | 90922 | 3 | 3 | 3 | 3 | 0.15 | >jgi\|Phyca11\|530396\|estExt2_fgenesh1_pm.C_PHYCAscaffold_630021 |
| 86 | 1 | 508289 | 110 | 17173 | 6 | 6 | 5 | 5 | 2.37 | >jgi\|Phyca11\|508289\|fgenesh2_kg.PHYCAscaffold_33_#_148_#_4101647:5 |
| 87 | 1 | 20647 | 109 | 32057 | 4 | 4 | 3 | 3 | 0.49 | >jgi\|Phyca11\|20647\|fgenesh1_pg.PHYCAscaffold_70_#_2 |
| 88 | 1 | 535625 | 108 | 223563 | 7 | 7 | 7 | 7 | 0.14 | >jgi\|Phyca11\|535625\|estExt2_fgenesh1_pg.C_PHYCAscaffold_390039 |
| 89 | 1 | 14958 | 105 | 141176 | 4 | 4 | 3 | 3 | 0.1 | >jgi\|Phyca11\|14958\|fgenesh1_pg.PHYCAscaffold_10_#_104 |
| 90 | 1 | 506576 | 104 | 72048 | 5 | 5 | 4 | 4 | 0.27 | >jgi\|Phyca11\|506576\|fgenesh2_kg.PHYCAscaffold_20_#_122_#_Contig362.1 |
| 91 | 1 | 542625 | 104 | 62190 | 3 | 3 | 3 | 3 | 0.23 | >jgi\|Phyca11\|542625\|estExt2_Genewise1Plus.C_PHYCAscaffold_90614 |
| 92 | 1 | 527083 | 103 | 43530 | 7 | 7 | 4 | 4 | 0.48 | >jgi\|Phyca11\|527083\|estExt2_fgenesh1_pm.C_PHYCAscaffold_150137 |
| 93 | 1 | 116914 | 103 | 30361 | 8 | 8 | 7 | 7 | 1.65 | >jgi\|Phyca11\|116914\|e_gw1.32.324.1 |
| 94 | 1 | 6173 | 103 | 109251 | 9 | 9 | 8 | 8 | 0.42 | >jgi\|Phyca11\|6173\|fgenesh1_pm.PHYCAscaffold_10_#_20 |
| 95 | 1 | 504594 | 101 | 48837 | 3 | 3 | 3 | 3 | 0.3 | >jgi\|Phyca11\|504594\|fgenesh2_kg.PHYCAscaffold_8_#_163_#_Contig1207.1 |
| 28 | 7 | 7659 | 99 | 158087 | 6 | 6 | 4 | 4 | 0.11 | >jgi\|Phyca11\|7659\|fgenesh1_pm.PHYCAscaffold_21_#_32 |
| 96 | 1 | 554070 | 99 | 135803 | 4 | 4 | 4 | 4 | 0.13 | >jgi\|Phyca11\|554070\|estExt2_Genewise1Plus.C_PHYCAscaffold_580178 |
| 97 | 1 | 509634 | 99 | 103749 | 3 | 3 | 3 | 3 | 0.13 | >jgi\|Phyca11\|509634\|fgenesh2_kg.PHYCAscaffold_48_#_37_#_gi\|189084256\|gb\|BT031772.1\| |
| 57 | 2 | 132086 | 97 | 42182 | 6 | 6 | 5 | 5 | 0.65 | >jgi\|Phyca11\|132086\|e_gw1.132.17.1 |
| 98 | 1 | 507227 | 96 | 53724 | 4 | 4 | 4 | 4 | 0.37 | >jgi\|Phyca11\|507227\|fgenesh2_kg.PHYCAscaffold_26_#_30_#_Contig197.1 |
| 99 | 1 | 566170 | 96 | 30514 | 6 | 6 | 4 | 4 | 0.74 | >jgi\|Phyca11\|566170\|estExt2_Genewise1.C_PHYCAscaffold_200232 |
| 100 | 1 | 576097 | 96 | 49574 | 3 | 3 | 3 | 3 | 0.29 | >jgi\|Phyca11\|576097\|estExt2_Genewise1.C_PHYCAscaffold_840019 |
| 101 | 1 | 507331 | 95 | 29680 | 7 | 7 | 5 | 5 | 1.04 | >jgi\|Phyca11\|507331\|fgenesh2_kg.PHYCAscaffold_27_#_5_#_gi\|189084902\|gb\|BT032433.1\| |
| 102 | 1 | 545771 | 93 | 116345 | 5 | 5 | 5 | 5 | 0.2 | >jgi\|Phyca11\|545771\|estExt2_Genewise1Plus.C_PHYCAscaffold_190083 |
| 103 | 1 | 120320 | 93 | 124349 | 6 | 6 | 5 | 5 | 0.19 | >jgi\|Phyca11\|120320\|e_gw1.41.116.1 |
| 104 | 1 | 574490 | 92 | 129810 | 7 | 7 | 6 | 6 | 0.22 | >jgi\|Phyca11\|574490\|estExt2_Genewise1.C_PHYCAscaffold_630028 |
| 105 | 1 | 510571 | 92 | 124747 | 7 | 7 | 7 | 7 | 0.27 | >jgi\|Phyca11\|510571\|fgenesh2_kg.PHYCAscaffold_63_#_10_#_gi\|189084688\|gb\|BT032204.1\| |
| 106 | 1 | 503902 | 90 | 35315 | 4 | 4 | 3 | 3 | 0.43 | >jgi\|Phyca11\|503902\|fgenesh2_kg.PHYCAscaffold_5_#_66_#_Contig14.1 |
| 107 | 1 | 532200 | 89 | 114748 | 2 | 2 | 2 | 2 | 0.08 | >jgi\|Phyca11\|532200\|estExt2_fgenesh1_pg.C_PHYCAscaffold_40145 |
| 109 | 1 | 507472 | 88 | 24177 | 3 | 3 | 3 | 3 | 0.69 | >jgi\|Phyca11\|507472\|fgenesh2_kg.PHYCAscaffold_27_#_146_#_Contig84.1 |
| 110 | 1 | 545598 | 87 | 111192 | 9 | 9 | 9 | 9 | 0.41 | >jgi\|Phyca11\|545598\|estExt2_Genewise1Plus.C_PHYCAscaffold_180327 |
| 111 | 1 | 126045 | 85 | 82427 | 3 | 3 | 3 | 3 | 0.17 | >jgi\|Phyca11\|126045\|e_gw1.61.41.1 |
| 112 | 1 | 118185 | 84 | 222850 | 3 | 3 | 3 | 3 | 0.06 | >jgi\|Phyca11\|118185\|e_gw1.35.65.1 |
| 113 | 1 | 506902 | 84 | 27776 | 2 | 2 | 2 | 2 | 0.36 | >jgi\|Phyca11\|506902\|fgenesh2_kg.PHYCAscaffold_23_#_33_#_gi\|189084893\|gb\|BT032424.1\| |
| 114 | 1 | 113262 | 84 | 45975 | 3 | 3 | 2 | 2 | 0.2 | >jgi\|Phyca11\|113262\|e_gw1.23.227.1 |
| 115 | 1 | 540959 | 82 | 21538 | 5 | 5 | 4 | 4 | 1.18 | >jgi\|Phyca11\|540959\|estExt2_Genewise1Plus.C_PHYCAscaffold_50907 |
| 116 | 1 | 561150 | 82 | 65782 | 4 | 4 | 4 | 4 | 0.3 | >jgi\|Phyca11\|561150\|estExt2_Genewise1.C_PHYCAscaffold_60312 |
| 117 | 1 | 550410 | 81 | 73974 | 5 | 5 | 4 | 4 | 0.26 | >jgi\|Phyca11\|550410\|estExt2_Genewise1Plus.C_PHYCAscaffold_370115 |
| 118 | 1 | 502925 | 81 | 12976 | 2 | 2 | 2 | 2 | 0.89 | >jgi\|Phyca11\|502925\|fgenesh2_kg.PHYCAscaffold_2_#_41_#_Contig2039.1 |
| 119 | 1 | 535506 | 79 | 74784 | 5 | 5 | 5 | 5 | 0.33 | >jgi\|Phyca11\|535506\|estExt2_fgenesh1_pg.C_PHYCAscaffold_370049 |
| 120 | 1 | 529559 | 79 | 122629 | 5 | 5 | 5 | 5 | 0.19 | >jgi\|Phyca11\|529559\|estExt2_fgenesh1_pm.C_PHYCAscaffold_460012 |
| 121 | 1 | 114981 | 79 | 28230 | 2 | 2 | 2 | 2 | 0.35 | >jgi\|Phyca11\|114981\|e_gw1.27.273.1 |
| 122 | 1 | 537506 | 79 | 25792 | 2 | 2 | 2 | 2 | 0.39 | >jgi\|Phyca11\|537506\|estExt2_fgenesh1_pg.C_PHYCAscaffold_950002 |
| 123 | 1 | 544770 | 79 | 108578 | 6 | 6 | 4 | 4 | 0.17 | >jgi\|Phyca11\|544770\|estExt2_Genewise1Plus.C_PHYCAscaffold_150540 |
| 124 | 1 | 510696 | 78 | 72798 | 2 | 2 | 2 | 2 | 0.12 | >jgi\|Phyca11\|510696\|fgenesh2_kg.PHYCAscaffold_65_#_36_#_4098202:1 |
| 125 | 1 | 111740 | 77 | 90538 | 4 | 4 | 4 | 4 | 0.21 | >jgi\|Phyca11\|111740\|e_gw1.20.14.1 |
| 126 | 1 | 5622 | 77 | 193584 | 3 | 3 | 3 | 3 | 0.07 | >jgi\|Phyca11\|5622\|fgenesh1_pm.PHYCAscaffold_6_#_142 |
| 117 | 2 | 508653 | 75 | 74131 | 6 | 6 | 5 | 5 | 0.33 | >jgi\|Phyca11\|508653\|fgenesh2_kg.PHYCAscaffold_37_#_28_#_Contig1670.1 |
| 127 | 1 | 562791 | 75 | 62172 | 4 | 4 | 3 | 3 | 0.23 | >jgi\|Phyca11\|562791\|estExt2_Genewise1.C_PHYCAscaffold_100242 |
| 128 | 1 | 10801 | 74 | 143858 | 3 | 3 | 3 | 3 | 0.09 | >jgi\|Phyca11\|10801\|fgenesh1_pm.PHYCAscaffold_55_#_22 |
| 129 | 1 | 562574 | 74 | 58435 | 6 | 6 | 5 | 5 | 0.44 | >jgi\|Phyca11\|562574\|estExt2_Genewise1.C_PHYCAscaffold_90628 |
| 130 | 1 | 528038 | 73 | 193379 | 4 | 4 | 3 | 3 | 0.07 | >jgi\|Phyca11\|528038\|estExt2_fgenesh1_pm.C_PHYCAscaffold_260026 |
| 131 | 1 | 505083 | 73 | 69264 | 2 | 2 | 2 | 2 | 0.13 | >jgi\|Phyca11\|505083\|fgenesh2_kg.PHYCAscaffold_11_#_29_#_gi\|189084466\|gb\|BT031982.1\| |
| 132 | 1 | 565311 | 72 | 60786 | 4 | 4 | 3 | 3 | 0.23 | >jgi\|Phyca11\|565311\|estExt2_Genewise1.C_PHYCAscaffold_170498 |
| 133 | 1 | 533182 | 71 | 114501 | 5 | 5 | 4 | 4 | 0.16 | >jgi\|Phyca11\|533182\|estExt2_fgenesh1_pg.C_PHYCAscaffold_110057 |
| 134 | 1 | 550454 | 70 | 115608 | 5 | 5 | 5 | 5 | 0.2 | >jgi\|Phyca11\|550454\|estExt2_Genewise1Plus.C_PHYCAscaffold_370174 |
| 135 | 1 | 12430 | 69 | 101309 | 4 | 4 | 3 | 3 | 0.13 | >jgi\|Phyca11\|12430\|fgenesh1_pm.PHYCAscaffold_125_#_1 |
| 136 | 1 | 507760 | 69 | 27768 | 2 | 2 | 2 | 2 | 0.36 | >jgi\|Phyca11\|507760\|fgenesh2_kg.PHYCAscaffold_30_#_29_#_Contig16.1 |
| 137 | 1 | 507516 | 68 | 34484 | 2 | 2 | 2 | 2 | 0.28 | >jgi\|Phyca11\|507516\|fgenesh2_kg.PHYCAscaffold_28_#_23_#_Contig24.1 |
| 138 | 1 | 9769 | 68 | 21549 | 2 | 2 | 2 | 2 | 0.48 | >jgi\|Phyca11\|9769\|fgenesh1_pm.PHYCAscaffold_41_#_63 |
| 47 | 2 | 526928 | 67 | 161520 | 2 | 2 | 2 | 2 | 0.05 | >jgi\|Phyca11\|526928\|estExt2_fgenesh1_pm.C_PHYCAscaffold_140061 |
| 139 | 1 | 545922 | 67 | 100176 | 6 | 6 | 6 | 6 | 0.29 | >jgi\|Phyca11\|545922\|estExt2_Genewise1Plus.C_PHYCAscaffold_190330 |
| 140 | 1 | 503358 | 67 | 51975 | 3 | 3 | 3 | 3 | 0.28 | >jgi\|Phyca11\|503358\|fgenesh2_kg.PHYCAscaffold_3_#_223_#_Contig173.1 |
| 141 | 1 | 505375 | 66 | 17847 | 4 | 4 | 4 | 4 | 1.55 | >jgi\|Phyca11\|505375\|fgenesh2_kg.PHYCAscaffold_13_#_25_#_4098936:2 |
| 142 | 1 | 526098 | 66 | 61646 | 4 | 4 | 4 | 4 | 0.32 | >jgi\|Phyca11\|526098\|estExt2_fgenesh1_pm.C_PHYCAscaffold_70057 |
| 143 | 1 | 538799 | 66 | 111167 | 4 | 4 | 4 | 4 | 0.17 | >jgi\|Phyca11\|538799\|estExt2_Genewise1Plus.C_PHYCAscaffold_20456 |
| 145 | 1 | 534661 | 65 | 136091 | 3 | 3 | 3 | 3 | 0.1 | >jgi\|Phyca11\|534661\|estExt2_fgenesh1_pg.C_PHYCAscaffold_260096 |
| 146 | 1 | 509548 | 65 | 90904 | 4 | 4 | 4 | 4 | 0.21 | >jgi\|Phyca11\|509548\|fgenesh2_kg.PHYCAscaffold_47_#_39_#_gi\|189084335\|gb\|BT031851.1\| |
| 28 | 8 | 504947 | 64 | 147031 | 4 | 4 | 2 | 2 | 0.06 | >jgi\|Phyca11\|504947\|fgenesh2_kg.PHYCAscaffold_10_#_107_#_Contig62.1 |
| 147 | 1 | 531975 | 64 | 72070 | 3 | 3 | 3 | 3 | 0.19 | >jgi\|Phyca11\|531975\|estExt2_fgenesh1_pg.C_PHYCAscaffold_30139 |
| 148 | 1 | 510504 | 64 | 38812 | 3 | 3 | 3 | 3 | 0.39 | >jgi\|Phyca11\|510504\|fgenesh2_kg.PHYCAscaffold_61_#_37_#_4101945:41 |
| 28 | 9 | 11692 | 63 | 140701 | 4 | 4 | 2 | 2 | 0.06 | >jgi\|Phyca11\|11692\|fgenesh1_pm.PHYCAscaffold_78_#_16 |
| 149 | 1 | 504126 | 63 | 122043 | 4 | 4 | 3 | 3 | 0.11 | >jgi\|Phyca11\|504126\|fgenesh2_kg.PHYCAscaffold_6_#_43_#_Contig2838.1 |
| 150 | 1 | 559135 | 62 | 119707 | 4 | 4 | 4 | 4 | 0.15 | >jgi\|Phyca11\|559135\|estExt2_Genewise1.C_PHYCAscaffold_21155 |
| 151 | 1 | 508065 | 60 | 61901 | 3 | 3 | 3 | 3 | 0.23 | >jgi\|Phyca11\|508065\|fgenesh2_kg.PHYCAscaffold_32_#_24_#_Contig137.1 |
| 152 | 1 | 525911 | 59 | 76975 | 1 | 1 | 1 | 1 | 0.06 | >jgi\|Phyca11\|525911\|estExt2_fgenesh1_pm.C_PHYCAscaffold_50244 |
| 153 | 1 | 507425 | 59 | 21112 | 5 | 5 | 3 | 3 | 0.82 | >jgi\|Phyca11\|507425\|fgenesh2_kg.PHYCAscaffold_27_#_99_#_4096835:1 |
| 155 | 1 | 568092 | 59 | 192568 | 2 | 2 | 2 | 2 | 0.05 | >jgi\|Phyca11\|568092\|estExt2_Genewise1.C_PHYCAscaffold_270331 |
| 156 | 1 | 104335 | 59 | 71675 | 5 | 5 | 4 | 4 | 0.27 | >jgi\|Phyca11\|104335\|e_gw1.9.161.1 |
| 157 | 1 | 503123 | 59 | 44559 | 6 | 6 | 4 | 4 | 0.46 | >jgi\|Phyca11\|503123\|fgenesh2_kg.PHYCAscaffold_2_#_239_#_Contig21.3 |
| 19 | 4 | 535813 | 58 | 215723 | 4 | 4 | 4 | 4 | 0.08 | >jgi\|Phyca11\|535813\|estExt2_fgenesh1_pg.C_PHYCAscaffold_420044 |
| 158 | 1 | 507299 | 58 | 86996 | 2 | 2 | 2 | 2 | 0.1 | >jgi\|Phyca11\|507299\|fgenesh2_kg.PHYCAscaffold_26_#_102_#_Contig1340.2 |
| 159 | 1 | 544983 | 58 | 32876 | 1 | 1 | 1 | 1 | 0.14 | >jgi\|Phyca11\|544983\|estExt2_Genewise1Plus.C_PHYCAscaffold_160492 |
| 160 | 1 | 508801 | 57 | 20272 | 3 | 3 | 3 | 3 | 0.86 | >jgi\|Phyca11\|508801\|fgenesh2_kg.PHYCAscaffold_38_#_53_#_Contig95.1 |
| 161 | 1 | 502708 | 57 | 26043 | 2 | 2 | 2 | 2 | 0.38 | >jgi\|Phyca11\|502708\|fgenesh2_kg.PHYCAscaffold_1_#_168_#_Contig881.1 |
| 162 | 1 | 504193 | 57 | 150606 | 3 | 3 | 3 | 3 | 0.09 | >jgi\|Phyca11\|504193\|fgenesh2_kg.PHYCAscaffold_6_#_110_#_gi\|189084605\|gb\|BT032121.1\| |
| 28 | 10 | 106036 | 56 | 140672 | 4 | 4 | 2 | 2 | 0.06 | >jgi\|Phyca11\|106036\|e_gw1.11.372.1 |
| 163 | 1 | 15064 | 55 | 94252 | 2 | 2 | 1 | 1 | 0.05 | >jgi\|Phyca11\|15064\|fgenesh1_pg.PHYCAscaffold_11_#_50 |
| 164 | 1 | 503099 | 54 | 52799 | 1 | 1 | 1 | 1 | 0.08 | >jgi\|Phyca11\|503099\|fgenesh2_kg.PHYCAscaffold_2_#_215_#_4096614:2 |
| 165 | 1 | 574456 | 54 | 85938 | 2 | 2 | 2 | 2 | 0.1 | >jgi\|Phyca11\|574456\|estExt2_Genewise1.C_PHYCAscaffold_620176 |
| 166 | 1 | 503898 | 54 | 14775 | 2 | 2 | 2 | 2 | 0.76 | >jgi\|Phyca11\|503898\|fgenesh2_kg.PHYCAscaffold_5_#_62_#_gi\|189083883\|gb\|BT031399.1\| |
| 53 | 2 | 54680 | 53 | 83997 | 2 | 2 | 2 | 2 | 0.11 | >jgi\|Phyca11\|54680\|gw1.21.144.1 |
| 167 | 1 | 106526 | 53 | 99721 | 2 | 2 | 2 | 2 | 0.09 | >jgi\|Phyca11\|106526\|e_gw1.12.724.1 |
| 168 | 1 | 552811 | 53 | 61340 | 1 | 1 | 1 | 1 | 0.07 | >jgi\|Phyca11\|552811\|estExt2_Genewise1Plus.C_PHYCAscaffold_490242 |
| 169 | 1 | 537849 | 53 | 31391 | 1 | 1 | 1 | 1 | 0.14 | >jgi\|Phyca11\|537849\|estExt2_fgenesh1_pg.C_PHYCAscaffold_3510001 |
| 170 | 1 | 504902 | 52 | 24490 | 1 | 1 | 1 | 1 | 0.19 | >jgi\|Phyca11\|504902\|fgenesh2_kg.PHYCAscaffold_10_#_62_#_4096919:1 |
| 171 | 1 | 532433 | 52 | 85584 | 2 | 2 | 2 | 2 | 0.1 | >jgi\|Phyca11\|532433\|estExt2_fgenesh1_pg.C_PHYCAscaffold_50170 |
| 172 | 1 | 109031 | 52 | 82587 | 1 | 1 | 1 | 1 | 0.05 | >jgi\|Phyca11\|109031\|e_gw1.16.193.1 |
| 173 | 1 | 539778 | 51 | 88487 | 2 | 2 | 2 | 2 | 0.1 | >jgi\|Phyca11\|539778\|estExt2_Genewise1Plus.C_PHYCAscaffold_40038 |
| 174 | 1 | 551381 | 51 | 51505 | 2 | 2 | 2 | 2 | 0.18 | >jgi\|Phyca11\|551381\|estExt2_Genewise1Plus.C_PHYCAscaffold_420020 |
| 175 | 1 | 106472 | 51 | 96852 | 4 | 4 | 4 | 4 | 0.19 | >jgi\|Phyca11\|106472\|e_gw1.12.486.1 |
| 176 | 1 | 564766 | 51 | 49103 | 1 | 1 | 1 | 1 | 0.09 | >jgi\|Phyca11\|564766\|estExt2_Genewise1.C_PHYCAscaffold_150604 |
| 177 | 1 | 124163 | 51 | 11399 | 1 | 1 | 1 | 1 | 0.44 | >jgi\|Phyca11\|124163\|e_gw1.53.42.1 |
| 178 | 1 | 4583 | 51 | 123284 | 1 | 1 | 1 | 1 | 0.04 | >jgi\|Phyca11\|4583\|fgenesh1_pm.PHYCAscaffold_2_#_179 |
| 179 | 1 | 510591 | 51 | 114255 | 2 | 2 | 2 | 2 | 0.08 | >jgi\|Phyca11\|510591\|fgenesh2_kg.PHYCAscaffold_63_#_30_#_Contig771.1 |
| 180 | 1 | 504469 | 50 | 23161 | 2 | 2 | 2 | 2 | 0.44 | >jgi\|Phyca11\|504469\|fgenesh2_kg.PHYCAscaffold_8_#_38_#_4101945:176 |
| 181 | 1 | 505233 | 50 | 20821 | 3 | 3 | 2 | 2 | 0.5 | >jgi\|Phyca11\|505233\|fgenesh2_kg.PHYCAscaffold_12_#_21_#_4100316:9 |
| 182 | 1 | 571228 | 50 | 47998 | 1 | 1 | 1 | 1 | 0.09 | >jgi\|Phyca11\|571228\|estExt2_Genewise1.C_PHYCAscaffold_410155 |
| 183 | 1 | 118774 | 49 | 107154 | 4 | 4 | 4 | 4 | 0.17 | >jgi\|Phyca11\|118774\|e_gw1.37.8.1 |
| 184 | 1 | 505046 | 48 | 80584 | 4 | 4 | 4 | 4 | 0.24 | >jgi\|Phyca11\|505046\|fgenesh2_kg.PHYCAscaffold_10_#_206_#_4097622:1 |
| 185 | 1 | 506983 | 48 | 22216 | 3 | 3 | 3 | 3 | 0.77 | >jgi\|Phyca11\|506983\|fgenesh2_kg.PHYCAscaffold_23_#_114_#_Contig3703.1 |
| 186 | 1 | 504258 | 48 | 22912 | 1 | 1 | 1 | 1 | 0.2 | >jgi\|Phyca11\|504258\|fgenesh2_kg.PHYCAscaffold_7_#_10_#_4099433:1 |
| 187 | 1 | 121937 | 48 | 61567 | 2 | 2 | 1 | 1 | 0.07 | >jgi\|Phyca11\|121937\|e_gw1.46.430.1 |
| 120 | 2 | 558677 | 47 | 99339 | 3 | 3 | 3 | 3 | 0.14 | >jgi\|Phyca11\|558677\|estExt2_Genewise1.C_PHYCAscaffold_20357 |
| 188 | 1 | 528395 | 47 | 333389 | 1 | 1 | 1 | 1 | 0.01 | >jgi\|Phyca11\|528395\|estExt2_fgenesh1_pm.C_PHYCAscaffold_290091 |
| 189 | 1 | 79187 | 47 | 26712 | 1 | 1 | 1 | 1 | 0.17 | >jgi\|Phyca11\|79187\|gw1.3.1012.1 |
| 190 | 1 | 506857 | 47 | 75869 | 2 | 2 | 2 | 2 | 0.12 | >jgi\|Phyca11\|506857\|fgenesh2_kg.PHYCAscaffold_22_#_96_#_gi\|189084480\|gb\|BT031996.1\| |
| 191 | 1 | 539189 | 47 | 16536 | 2 | 2 | 2 | 2 | 0.66 | >jgi\|Phyca11\|539189\|estExt2_Genewise1Plus.C_PHYCAscaffold_30046 |
| 192 | 1 | 507292 | 46 | 28550 | 2 | 2 | 2 | 2 | 0.34 | >jgi\|Phyca11\|507292\|fgenesh2_kg.PHYCAscaffold_26_#_95_#_4096992:5 |
| 193 | 1 | 96024 | 46 | 36676 | 1 | 1 | 1 | 1 | 0.12 | >jgi\|Phyca11\|96024\|e_gw1.1.1208.1 |
| 194 | 1 | 510172 | 46 | 43700 | 2 | 2 | 2 | 2 | 0.21 | >jgi\|Phyca11\|510172\|fgenesh2_kg.PHYCAscaffold_54_#_66_#_4101945:150 |
| 195 | 1 | 506540 | 46 | 12303 | 1 | 1 | 1 | 1 | 0.4 | >jgi\|Phyca11\|506540\|fgenesh2_kg.PHYCAscaffold_20_#_86_#_Contig164.1 |
| 196 | 1 | 504531 | 45 | 15100 | 2 | 2 | 2 | 2 | 0.74 | >jgi\|Phyca11\|504531\|fgenesh2_kg.PHYCAscaffold_8_#_100_#_4101945:12 |
| 197 | 1 | 569720 | 45 | 58735 | 3 | 3 | 3 | 3 | 0.24 | >jgi\|Phyca11\|569720\|estExt2_Genewise1.C_PHYCAscaffold_330350 |
| 198 | 1 | 34030 | 45 | 111847 | 4 | 4 | 3 | 3 | 0.12 | >jgi\|Phyca11\|34030\|gw1.27.9.1 |
| 199 | 1 | 559703 | 45 | 13273 | 2 | 2 | 1 | 1 | 0.37 | >jgi\|Phyca11\|559703\|estExt2_Genewise1.C_PHYCAscaffold_31039 |
| 200 | 1 | 539116 | 45 | 56269 | 3 | 3 | 3 | 3 | 0.25 | >jgi\|Phyca11\|539116\|estExt2_Genewise1Plus.C_PHYCAscaffold_20989 |
| 201 | 1 | 531963 | 44 | 89084 | 2 | 2 | 2 | 2 | 0.1 | >jgi\|Phyca11\|531963\|estExt2_fgenesh1_pg.C_PHYCAscaffold_30124 |
| 202 | 1 | 506531 | 44 | 68497 | 3 | 3 | 3 | 3 | 0.21 | >jgi\|Phyca11\|506531\|fgenesh2_kg.PHYCAscaffold_20_#_77_#_Contig273.1 |
| 203 | 1 | 21359 | 44 | 63609 | 2 | 2 | 2 | 2 | 0.14 | >jgi\|Phyca11\|21359\|fgenesh1_pg.PHYCAscaffold_92_#_14 |
| 204 | 1 | 511668 | 44 | 47952 | 1 | 1 | 1 | 1 | 0.09 | >jgi\|Phyca11\|511668\|fgenesh2_kg.PHYCAscaffold_94_#_19_#_4101945:62 |
| 205 | 1 | 114899 | 44 | 191346 | 2 | 2 | 2 | 2 | 0.05 | >jgi\|Phyca11\|114899\|e_gw1.27.1.1 |
| 206 | 1 | 527239 | 43 | 123962 | 3 | 3 | 3 | 3 | 0.11 | >jgi\|Phyca11\|527239\|estExt2_fgenesh1_pm.C_PHYCAscaffold_170121 |
| 140 | 2 | 503356 | 42 | 49536 | 2 | 2 | 2 | 2 | 0.19 | >jgi\|Phyca11\|503356\|fgenesh2_kg.PHYCAscaffold_3_#_221_#_4099680:2 |
| 207 | 1 | 553648 | 42 | 54839 | 1 | 1 | 1 | 1 | 0.08 | >jgi\|Phyca11\|553648\|estExt2_Genewise1Plus.C_PHYCAscaffold_540171 |
| 208 | 1 | 527845 | 42 | 43946 | 1 | 1 | 1 | 1 | 0.1 | >jgi\|Phyca11\|527845\|estExt2_fgenesh1_pm.C_PHYCAscaffold_230070 |
| 209 | 1 | 535286 | 42 | 57740 | 1 | 1 | 1 | 1 | 0.08 | >jgi\|Phyca11\|535286\|estExt2_fgenesh1_pg.C_PHYCAscaffold_330120 |
| 210 | 1 | 577778 | 42 | 33335 | 1 | 1 | 1 | 1 | 0.14 | >jgi\|Phyca11\|577778\|estExt2_Genewise1.C_PHYCAscaffold_3970001 |
| 211 | 1 | 533155 | 41 | 182767 | 3 | 3 | 3 | 3 | 0.07 | >jgi\|Phyca11\|533155\|estExt2_fgenesh1_pg.C_PHYCAscaffold_110021 |
| 212 | 1 | 503545 | 41 | 98917 | 1 | 1 | 1 | 1 | 0.04 | >jgi\|Phyca11\|503545\|fgenesh2_kg.PHYCAscaffold_4_#_84_#_Contig5.2 |
| 213 | 1 | 19814 | 41 | 73726 | 1 | 1 | 1 | 1 | 0.06 | >jgi\|Phyca11\|19814\|fgenesh1_pg.PHYCAscaffold_52_#_66 |
| 214 | 1 | 506922 | 41 | 14320 | 1 | 1 | 1 | 1 | 0.34 | >jgi\|Phyca11\|506922\|fgenesh2_kg.PHYCAscaffold_23_#_53_#_4101134:1 |
| 70 | 2 | 53999 | 40 | 109401 | 3 | 3 | 3 | 3 | 0.12 | >jgi\|Phyca11\|53999\|gw1.13.151.1 |
| 215 | 1 | 533060 | 40 | 90951 | 3 | 3 | 2 | 2 | 0.1 | >jgi\|Phyca11\|533060\|estExt2_fgenesh1_pg.C_PHYCAscaffold_100047 |
| 216 | 1 | 510818 | 40 | 75956 | 1 | 1 | 1 | 1 | 0.06 | >jgi\|Phyca11\|510818\|fgenesh2_kg.PHYCAscaffold_67_#_29_#_Contig288.2 |
| 217 | 1 | 506742 | 40 | 79728 | 2 | 2 | 2 | 2 | 0.11 | >jgi\|Phyca11\|506742\|fgenesh2_kg.PHYCAscaffold_21_#_114_#_Contig1308.1 |
| 218 | 1 | 538160 | 40 | 6551 | 1 | 1 | 1 | 1 | 0.85 | >jgi\|Phyca11\|538160\|estExt2_Genewise1Plus.C_PHYCAscaffold_10551 |
| 219 | 1 | 509774 | 39 | 85263 | 4 | 4 | 4 | 4 | 0.22 | >jgi\|Phyca11\|509774\|fgenesh2_kg.PHYCAscaffold_49_#_66_#_Contig174.1 |
| 220 | 1 | 7598 | 39 | 17660 | 1 | 1 | 1 | 1 | 0.27 | >jgi\|Phyca11\|7598\|fgenesh1_pm.PHYCAscaffold_20_#_113 |
| 221 | 1 | 529934 | 39 | 121660 | 3 | 3 | 3 | 3 | 0.11 | >jgi\|Phyca11\|529934\|estExt2_fgenesh1_pm.C_PHYCAscaffold_510045 |
| 222 | 1 | 565700 | 39 | 55309 | 2 | 2 | 2 | 2 | 0.17 | >jgi\|Phyca11\|565700\|estExt2_Genewise1.C_PHYCAscaffold_190016 |
| 223 | 1 | 507479 | 39 | 44888 | 1 | 1 | 1 | 1 | 0.1 | >jgi\|Phyca11\|507479\|fgenesh2_kg.PHYCAscaffold_27_#_153_#_4098657:2 |
| 224 | 1 | 529283 | 39 | 60530 | 2 | 2 | 2 | 2 | 0.15 | >jgi\|Phyca11\|529283\|estExt2_fgenesh1_pm.C_PHYCAscaffold_410034 |
| 225 | 1 | 562935 | 39 | 110180 | 1 | 1 | 1 | 1 | 0.04 | >jgi\|Phyca11\|562935\|estExt2_Genewise1.C_PHYCAscaffold_100543 |
| 226 | 1 | 10889 | 39 | 36420 | 1 | 1 | 1 | 1 | 0.12 | >jgi\|Phyca11\|10889\|fgenesh1_pm.PHYCAscaffold_58_#_1 |
| 227 | 1 | 508469 | 39 | 21033 | 1 | 1 | 1 | 1 | 0.22 | >jgi\|Phyca11\|508469\|fgenesh2_kg.PHYCAscaffold_35_#_52_#_4098364:1 |
| 228 | 1 | 526703 | 39 | 111673 | 1 | 1 | 1 | 1 | 0.04 | >jgi\|Phyca11\|526703\|estExt2_fgenesh1_pm.C_PHYCAscaffold_120048 |
| 229 | 1 | 99958 | 38 | 49559 | 1 | 1 | 1 | 1 | 0.09 | >jgi\|Phyca11\|99958\|e_gw1.4.612.1 |
| 230 | 1 | 503568 | 38 | 49308 | 2 | 2 | 2 | 2 | 0.19 | >jgi\|Phyca11\|503568\|fgenesh2_kg.PHYCAscaffold_4_#_107_#_4097947:1 |
| 231 | 1 | 510914 | 38 | 17277 | 3 | 3 | 3 | 3 | 1.06 | >jgi\|Phyca11\|510914\|fgenesh2_kg.PHYCAscaffold_71_#_14_#_gi\|189083867\|gb\|BT031383.1\| |
| 232 | 1 | 21516 | 38 | 40377 | 1 | 1 | 1 | 1 | 0.11 | >jgi\|Phyca11\|21516\|fgenesh1_pg.PHYCAscaffold_99_#_12 |
| 233 | 1 | 505405 | 38 | 46756 | 2 | 2 | 2 | 2 | 0.2 | >jgi\|Phyca11\|505405\|fgenesh2_kg.PHYCAscaffold_13_#_55_#_4101849:4 |
| 234 | 1 | 528366 | 38 | 101027 | 2 | 2 | 2 | 2 | 0.09 | >jgi\|Phyca11\|528366\|estExt2_fgenesh1_pm.C_PHYCAscaffold_290052 |
| 235 | 1 | 8284 | 38 | 59975 | 1 | 1 | 1 | 1 | 0.07 | >jgi\|Phyca11\|8284\|fgenesh1_pm.PHYCAscaffold_27_#_30 |
| 237 | 1 | 534321 | 37 | 65568 | 3 | 3 | 3 | 3 | 0.22 | >jgi\|Phyca11\|534321\|estExt2_fgenesh1_pg.C_PHYCAscaffold_220091 |
| 238 | 1 | 559789 | 37 | 115359 | 1 | 1 | 1 | 1 | 0.04 | >jgi\|Phyca11\|559789\|estExt2_Genewise1.C_PHYCAscaffold_40097 |
| 239 | 1 | 510339 | 37 | 24199 | 1 | 1 | 1 | 1 | 0.19 | >jgi\|Phyca11\|510339\|fgenesh2_kg.PHYCAscaffold_58_#_52_#_Contig5159.1 |
| 240 | 1 | 116963 | 36 | 95001 | 1 | 1 | 1 | 1 | 0.05 | >jgi\|Phyca11\|116963\|e_gw1.32.12.1 |
| 241 | 1 | 17248 | 36 | 23352 | 2 | 2 | 2 | 2 | 0.44 | >jgi\|Phyca11\|17248\|fgenesh1_pg.PHYCAscaffold_26_#_51 |
| 242 | 1 | 510582 | 36 | 52105 | 3 | 3 | 3 | 3 | 0.28 | >jgi\|Phyca11\|510582\|fgenesh2_kg.PHYCAscaffold_63_#_21_#_Contig2224.1 |
| 243 | 1 | 511272 | 36 | 50402 | 1 | 1 | 1 | 1 | 0.09 | >jgi\|Phyca11\|511272\|fgenesh2_kg.PHYCAscaffold_80_#_16_#_4101869:2 |
| 244 | 1 | 504523 | 36 | 36578 | 2 | 2 | 2 | 2 | 0.26 | >jgi\|Phyca11\|504523\|fgenesh2_kg.PHYCAscaffold_8_#_92_#_4101945:54 |
| 245 | 1 | 503414 | 36 | 55335 | 2 | 2 | 1 | 1 | 0.17 | >jgi\|Phyca11\|503414\|fgenesh2_kg.PHYCAscaffold_3_#_279_#_Contig81.1 |
| 246 | 1 | 52778 | 36 | 21158 | 4 | 4 | 1 | 1 | 0.22 | >jgi\|Phyca11\|52778\|gw1.859.2.1 |
| 247 | 1 | 545320 | 36 | 95221 | 2 | 2 | 2 | 2 | 0.09 | >jgi\|Phyca11\|545320\|estExt2_Genewise1Plus.C_PHYCAscaffold_170465 |
| 248 | 1 | 508178 | 36 | 46273 | 1 | 1 | 1 | 1 | 0.1 | >jgi\|Phyca11\|508178\|fgenesh2_kg.PHYCAscaffold_33_#_37_#_Contig486.1 |
| 249 | 1 | 534354 | 36 | 105373 | 1 | 1 | 1 | 1 | 0.04 | >jgi\|Phyca11\|534354\|estExt2_fgenesh1_pg.C_PHYCAscaffold_230001 |
| 250 | 1 | 7927 | 36 | 13927 | 1 | 1 | 1 | 1 | 0.35 | >jgi\|Phyca11\|7927\|fgenesh1_pm.PHYCAscaffold_23_#_75 |
| 251 | 1 | 571672 | 36 | 299795 | 4 | 4 | 4 | 4 | 0.06 | >jgi\|Phyca11\|571672\|estExt2_Genewise1.C_PHYCAscaffold_430183 |
| 252 | 1 | 131382 | 35 | 271479 | 1 | 1 | 1 | 1 | 0.02 | >jgi\|Phyca11\|131382\|e_gw1.105.3.1 |
| 253 | 1 | 554212 | 35 | 130584 | 1 | 1 | 1 | 1 | 0.03 | >jgi\|Phyca11\|554212\|estExt2_Genewise1Plus.C_PHYCAscaffold_600025 |
| 254 | 1 | 542607 | 35 | 51338 | 2 | 2 | 2 | 2 | 0.18 | >jgi\|Phyca11\|542607\|estExt2_Genewise1Plus.C_PHYCAscaffold_90588 |
| 255 | 1 | 526447 | 35 | 54545 | 1 | 1 | 1 | 1 | 0.08 | >jgi\|Phyca11\|526447\|estExt2_fgenesh1_pm.C_PHYCAscaffold_90195 |
| 256 | 1 | 536953 | 34 | 269706 | 1 | 1 | 1 | 1 | 0.02 | >jgi\|Phyca11\|536953\|estExt2_fgenesh1_pg.C_PHYCAscaffold_680021 |
| 257 | 1 | 506723 | 34 | 38939 | 2 | 2 | 2 | 2 | 0.24 | >jgi\|Phyca11\|506723\|fgenesh2_kg.PHYCAscaffold_21_#_95_#_4101605:2 |
| 258 | 1 | 16801 | 34 | 59966 | 2 | 2 | 2 | 2 | 0.15 | >jgi\|Phyca11\|16801\|fgenesh1_pg.PHYCAscaffold_22_#_80 |
| 259 | 1 | 502842 | 34 | 47240 | 1 | 1 | 1 | 1 | 0.09 | >jgi\|Phyca11\|502842\|fgenesh2_kg.PHYCAscaffold_1_#_302_#_4096672:1 |
| 260 | 1 | 509041 | 34 | 22172 | 2 | 2 | 2 | 2 | 0.46 | >jgi\|Phyca11\|509041\|fgenesh2_kg.PHYCAscaffold_41_#_37_#_Contig2305.1 |
| 261 | 1 | 541906 | 34 | 48490 | 2 | 2 | 2 | 2 | 0.19 | >jgi\|Phyca11\|541906\|estExt2_Genewise1Plus.C_PHYCAscaffold_80096 |
| 262 | 1 | 535457 | 34 | 21742 | 1 | 1 | 1 | 1 | 0.21 | >jgi\|Phyca11\|535457\|estExt2_fgenesh1_pg.C_PHYCAscaffold_360050 |
| 263 | 1 | 551663 | 34 | 112129 | 2 | 2 | 2 | 2 | 0.08 | >jgi\|Phyca11\|551663\|estExt2_Genewise1Plus.C_PHYCAscaffold_430135 |
| 264 | 1 | 133548 | 34 | 8927 | 1 | 1 | 1 | 1 | 0.58 | >jgi\|Phyca11\|133548\|e_gw1.548.9.1 |
| 265 | 1 | 550970 | 33 | 77497 | 2 | 2 | 1 | 1 | 0.06 | >jgi\|Phyca11\|550970\|estExt2_Genewise1Plus.C_PHYCAscaffold_400016 |
| 267 | 1 | 534326 | 33 | 99204 | 2 | 2 | 2 | 2 | 0.09 | >jgi\|Phyca11\|534326\|estExt2_fgenesh1_pg.C_PHYCAscaffold_220096 |
| 268 | 1 | 508162 | 33 | 43789 | 1 | 1 | 1 | 1 | 0.1 | >jgi\|Phyca11\|508162\|fgenesh2_kg.PHYCAscaffold_33_#_21_#_4101225:1 |
| 269 | 1 | 10281 | 33 | 51491 | 2 | 2 | 2 | 2 | 0.18 | >jgi\|Phyca11\|10281\|fgenesh1_pm.PHYCAscaffold_48_#_18 |
| 270 | 1 | 538776 | 32 | 17625 | 1 | 1 | 1 | 1 | 0.27 | >jgi\|Phyca11\|538776\|estExt2_Genewise1Plus.C_PHYCAscaffold_20418 |
| 271 | 1 | 540712 | 32 | 117259 | 1 | 1 | 1 | 1 | 0.04 | >jgi\|Phyca11\|540712\|estExt2_Genewise1Plus.C_PHYCAscaffold_50468 |
| 272 | 1 | 558170 | 32 | 47229 | 2 | 2 | 2 | 2 | 0.2 | >jgi\|Phyca11\|558170\|estExt2_Genewise1.C_PHYCAscaffold_10741 |
| 273 | 1 | 13027 | 32 | 16189 | 1 | 1 | 1 | 1 | 0.29 | >jgi\|Phyca11\|13027\|fgenesh1_pg.PHYCAscaffold_2_#_123 |
| 274 | 1 | 9799 | 32 | 55300 | 1 | 1 | 1 | 1 | 0.08 | >jgi\|Phyca11\|9799\|fgenesh1_pm.PHYCAscaffold_42_#_7 |
| 95 | 2 | 116241 | 31 | 44437 | 2 | 2 | 2 | 2 | 0.21 | >jgi\|Phyca11\|116241\|e_gw1.30.536.1 |
| 275 | 1 | 575591 | 31 | 52834 | 1 | 1 | 1 | 1 | 0.08 | >jgi\|Phyca11\|575591\|estExt2_Genewise1.C_PHYCAscaffold_760055 |
| 276 | 1 | 21126 | 31 | 74492 | 1 | 1 | 1 | 1 | 0.06 | >jgi\|Phyca11\|21126\|fgenesh1_pg.PHYCAscaffold_82_#_34 |
| 277 | 1 | 560279 | 31 | 47305 | 1 | 1 | 1 | 1 | 0.09 | >jgi\|Phyca11\|560279\|estExt2_Genewise1.C_PHYCAscaffold_40823 |
| 278 | 1 | 569822 | 30 | 101155 | 1 | 1 | 1 | 1 | 0.04 | >jgi\|Phyca11\|569822\|estExt2_Genewise1.C_PHYCAscaffold_340150 |
| 279 | 1 | 503888 | 30 | 22688 | 1 | 1 | 1 | 1 | 0.2 | >jgi\|Phyca11\|503888\|fgenesh2_kg.PHYCAscaffold_5_#_52_#_Contig36.1 |
| 281 | 1 | 534256 | 30 | 49992 | 2 | 2 | 2 | 2 | 0.19 | >jgi\|Phyca11\|534256\|estExt2_fgenesh1_pg.C_PHYCAscaffold_210124 |
| 282 | 1 | 505362 | 30 | 15881 | 1 | 1 | 1 | 1 | 0.3 | >jgi\|Phyca11\|505362\|fgenesh2_kg.PHYCAscaffold_13_#_12_#_Contig171.1 |
| 283 | 1 | 504712 | 30 | 23021 | 1 | 1 | 1 | 1 | 0.2 | >jgi\|Phyca11\|504712\|fgenesh2_kg.PHYCAscaffold_9_#_103_#_Contig5242.1 |
| 285 | 1 | 505013 | 29 | 27838 | 2 | 2 | 2 | 2 | 0.35 | >jgi\|Phyca11\|505013\|fgenesh2_kg.PHYCAscaffold_10_#_173_#_gi\|189084885\|gb\|BT032416.1\| |
| 286 | 1 | 511514 | 29 | 6365 | 1 | 1 | 1 | 1 | 0.88 | >jgi\|Phyca11\|511514\|fgenesh2_kg.PHYCAscaffold_87_#_20_#_4097666:2 |
| 288 | 1 | 504114 | 29 | 98692 | 1 | 1 | 1 | 1 | 0.04 | >jgi\|Phyca11\|504114\|fgenesh2_kg.PHYCAscaffold_6_#_31_#_Contig2820.1 |
| 289 | 1 | 96835 | 29 | 50399 | 1 | 1 | 1 | 1 | 0.09 | >jgi\|Phyca11\|96835\|e_gw1.1.71.1 |
| 291 | 1 | 74521 | 29 | 64501 | 1 | 1 | 1 | 1 | 0.07 | >jgi\|Phyca11\|74521\|gw1.15.657.1 |
| 292 | 1 | 20863 | 29 | 71421 | 1 | 1 | 1 | 1 | 0.06 | >jgi\|Phyca11\|20863\|fgenesh1_pg.PHYCAscaffold_74_#_34 |
| 293 | 1 | 545078 | 29 | 60384 | 1 | 1 | 1 | 1 | 0.07 | >jgi\|Phyca11\|545078\|estExt2_Genewise1Plus.C_PHYCAscaffold_170080 |
| 294 | 1 | 508983 | 29 | 33906 | 2 | 2 | 2 | 2 | 0.28 | >jgi\|Phyca11\|508983\|fgenesh2_kg.PHYCAscaffold_40_#_72_#_Contig64.1 |
| 295 | 1 | 511304 | 29 | 48105 | 1 | 1 | 1 | 1 | 0.09 | >jgi\|Phyca11\|511304\|fgenesh2_kg.PHYCAscaffold_80_#_48_#_4098482:7 |
| 296 | 1 | 504590 | 28 | 23475 | 1 | 1 | 1 | 1 | 0.2 | >jgi\|Phyca11\|504590\|fgenesh2_kg.PHYCAscaffold_8_#_159_#_Contig101.1 |
| 297 | 1 | 102887 | 28 | 44261 | 1 | 1 | 1 | 1 | 0.1 | >jgi\|Phyca11\|102887\|e_gw1.7.1087.1 |
| 298 | 1 | 509308 | 28 | 35212 | 2 | 2 | 2 | 2 | 0.27 | >jgi\|Phyca11\|509308\|fgenesh2_kg.PHYCAscaffold_43_#_106_#_Contig105.1 |
| 299 | 1 | 506325 | 28 | 15185 | 1 | 1 | 1 | 1 | 0.32 | >jgi\|Phyca11\|506325\|fgenesh2_kg.PHYCAscaffold_19_#_31_#_4100643:1 |
| 300 | 1 | 545106 | 28 | 105339 | 1 | 1 | 1 | 1 | 0.04 | >jgi\|Phyca11\|545106\|estExt2_Genewise1Plus.C_PHYCAscaffold_170111 |
| 301 | 1 | 507961 | 28 | 82552 | 1 | 1 | 1 | 1 | 0.05 | >jgi\|Phyca11\|507961\|fgenesh2_kg.PHYCAscaffold_31_#_42_#_gi\|189092330\|gb\|BT032284.1\| |
| 302 | 1 | 508721 | 28 | 90477 | 1 | 1 | 1 | 1 | 0.05 | >jgi\|Phyca11\|508721\|fgenesh2_kg.PHYCAscaffold_37_#_96_#_4098316:1 |
| 303 | 1 | 527718 | 28 | 112622 | 1 | 1 | 1 | 1 | 0.04 | >jgi\|Phyca11\|527718\|estExt2_fgenesh1_pm.C_PHYCAscaffold_220027 |
| 305 | 1 | 505330 | 28 | 10735 | 1 | 1 | 1 | 1 | 0.47 | >jgi\|Phyca11\|505330\|fgenesh2_kg.PHYCAscaffold_12_#_118_#_4101945:268 |
| 306 | 1 | 509713 | 28 | 7861 | 1 | 1 | 1 | 1 | 0.67 | >jgi\|Phyca11\|509713\|fgenesh2_kg.PHYCAscaffold_49_#_5_#_gi\|189084943\|gb\|BT032475.1\| |
| 307 | 1 | 506552 | 28 | 30387 | 1 | 1 | 1 | 1 | 0.15 | >jgi\|Phyca11\|506552\|fgenesh2_kg.PHYCAscaffold_20_#_98_#_Contig1804.1 |
| 309 | 1 | 546414 | 27 | 83340 | 2 | 2 | 2 | 2 | 0.11 | >jgi\|Phyca11\|546414\|estExt2_Genewise1Plus.C_PHYCAscaffold_210088 |
| 310 | 1 | 109739 | 27 | 70567 | 1 | 1 | 1 | 1 | 0.06 | >jgi\|Phyca11\|109739\|e_gw1.17.194.1 |
| 311 | 1 | 33678 | 27 | 186894 | 1 | 1 | 1 | 1 | 0.02 | >jgi\|Phyca11\|33678\|gw1.11.1.1 |
| 312 | 1 | 128603 | 27 | 62433 | 1 | 1 | 1 | 1 | 0.07 | >jgi\|Phyca11\|128603\|e_gw1.77.203.1 |
| 313 | 1 | 532207 | 27 | 123366 | 1 | 1 | 1 | 1 | 0.04 | >jgi\|Phyca11\|532207\|estExt2_fgenesh1_pg.C_PHYCAscaffold_40153 |
| 314 | 1 | 570824 | 27 | 52801 | 1 | 1 | 1 | 1 | 0.08 | >jgi\|Phyca11\|570824\|estExt2_Genewise1.C_PHYCAscaffold_390124 |
| 315 | 1 | 15710 | 27 | 149674 | 1 | 1 | 1 | 1 | 0.03 | >jgi\|Phyca11\|15710\|fgenesh1_pg.PHYCAscaffold_15_#_48 |
| 316 | 1 | 511433 | 27 | 77859 | 2 | 2 | 2 | 2 | 0.12 | >jgi\|Phyca11\|511433\|fgenesh2_kg.PHYCAscaffold_85_#_1_#_gi\|189084627\|gb\|BT032143.1\| |
| 317 | 1 | 533830 | 27 | 47221 | 1 | 1 | 1 | 1 | 0.09 | >jgi\|Phyca11\|533830\|estExt2_fgenesh1_pg.C_PHYCAscaffold_170166 |
| 318 | 1 | 553484 | 26 | 89673 | 1 | 1 | 1 | 1 | 0.05 | >jgi\|Phyca11\|553484\|estExt2_Genewise1Plus.C_PHYCAscaffold_530198 |
| 319 | 1 | 540044 | 26 | 134595 | 1 | 1 | 1 | 1 | 0.03 | >jgi\|Phyca11\|540044\|estExt2_Genewise1Plus.C_PHYCAscaffold_40375 |
| 320 | 1 | 534485 | 26 | 47695 | 2 | 2 | 2 | 2 | 0.19 | >jgi\|Phyca11\|534485\|estExt2_fgenesh1_pg.C_PHYCAscaffold_240045 |
| 321 | 1 | 527499 | 26 | 204833 | 2 | 2 | 2 | 2 | 0.04 | >jgi\|Phyca11\|527499\|estExt2_fgenesh1_pm.C_PHYCAscaffold_200021 |
| 322 | 1 | 504276 | 26 | 76273 | 1 | 1 | 1 | 1 | 0.06 | >jgi\|Phyca11\|504276\|fgenesh2_kg.PHYCAscaffold_7_#_28_#_Contig229.1 |
| 323 | 1 | 568556 | 26 | 82124 | 1 | 1 | 1 | 1 | 0.05 | >jgi\|Phyca11\|568556\|estExt2_Genewise1.C_PHYCAscaffold_290239 |
| 324 | 1 | 527368 | 26 | 51532 | 1 | 1 | 1 | 1 | 0.09 | >jgi\|Phyca11\|527368\|estExt2_fgenesh1_pm.C_PHYCAscaffold_180094 |
| 325 | 1 | 505958 | 26 | 37035 | 1 | 1 | 1 | 1 | 0.12 | >jgi\|Phyca11\|505958\|fgenesh2_kg.PHYCAscaffold_17_#_43_#_4096633:2 |
| 326 | 1 | 6335 | 26 | 100909 | 2 | 2 | 2 | 2 | 0.09 | >jgi\|Phyca11\|6335\|fgenesh1_pm.PHYCAscaffold_11_#_51 |
| 327 | 1 | 541134 | 26 | 68189 | 1 | 1 | 1 | 1 | 0.06 | >jgi\|Phyca11\|541134\|estExt2_Genewise1Plus.C_PHYCAscaffold_60201 |
| 328 | 1 | 510755 | 26 | 99322 | 1 | 1 | 1 | 1 | 0.04 | >jgi\|Phyca11\|510755\|fgenesh2_kg.PHYCAscaffold_66_#_29_#_Contig180.1 |
| 329 | 1 | 505505 | 26 | 36287 | 2 | 2 | 2 | 2 | 0.26 | >jgi\|Phyca11\|505505\|fgenesh2_kg.PHYCAscaffold_13_#_155_#_4100209:1 |
| 330 | 1 | 13219 | 26 | 57130 | 2 | 2 | 1 | 1 | 0.08 | >jgi\|Phyca11\|13219\|fgenesh1_pg.PHYCAscaffold_3_#_32 |
| 331 | 1 | 503988 | 25 | 25954 | 1 | 1 | 1 | 1 | 0.18 | >jgi\|Phyca11\|503988\|fgenesh2_kg.PHYCAscaffold_5_#_152_#_Contig1373.1 |
| 332 | 1 | 573771 | 25 | 91543 | 1 | 1 | 1 | 1 | 0.05 | >jgi\|Phyca11\|573771\|estExt2_Genewise1.C_PHYCAscaffold_550212 |
| 333 | 1 | 129924 | 25 | 37470 | 2 | 2 | 2 | 2 | 0.25 | >jgi\|Phyca11\|129924\|e_gw1.89.53.1 |
| 335 | 1 | 7562 | 25 | 142125 | 1 | 1 | 1 | 1 | 0.03 | >jgi\|Phyca11\|7562\|fgenesh1_pm.PHYCAscaffold_20_#_77 |
| 336 | 1 | 503485 | 25 | 22210 | 1 | 1 | 1 | 1 | 0.21 | >jgi\|Phyca11\|503485\|fgenesh2_kg.PHYCAscaffold_4_#_24_#_Contig5923.1 |
| 337 | 1 | 20237 | 25 | 108201 | 1 | 1 | 1 | 1 | 0.04 | >jgi\|Phyca11\|20237\|fgenesh1_pg.PHYCAscaffold_60_#_23 |
| 338 | 1 | 8252 | 24 | 49212 | 3 | 3 | 3 | 3 | 0.3 | >jgi\|Phyca11\|8252\|fgenesh1_pm.PHYCAscaffold_26_#_103 |
| 339 | 1 | 10181 | 24 | 205349 | 1 | 1 | 1 | 1 | 0.02 | >jgi\|Phyca11\|10181\|fgenesh1_pm.PHYCAscaffold_47_#_10 |
| 340 | 1 | 126828 | 24 | 15906 | 1 | 1 | 1 | 1 | 0.3 | >jgi\|Phyca11\|126828\|e_gw1.65.227.1 |
| 341 | 1 | 545398 | 24 | 123752 | 1 | 1 | 1 | 1 | 0.04 | >jgi\|Phyca11\|545398\|estExt2_Genewise1Plus.C_PHYCAscaffold_170580 |
| 342 | 1 | 571712 | 24 | 49744 | 1 | 1 | 1 | 1 | 0.09 | >jgi\|Phyca11\|571712\|estExt2_Genewise1.C_PHYCAscaffold_430233 |
| 343 | 1 | 561387 | 24 | 44291 | 1 | 1 | 1 | 1 | 0.1 | >jgi\|Phyca11\|561387\|estExt2_Genewise1.C_PHYCAscaffold_60808 |
| 344 | 1 | 8949 | 24 | 85750 | 1 | 1 | 1 | 1 | 0.05 | >jgi\|Phyca11\|8949\|fgenesh1_pm.PHYCAscaffold_32_#_84 |
| 345 | 1 | 6476 | 24 | 130861 | 2 | 2 | 2 | 2 | 0.07 | >jgi\|Phyca11\|6476\|fgenesh1_pm.PHYCAscaffold_12_#_56 |
| 346 | 1 | 502681 | 23 | 34485 | 1 | 1 | 1 | 1 | 0.13 | >jgi\|Phyca11\|502681\|fgenesh2_kg.PHYCAscaffold_1_#_141_#_4097655:1 |
| 347 | 1 | 99228 | 23 | 29295 | 1 | 1 | 1 | 1 | 0.16 | >jgi\|Phyca11\|99228\|e_gw1.3.1145.1 |
| 348 | 1 | 13207 | 23 | 35953 | 1 | 1 | 1 | 1 | 0.13 | >jgi\|Phyca11\|13207\|fgenesh1_pg.PHYCAscaffold_3_#_20 |
| 349 | 1 | 6432 | 23 | 251898 | 2 | 2 | 2 | 2 | 0.03 | >jgi\|Phyca11\|6432\|fgenesh1_pm.PHYCAscaffold_12_#_12 |
| 351 | 1 | 105275 | 23 | 34215 | 1 | 1 | 1 | 1 | 0.13 | >jgi\|Phyca11\|105275\|e_gw1.10.326.1 |
| 352 | 1 | 16541 | 23 | 175762 | 1 | 1 | 1 | 1 | 0.02 | >jgi\|Phyca11\|16541\|fgenesh1_pg.PHYCAscaffold_20_#_129 |
| 354 | 1 | 103907 | 23 | 39624 | 1 | 1 | 1 | 1 | 0.11 | >jgi\|Phyca11\|103907\|e_gw1.8.398.1 |
| 355 | 1 | 506416 | 23 | 98155 | 1 | 1 | 1 | 1 | 0.04 | >jgi\|Phyca11\|506416\|fgenesh2_kg.PHYCAscaffold_19_#_122_#_4102003:2 |
| 356 | 1 | 527885 | 23 | 92705 | 1 | 1 | 1 | 1 | 0.05 | >jgi\|Phyca11\|527885\|estExt2_fgenesh1_pm.C_PHYCAscaffold_230116 |
| 357 | 1 | 528255 | 23 | 94441 | 1 | 1 | 1 | 1 | 0.05 | >jgi\|Phyca11\|528255\|estExt2_fgenesh1_pm.C_PHYCAscaffold_280010 |
| 358 | 1 | 508067 | 23 | 24183 | 2 | 2 | 2 | 2 | 0.42 | >jgi\|Phyca11\|508067\|fgenesh2_kg.PHYCAscaffold_32_#_26_#_Contig1225.1 |
| 359 | 1 | 7637 | 22 | 214844 | 1 | 1 | 1 | 1 | 0.02 | >jgi\|Phyca11\|7637\|fgenesh1_pm.PHYCAscaffold_21_#_10 |
| 360 | 1 | 504366 | 22 | 39976 | 1 | 1 | 1 | 1 | 0.11 | >jgi\|Phyca11\|504366\|fgenesh2_kg.PHYCAscaffold_7_#_118_#_Contig309.1 |
| 361 | 1 | 109320 | 22 | 41148 | 1 | 1 | 1 | 1 | 0.11 | >jgi\|Phyca11\|109320\|e_gw1.16.118.1 |
| 362 | 1 | 503926 | 22 | 17112 | 1 | 1 | 1 | 1 | 0.28 | >jgi\|Phyca11\|503926\|fgenesh2_kg.PHYCAscaffold_5_#_90_#_4097723:1 |
| 363 | 1 | 113812 | 22 | 20248 | 1 | 1 | 1 | 1 | 0.23 | >jgi\|Phyca11\|113812\|e_gw1.25.423.1 |
| 364 | 1 | 97828 | 22 | 32060 | 1 | 1 | 1 | 1 | 0.14 | >jgi\|Phyca11\|97828\|e_gw1.2.922.1 |
| 365 | 1 | 506791 | 22 | 115416 | 1 | 1 | 1 | 1 | 0.04 | >jgi\|Phyca11\|506791\|fgenesh2_kg.PHYCAscaffold_22_#_30_#_gi\|189084047\|gb\|BT031563.1\| |
| 366 | 1 | 506572 | 22 | 14119 | 1 | 1 | 1 | 1 | 0.34 | >jgi\|Phyca11\|506572\|fgenesh2_kg.PHYCAscaffold_20_#_118_#_gi\|189083878\|gb\|BT031394.1\| |
| 367 | 1 | 541822 | 22 | 27426 | 1 | 1 | 1 | 1 | 0.17 | >jgi\|Phyca11\|541822\|estExt2_Genewise1Plus.C_PHYCAscaffold_70758 |
| 368 | 1 | 504252 | 22 | 41927 | 1 | 1 | 1 | 1 | 0.11 | >jgi\|Phyca11\|504252\|fgenesh2_kg.PHYCAscaffold_7_#_4_#_Contig922.1 |
| 369 | 1 | 508077 | 22 | 152340 | 1 | 1 | 1 | 1 | 0.03 | >jgi\|Phyca11\|508077\|fgenesh2_kg.PHYCAscaffold_32_#_36_#_Contig2846.1 |
| 370 | 1 | 122536 | 22 | 85692 | 1 | 1 | 1 | 1 | 0.05 | >jgi\|Phyca11\|122536\|e_gw1.48.64.1 |
| 371 | 1 | 507766 | 22 | 55815 | 1 | 1 | 1 | 1 | 0.08 | >jgi\|Phyca11\|507766\|fgenesh2_kg.PHYCAscaffold_30_#_35_#_Contig256.1 |
| 372 | 1 | 129462 | 22 | 204802 | 2 | 2 | 2 | 2 | 0.04 | >jgi\|Phyca11\|129462\|e_gw1.84.2.1 |
| 373 | 1 | 566774 | 21 | 51922 | 1 | 1 | 1 | 1 | 0.09 | >jgi\|Phyca11\|566774\|estExt2_Genewise1.C_PHYCAscaffold_220429 |
| 374 | 1 | 75556 | 21 | 49881 | 1 | 1 | 1 | 1 | 0.09 | >jgi\|Phyca11\|75556\|gw1.21.649.1 |
| 375 | 1 | 542873 | 21 | 55889 | 1 | 1 | 1 | 1 | 0.08 | >jgi\|Phyca11\|542873\|estExt2_Genewise1Plus.C_PHYCAscaffold_100329 |
| 376 | 1 | 511117 | 21 | 18412 | 1 | 1 | 1 | 1 | 0.26 | >jgi\|Phyca11\|511117\|fgenesh2_kg.PHYCAscaffold_76_#_18_#_Contig2045.1 |
| 377 | 1 | 55483 | 21 | 63394 | 1 | 1 | 1 | 1 | 0.07 | >jgi\|Phyca11\|55483\|gw1.18.166.1 |
| 378 | 1 | 39322 | 21 | 29698 | 1 | 1 | 1 | 1 | 0.15 | >jgi\|Phyca11\|39322\|gw1.83.18.1 |
| 379 | 1 | 506779 | 21 | 50948 | 1 | 1 | 1 | 1 | 0.09 | >jgi\|Phyca11\|506779\|fgenesh2_kg.PHYCAscaffold_22_#_18_#_Contig4797.1 |
| 380 | 1 | 511445 | 21 | 89242 | 1 | 1 | 1 | 1 | 0.05 | >jgi\|Phyca11\|511445\|fgenesh2_kg.PHYCAscaffold_85_#_13_#_Contig2917.1 |
| 381 | 1 | 576261 | 21 | 114090 | 1 | 1 | 1 | 1 | 0.04 | >jgi\|Phyca11\|576261\|estExt2_Genewise1.C_PHYCAscaffold_860021 |
| 382 | 1 | 506946 | 20 | 62348 | 1 | 1 | 1 | 1 | 0.07 | >jgi\|Phyca11\|506946\|fgenesh2_kg.PHYCAscaffold_23_#_77_#_Contig155.1 |
| 383 | 1 | 507525 | 20 | 56350 | 1 | 1 | 1 | 1 | 0.08 | >jgi\|Phyca11\|507525\|fgenesh2_kg.PHYCAscaffold_28_#_32_#_Contig2244.1 |
| 384 | 1 | 510359 | 20 | 70048 | 1 | 1 | 1 | 1 | 0.06 | >jgi\|Phyca11\|510359\|fgenesh2_kg.PHYCAscaffold_58_#_72_#_Contig2242.1 |
| 385 | 1 | 577035 | 20 | 20283 | 1 | 1 | 1 | 1 | 0.23 | >jgi\|Phyca11\|577035\|estExt2_Genewise1.C_PHYCAscaffold_1000103 |
| 386 | 1 | 574588 | 20 | 146603 | 1 | 1 | 1 | 1 | 0.03 | >jgi\|Phyca11\|574588\|estExt2_Genewise1.C_PHYCAscaffold_640065 |
| 387 | 1 | 561666 | 20 | 29051 | 1 | 1 | 1 | 1 | 0.16 | >jgi\|Phyca11\|561666\|estExt2_Genewise1.C_PHYCAscaffold_70606 |
| 388 | 1 | 504259 | 20 | 129675 | 1 | 1 | 1 | 1 | 0.03 | >jgi\|Phyca11\|504259\|fgenesh2_kg.PHYCAscaffold_7_#_11_#_Contig1696.1 |
| 389 | 1 | 547116 | 20 | 42183 | 1 | 1 | 1 | 1 | 0.11 | >jgi\|Phyca11\|547116\|estExt2_Genewise1Plus.C_PHYCAscaffold_230401 |
| 390 | 1 | 545267 | 19 | 73776 | 1 | 1 | 1 | 1 | 0.06 | >jgi\|Phyca11\|545267\|estExt2_Genewise1Plus.C_PHYCAscaffold_170368 |
| 391 | 1 | 4484 | 19 | 164721 | 1 | 1 | 1 | 1 | 0.03 | >jgi\|Phyca11\|4484\|fgenesh1_pm.PHYCAscaffold_2_#_80 |
| 392 | 1 | 527927 | 19 | 74101 | 1 | 1 | 1 | 1 | 0.06 | >jgi\|Phyca11\|527927\|estExt2_fgenesh1_pm.C_PHYCAscaffold_240054 |
| 393 | 1 | 525762 | 19 | 117149 | 1 | 1 | 1 | 1 | 0.04 | >jgi\|Phyca11\|525762\|estExt2_fgenesh1_pm.C_PHYCAscaffold_50055 |
| 394 | 1 | 570139 | 19 | 57657 | 1 | 1 | 1 | 1 | 0.08 | >jgi\|Phyca11\|570139\|estExt2_Genewise1.C_PHYCAscaffold_350336 |
| 395 | 1 | 541200 | 19 | 15076 | 1 | 1 | 1 | 1 | 0.32 | >jgi\|Phyca11\|541200\|estExt2_Genewise1Plus.C_PHYCAscaffold_60305 |
| 396 | 1 | 577453 | 19 | 46059 | 1 | 1 | 1 | 1 | 0.1 | >jgi\|Phyca11\|577453\|estExt2_Genewise1.C_PHYCAscaffold_1190011 |
| 397 | 1 | 504915 | 19 | 52560 | 1 | 1 | 1 | 1 | 0.08 | >jgi\|Phyca11\|504915\|fgenesh2_kg.PHYCAscaffold_10_#_75_#_4096762:2 |
| 398 | 1 | 507091 | 19 | 50214 | 1 | 1 | 1 | 1 | 0.09 | >jgi\|Phyca11\|507091\|fgenesh2_kg.PHYCAscaffold_24_#_83_#_4100347:2 |
| 28 | 11 | 509735 | 18 | 28698 | 2 | 2 | 2 | 2 | 0.34 | >jgi\|Phyca11\|509735\|fgenesh2_kg.PHYCAscaffold_49_#_27_#_4097066:1 |
| 399 | 1 | 505546 | 18 | 24844 | 1 | 1 | 1 | 1 | 0.18 | >jgi\|Phyca11\|505546\|fgenesh2_kg.PHYCAscaffold_14_#_27_#_Contig1965.1 |
| 400 | 1 | 532971 | 18 | 57903 | 1 | 1 | 1 | 1 | 0.08 | >jgi\|Phyca11\|532971\|estExt2_fgenesh1_pg.C_PHYCAscaffold_90155 |
| 401 | 1 | 577365 | 18 | 67399 | 1 | 1 | 1 | 1 | 0.07 | >jgi\|Phyca11\|577365\|estExt2_Genewise1.C_PHYCAscaffold_1090022 |
| 402 | 1 | 511466 | 18 | 78712 | 1 | 1 | 1 | 1 | 0.06 | >jgi\|Phyca11\|511466\|fgenesh2_kg.PHYCAscaffold_86_#_3_#_gi\|189084813\|gb\|BT032344.1\| |
| 404 | 1 | 510315 | 18 | 30905 | 1 | 1 | 1 | 1 | 0.15 | >jgi\|Phyca11\|510315\|fgenesh2_kg.PHYCAscaffold_58_#_28_#_Contig4113.1 |
| 405 | 1 | 549506 | 18 | 69618 | 1 | 1 | 1 | 1 | 0.06 | >jgi\|Phyca11\|549506\|estExt2_Genewise1Plus.C_PHYCAscaffold_330019 |
| 406 | 1 | 123621 | 18 | 52506 | 1 | 1 | 1 | 1 | 0.08 | >jgi\|Phyca11\|123621\|e_gw1.51.113.1 |
| 407 | 1 | 83443 | 18 | 7058 | 1 | 1 | 1 | 1 | 0.77 | >jgi\|Phyca11\|83443\|gw1.30.562.1 |
| 408 | 1 | 127727 | 18 | 113405 | 2 | 2 | 1 | 1 | 0.04 | >jgi\|Phyca11\|127727\|e_gw1.71.37.1 |
| 409 | 1 | 503852 | 18 | 17433 | 1 | 1 | 1 | 1 | 0.27 | >jgi\|Phyca11\|503852\|fgenesh2_kg.PHYCAscaffold_5_#_16_#_4101529:5 |
| 410 | 1 | 9440 | 18 | 99244 | 1 | 1 | 1 | 1 | 0.04 | >jgi\|Phyca11\|9440\|fgenesh1_pm.PHYCAscaffold_37_#_91 |
| 411 | 1 | 114819 | 18 | 35115 | 1 | 1 | 1 | 1 | 0.13 | >jgi\|Phyca11\|114819\|e_gw1.27.360.1 |
| 412 | 1 | 508791 | 18 | 98965 | 1 | 1 | 1 | 1 | 0.04 | >jgi\|Phyca11\|508791\|fgenesh2_kg.PHYCAscaffold_38_#_43_#_Contig2859.1 |
| 413 | 1 | 527220 | 18 | 70992 | 1 | 1 | 1 | 1 | 0.06 | >jgi\|Phyca11\|527220\|estExt2_fgenesh1_pm.C_PHYCAscaffold_170102 |
| 414 | 1 | 503312 | 17 | 19172 | 1 | 1 | 1 | 1 | 0.24 | >jgi\|Phyca11\|503312\|fgenesh2_kg.PHYCAscaffold_3_#_177_#_4097869:1 |
| 415 | 1 | 129692 | 17 | 62217 | 1 | 1 | 1 | 1 | 0.07 | >jgi\|Phyca11\|129692\|e_gw1.86.45.1 |
| 416 | 1 | 505595 | 17 | 37961 | 1 | 1 | 1 | 1 | 0.12 | >jgi\|Phyca11\|505595\|fgenesh2_kg.PHYCAscaffold_14_#_76_#_Contig4404.1 |
| 417 | 1 | 507030 | 17 | 41223 | 1 | 1 | 1 | 1 | 0.11 | >jgi\|Phyca11\|507030\|fgenesh2_kg.PHYCAscaffold_24_#_22_#_Contig274.1 |
| 418 | 1 | 573749 | 17 | 86552 | 1 | 1 | 1 | 1 | 0.05 | >jgi\|Phyca11\|573749\|estExt2_Genewise1.C_PHYCAscaffold_550157 |
| 419 | 1 | 511317 | 17 | 76318 | 1 | 1 | 1 | 1 | 0.06 | >jgi\|Phyca11\|511317\|fgenesh2_kg.PHYCAscaffold_80_#_61_#_gi\|189084555\|gb\|BT032071.1\| |
| 420 | 1 | 568375 | 17 | 60822 | 1 | 1 | 1 | 1 | 0.07 | >jgi\|Phyca11\|568375\|estExt2_Genewise1.C_PHYCAscaffold_280427 |
| 421 | 1 | 547393 | 17 | 29066 | 1 | 1 | 1 | 1 | 0.16 | >jgi\|Phyca11\|547393\|estExt2_Genewise1Plus.C_PHYCAscaffold_240415 |
| 422 | 1 | 506190 | 17 | 20826 | 1 | 1 | 1 | 1 | 0.22 | >jgi\|Phyca11\|506190\|fgenesh2_kg.PHYCAscaffold_18_#_63_#_gi\|189084437\|gb\|BT031953.1\| |
| 423 | 1 | 128908 | 17 | 31834 | 1 | 1 | 1 | 1 | 0.14 | >jgi\|Phyca11\|128908\|e_gw1.79.95.1 |
| 424 | 1 | 510248 | 17 | 67031 | 1 | 1 | 1 | 1 | 0.07 | >jgi\|Phyca11\|510248\|fgenesh2_kg.PHYCAscaffold_55_#_65_#_gi\|189084581\|gb\|BT032097.1\| |
| 425 | 1 | 535142 | 17 | 56284 | 1 | 1 | 1 | 1 | 0.08 | >jgi\|Phyca11\|535142\|estExt2_fgenesh1_pg.C_PHYCAscaffold_320051 |
| 426 | 1 | 573119 | 17 | 52311 | 1 | 1 | 1 | 1 | 0.08 | >jgi\|Phyca11\|573119\|estExt2_Genewise1.C_PHYCAscaffold_510194 |
| 427 | 1 | 569841 | 16 | 15587 | 1 | 1 | 1 | 1 | 0.31 | >jgi\|Phyca11\|569841\|estExt2_Genewise1.C_PHYCAscaffold_340180 |
| 428 | 1 | 119154 | 16 | 53093 | 1 | 1 | 1 | 1 | 0.08 | >jgi\|Phyca11\|119154\|e_gw1.37.173.1 |
| 429 | 1 | 503860 | 16 | 115248 | 2 | 2 | 2 | 2 | 0.08 | >jgi\|Phyca11\|503860\|fgenesh2_kg.PHYCAscaffold_5_#_24_#_gi\|189084064\|gb\|BT031580.1\| |
| 430 | 1 | 19832 | 16 | 37148 | 1 | 1 | 1 | 1 | 0.12 | >jgi\|Phyca11\|19832\|fgenesh1_pg.PHYCAscaffold_52_#_84 |
| 431 | 1 | 524980 | 16 | 63621 | 2 | 2 | 2 | 2 | 0.14 | >jgi\|Phyca11\|524980\|estExt2_fgenesh1_pm.C_PHYCAscaffold_10205 |
| 432 | 1 | 16379 | 16 | 16484 | 1 | 1 | 1 | 1 | 0.29 | >jgi\|Phyca11\|16379\|fgenesh1_pg.PHYCAscaffold_19_#_126 |
| 433 | 1 | 20438 | 16 | 80714 | 1 | 1 | 1 | 1 | 0.05 | >jgi\|Phyca11\|20438\|fgenesh1_pg.PHYCAscaffold_64_#_24 |
| 434 | 1 | 570034 | 16 | 115203 | 1 | 1 | 1 | 1 | 0.04 | >jgi\|Phyca11\|570034\|estExt2_Genewise1.C_PHYCAscaffold_350131 |
| 435 | 1 | 565873 | 16 | 64634 | 1 | 1 | 1 | 1 | 0.07 | >jgi\|Phyca11\|565873\|estExt2_Genewise1.C_PHYCAscaffold_190295 |
| 436 | 1 | 98941 | 16 | 18397 | 1 | 1 | 1 | 1 | 0.26 | >jgi\|Phyca11\|98941\|e_gw1.3.1180.1 |
| 437 | 1 | 506604 | 16 | 136490 | 1 | 1 | 1 | 1 | 0.03 | >jgi\|Phyca11\|506604\|fgenesh2_kg.PHYCAscaffold_20_#_150_#_Contig1038.1 |
| 438 | 1 | 502973 | 16 | 22064 | 1 | 1 | 1 | 1 | 0.21 | >jgi\|Phyca11\|502973\|fgenesh2_kg.PHYCAscaffold_2_#_89_#_4101945:287 |
| 439 | 1 | 124731 | 15 | 29450 | 1 | 1 | 1 | 1 | 0.15 | >jgi\|Phyca11\|124731\|e_gw1.54.272.1 |
| 440 | 1 | 529913 | 15 | 57909 | 1 | 1 | 1 | 1 | 0.08 | >jgi\|Phyca11\|529913\|estExt2_fgenesh1_pm.C_PHYCAscaffold_510016 |
| 441 | 1 | 571966 | 15 | 97924 | 1 | 1 | 1 | 1 | 0.04 | >jgi\|Phyca11\|571966\|estExt2_Genewise1.C_PHYCAscaffold_460011 |
| 442 | 1 | 511938 | 15 | 46144 | 1 | 1 | 1 | 1 | 0.1 | >jgi\|Phyca11\|511938\|fgenesh2_kg.PHYCAscaffold_105_#_36_#_4097069:3 |
| 443 | 1 | 508267 | 15 | 138934 | 1 | 1 | 1 | 1 | 0.03 | >jgi\|Phyca11\|508267\|fgenesh2_kg.PHYCAscaffold_33_#_126_#_Contig952.1 |
| 444 | 1 | 575295 | 15 | 86674 | 1 | 1 | 1 | 1 | 0.05 | >jgi\|Phyca11\|575295\|estExt2_Genewise1.C_PHYCAscaffold_720073 |
| 445 | 1 | 106531 | 15 | 23657 | 1 | 1 | 1 | 1 | 0.19 | >jgi\|Phyca11\|106531\|e_gw1.12.838.1 |
| 446 | 1 | 14250 | 15 | 112630 | 1 | 1 | 1 | 1 | 0.04 | >jgi\|Phyca11\|14250\|fgenesh1_pg.PHYCAscaffold_6_#_172 |
| 447 | 1 | 503558 | 15 | 39601 | 1 | 1 | 1 | 1 | 0.11 | >jgi\|Phyca11\|503558\|fgenesh2_kg.PHYCAscaffold_4_#_97_#_Contig683.1 |
| 448 | 1 | 536809 | 15 | 20370 | 1 | 1 | 1 | 1 | 0.23 | >jgi\|Phyca11\|536809\|estExt2_fgenesh1_pg.C_PHYCAscaffold_640020 |
| 449 | 1 | 96553 | 15 | 49247 | 1 | 1 | 1 | 1 | 0.09 | >jgi\|Phyca11\|96553\|e_gw1.1.45.1 |
| 450 | 1 | 545230 | 14 | 92277 | 1 | 1 | 1 | 1 | 0.05 | >jgi\|Phyca11\|545230\|estExt2_Genewise1Plus.C_PHYCAscaffold_170321 |
| 451 | 1 | 104701 | 14 | 37214 | 1 | 1 | 1 | 1 | 0.12 | >jgi\|Phyca11\|104701\|e_gw1.9.628.1 |
| 452 | 1 | 525091 | 14 | 19610 | 1 | 1 | 1 | 1 | 0.24 | >jgi\|Phyca11\|525091\|estExt2_fgenesh1_pm.C_PHYCAscaffold_20022 |
| 453 | 1 | 503542 | 14 | 53948 | 1 | 1 | 1 | 1 | 0.08 | >jgi\|Phyca11\|503542\|fgenesh2_kg.PHYCAscaffold_4_#_81_#_Contig1894.1 |
| 454 | 1 | 19969 | 14 | 13600 | 1 | 1 | 1 | 1 | 0.36 | >jgi\|Phyca11\|19969\|fgenesh1_pg.PHYCAscaffold_54_#_66 |
| 455 | 1 | 113258 | 14 | 34927 | 1 | 1 | 1 | 1 | 0.13 | >jgi\|Phyca11\|113258\|e_gw1.23.56.1 |
